# Association of haplotypes of AAP family amino acid transporters with nitrogen response and nitrogen use efficiency in rice grown under hydroponics and field conditions

**DOI:** 10.1101/2025.08.12.669810

**Authors:** Haritha Si, Kanya Rai, Rakesh Pandey, Archana Sahani, Ranjith Ellur, Shailendra K Jha, Viswanathan Chinnusamy, Lekshmy Sathee

## Abstract

The low nitrogen use efficiency (NUE) of staple food crops like rice has both economic and environmental impact. Amino acids taken up from soil or synthesized in source tissues and transported to sink tissues influence plant development and NUE. Amino acid permeases (AAPs) mediate amino acid uptake and transport and influence plant development, yield and NUE. NUE is a complex trait critical for improving rice productivity under variable N supply. Amino acid permeases such as OsAAP3, OsAAP5, and OsAAP11 have been shown to function as negative regulators of NUE in Japonica rice, yet their roles in Indica rice remain to be fully elucidated. In this study, we evaluated selected Indica genotypes under contrasting N regimes—seedlings were tested under high nitrate (HN), high ammonium (HA), and low N (LN) conditions, and field performance was assessed under N120 (optimum N) and N0 (low N). Our aim was to relate performance differences to the SNP status in these three genes, thereby clarifying their influence on N uptake, assimilation, yield potential, and N stress adaptation. ARC 10799 had non-synonymous SNP in OsAAP3, OsAAP5 and OsAAP11. Rice accessions like “Local,” “NCS901,” and “Bhainsa Mundariya,” showed superior biomass accumulation relative to MTU1010. Interestingly, the trend observed in the NUE calculated in seedling stage was similar to those proven in field evaluation and this proves that in general, genotypes with non-synonymous mutation in AAP3, AAP5 and AAP11 in comparison to Japonica showed better growth and N content. Data suggests an N induced regulation of physiological parameters and the changes in NUE parameters was at least in part, related to variation in biomass, plant height, photosynthesis and pigment content related parameters. The comparatively dissimilar trends in shown by different AAP haplotypes indicates the possibility of high or low correlation trait-wise, offering an opportunity to identify significant contrasts in a divergent set of AAP haplotypes. The previous reports and current findings opens new avenues and insights for improving rice yield, quality and NUE by changes in AAP haplotypes that occur naturally or created by precise genome editing.

## Introduction

Rice is the staple diet for half of the world population and the Asia-Pacific region is the topmost producer and consumer of rice which ceaselessly adds about 51 million rice consumers annually (Papademetriou, 2000). Taking into consideration the estimated projection of the world population growth which is about to reach 8.6 billion by 2030 (UN, 2022) and the 17 Sustainable Development Goals (SDGs) which were adopted by the UN in 2015 to advance both human development and environmental preservation by 2030 (UN, 2015), it is necessary to enhance global rice production in a sustainable manner which is currently forecasted to reach 532.9 million tonnes in 2024/2025 (Childs and Jarrell, 2025). Nitrogen (N) is an indispensable element which is the most abundant macronutrient yet the most limiting factor for plant growth. Nitrogen is a part of all building blocks of life such as nucleic acids, amino acids, proteins and various metabolites (Ohyama, 2010). The application of N fertilizers since the 1960s drastically increased the yield in modern rice cultivars but the excess application has accelerated environmental pollution due to run-off into waterbodies (Wu et al., 2024). Cereals utilize 55% of global nitrogen (N) fertilizer (58 Tg N) (IFA, 2016) yet the recovery of N is low (30-50%) (Ladha et al., 2016). Nitrogen Use Efficiency (NUE) is a trait comprising of complex interactions, genetic and environmental effects (Xu et al., 2012) and improvement of NUE ought to be one of the main goals in rice breeding for the effective N uptake, N assimilation and N remobilization of the available N resources (Cho et al., 2007). High NUE rice cultivars bearing abundant productive tillers paves the way for sustainable increase of rice production along with proper N management (Wu et al., 2024). This would expedite the efforts to reach the SDGs such as zero hunger (SDG 2), good health and well-being (SDG 3), clean water and sanitation (SDG 6), climate action (SDG 13), life on land (SDG 15) and especially responsible consumption and production (SDG 12) (Ladha et al., 2020).

Amino acids are known to alter the shoot and root architecture, regulate flowering time, seed number, size and quality, plant defence to stress and also function as signal molecules. For instance, glutamine acts as a sensor of N status and is responsible for feedback regulation of NH_4_^+^ uptake in rice (Guo *et al*., 2021; Dinkeloo *et al*., 2018). Thus, amino acids which are transported to various source and sinks in plants contribute to NUE and influence the plant growth and development in N-rich and N-deficit environments (Perchlik et al., 2017). Several families of transporters have been involved in amino acid uptake and transport. These include the Amino Acid/Auxin Permease family (AAAP), the Amino acid-Polyamine-Organocation family (APC) and the Usually Multiple Amino Acids Move In and Out Transporter (UMAMIT). AAPs are transmembrane amino acid transporters (AAT) belonging to the Amino Acid/Auxin Permease family (AAAP) and play an indispensable role in loading of amino acids for N sink and supply (Tegeder and Ward, 2012). They show moderate affinity and broad specificity to substrates (Tegeder *et al*., 2018). AAP1/NAT2 (Amino Acid Permease 1/ Neutral Amino Acid Transport system 2) was the first AAP to be identified in *Arabidopsis* (Frommer *et al*., 1993; Hsu *et al*., 1993) and later many AAP family transporters have been identified in various plant species viz. 8 in *Arabidopsis* (Taylor *et al*., 2015), 19 in japonica rice (Tegeder and Ward, 2012), *Solanum tuberosum* (Koch *et al*., 2003), *Populus tremula* (Couturier *et al*., 2010), *Ricinus communis* (Fischer *et al*., 1998), *Nepenthes alata* (Schulze *et al*., 1999), legumes such as *Pisum sativum* (Tegeder *et al*., 2000) and *Vicia faba* (Miranda *et al*., 2001).

Haplotype is a combination of nucleotides or DNA markers present in polymorphic sites which are inherited together in the same chromosomal segment (Stephens et al., 2001; Lu et al., 2010). Although most of the genes which contribute to the major traits have been already identified, still the information regarding the haplotype combinations of key genes in rice is lacking and several haplotype analysis studies have been done on various NUE genes in rice such as *OsNPF6.1, DNR1, MYB61, SBM1, NGR2, OsNR2, NGR5, OsTCP1, ARE1, DEP1, OsNAC42, OsNLP4, NRT1.1B, TOND1* (Li et al., 2022). The haplotype studies have been done in various *OsAAPs* such as *OsAAP1*, *OsAAP3, OsAAP4, OsSAAP5, OsAAP6* in Indica and Japonica varieties. The Indica haplotype of OsAAP5 due to a 51 bp insertion in the promoter sequence produced lesser *OsAAP5* transcripts and produced more tillers than Japonica haplotypes (Wang et al., 2019). Haplotype analysis of *OsAAP1* in both indica and non indica subpopulations, revealed that Hap5 has a favourable allele for effective tiller number (Ren et al., 2021). 8 haplotypes divided into 2 subgroups were found in *OsAAP6* in which subgroup A showed improved grain protein content (Peng et al., 2014). The rice population has been divided into 25 *OsAAP3* and 5 *OsAAP4* haplotypes in which *OsAAP3* and *OsAAP4* showed negative and positive correlations with NUE and grain yield respectively (Lu et al., 2018, Fang et al., 2021). These studies provide various insights about the natural genetic variation which contributes to the diversity of rice growth and development, adaptation to stress conditions and can be potential targets to improve NUE and yield (Guo et al., 2021). In the present study, we undertook the identification haplotypes for the genes *OsAAP3, OsAAP5* and *OsAAP11* from IRRI 3K Rice Genome Panel using SNP Seek database and selected lines were evaluated for physiological in response to different N treatments in hydroponics and field.

## Materials and methods

### Identification of AAP haplotypes

Haplotype analysis of *OsAAP3, OsAAP5* and *OsAAP11* genes was carried out in SNP seek database (https://snpseek.irri.org/) (Mansueto *et al*., 2017). The reference genome used in this study is Nipponbare. Among the 3024 rice accessions present in the 3K Rice genome panel, 354 accessions were available which included the following subpopulation (21 admix, 21 aro, 64 aus, 5 japx, 14 subtrop,4 temp, 6 trop, 221 ind (5 ind1A, 6 ind1B, 111 ind2, 2 ind3, 97 indx), 13 *Oryza sativa* Indica accessions (4 indx, 8 ind2, 1 ind1B) were selected which had only non-synonymous SNPs in the following genes viz., *OsAAP3, OsAAP5, OsAAP11* and one accession (indx) was selected which had haplotype similar to the reference genome Japonica Nipponbare for the genes *OsAAP3, OsAAP5, OsAAP11*. From the 3K Rice Genome Panel of IRRI, 14 *Oryza sativa* Indica accessions were selected and grown along with *Oryza sativa* Indica c*v*MTU1010 rice accession **(Supplementary Table 1, Fig 1a)**.

**Fig. 1.**
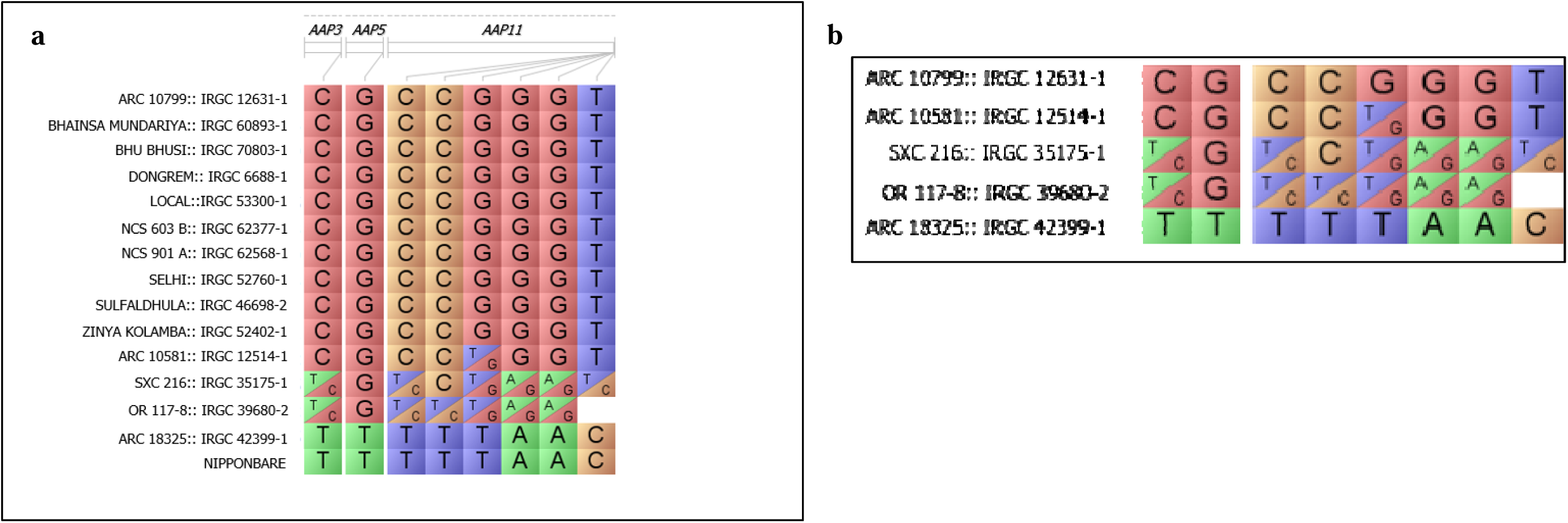
Genotypes selected for hydroponic evaluation and field (a) the haplotype details in comparison to Japonica cultivar Nipponbare is shown for the selected Indica genotypes having non-synonymous SNPs for *OsAAP3, OsAAP5* and *OsAAP11* (b) the haplotype details in comparison to Japonica cultivar Nipponbare is shown for the selected Indica genotypes for field evaluation. Data is retrieved from m the IRRI 3k rice panel SNP-SEEK database and viewed in Flapjack 1.22.04.21 software. ARC 18325:: IRGC 42399-1 has the same AAP haplotype as Nipponbare.

### Plant growth in hydroponics

The study was conducted at Division of Plant Physiology and National Phytotron Facility, ICAR-Indian Agricultural Research Institute, New Delhi during October 2023 – August 2024. The aim was to perform physiological evaluation of rice accessions comprising of the identified haplotypes along with popular rice cultivar, MTU1010 under High NO_3_^-^ (HN), High NH_4_^+^ (HA), Low N (LN) treatments in hydroponics under controlled environment conditions. The 14 *Oryza sativa indica* rice accessions along with MTU1010 were raised in hydroponics under controlled environment conditions up to seedling stage for 30 days in the National Phytotron facility, IARI, New Delhi **(Supplementary Table 1, Fig 1a**, **Fig 2)**. The seeds were rinsed with double distilled water and then surface sterilized with 2% sodium hypochlorite solution for 5 minutes and then thoroughly washed with double distilled water to remove the remaining traces of the sterilizing agent. The sterilized seeds were wrapped in a moist germination paper and then shifted to a beaker containing distilled water which was then kept in BOD incubator at 30°C for germination. After 6 days, the germinated seedlings were transplanted in the hydroponics system under three various treatments (HN: 7mM NO_3_^-^ and 0.5mM NH_4_^+^, HA: 7mM NH_4_^+^ and 0.5mM NO_3_^-^, LN: 0.24mM N). The seedlings were held with Styrofoam plugs in thermocol sheets kept in plastic tubs of 20 litre capacity. A total of 4 replicates and 12 seedlings for each treatment were divided into two trays with each tray containing 2 replicates with 3 plants in each following Completely Randomized design (CRD). The trays contained 20 litre of nutrient media solution of the composition as followed by (Jagadhesan et al., 2020) **(Fig. 2)**

**Fig 2.**
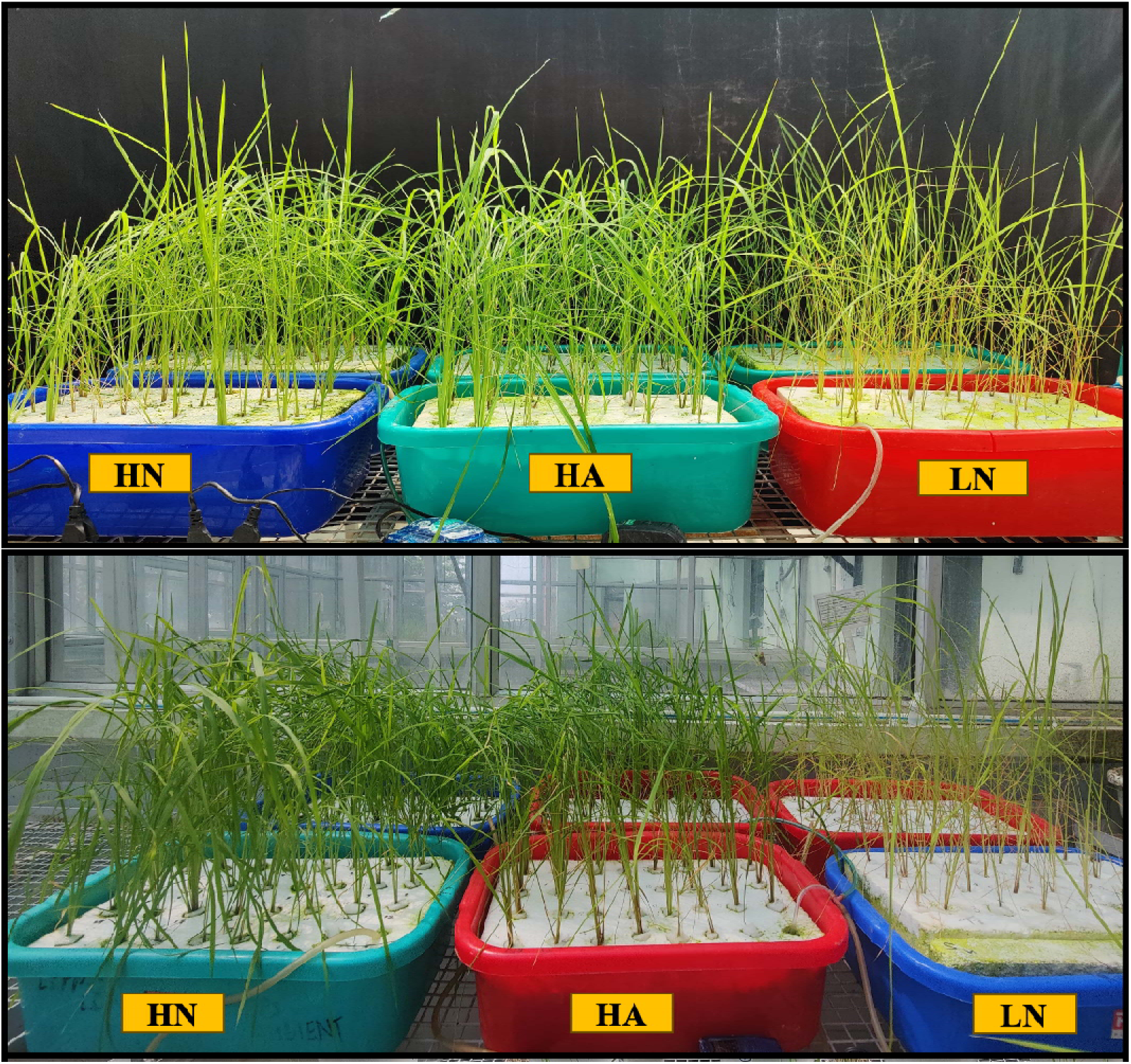
Representative images showing the growth of rice genotypes belonging to different AAP3, AAP5 and AAP11 haplotypes in hydroponics under three different N treatments such as high nitrate (HN), high ammonium (HA) and nitrogen deficient (LN) conditions (HN: 7mM NO_3_^-^ and 0.5mM NH_4_^+^, HA: 7mM NH_4_^+^ and 0.5mM NO_3_^-^, LN: 0.24mM N)

### Morphophysiological analysis of seedlings grown in hydroponics

Shoot fresh weight (SFW), shoot dry weight (SDW), plant height (PH) and leaf area (LA) (using LiCOR 3100, Lincon Nebraska, USA) were measured in 30 days old rice seedlings grown in hydroponics. The root system architecture measuring the following traits such as Total Root Length (TRL), Total Root Surface Area (TRSA), Total Root Volume, Length Of Lateral Root (Diameter≤0.5mm), Surface Area Of Lateral Root (LRSA), Volume Of Lateral Root (LRV), Length Of Main Root (Diameter>0.5mm) (MRL), Surface Area Of Main Root (Diameter>0.5mm) (MRSA), Volume Of Main Root (Diameter>0.5mm) (MRV), Root Diameter (RD), Root tips (Tips), Root forks (Forks) were done in root scanner (Win-RHIZO, Regent Instruments Inc.) and the images were saved in.TIF format which were later analyzed from WIN-Rhizo software. Chlorophyll content Index (CCI), was measured by Optical Science CCM-200 plus Chlorophyll Content Meter (CCM) by averaging the CCI of 10 leaf samples. Tissue NO_3_^-^ content was estimated using the method followed by Downes, 1978. The Total N content of the shoots was estimated using Kjeldahl’s method (Jones, 1988, Jagadhesan et al., 2020) and their NUE was calculated **(Supplementary** Figure 1a-b). Assay of N assimilation enzymes for the selected genotypes such as Nitrate reductase (NR, Nair and Abrol, 1976), Glutamine synthetase (GS), Glutamate synthase (GOGAT), Glutamate dehydrogenase (GDH) (Mohanty and Fletcher, 1980) and estimation of soluble protein content (Bradford, 1976) were performed.

### Field evaluation of rice genotypes for NUE and yield traits

The selected genotypes for field evaluation i.e. ARC 10799, ARC 18325, ARC 10581, SXC 216, OR 117-8 along with the high yielding check genotype MTU1010 were obtained from the Division of Genetics, ICAR-IARI (Fig 1b). The plants were raised during Kharif season (June-Oct 2024) under field conditions with recommended N, 120 Kg ha^−1^ (N^+^) and without N fertilizer application (N^-^) at Division of Plant Physiology, ICAR-IARI, New Delhi. The surface sterilized seeds were planted in nursery by broadcasting method and 25-30 days old seedlings were transplanted into the field with identification tags for each genotype **(Supplementary** Figure 2). Full dosage of P_2_O_5_ (as Single Super Phosphate) and K (as Muriate of Potash) was applied as a basal dose. 50% N was applied during field preparation before planting in N+ field, remaining 50% was applied as 2 split doses during early- and late-vegetative stages of the crop. The LI-6400XT PorTable Photosynthesis System, developed by LI-COR in Lincoln, Nebraska, USA, was used to measure various crucial plant physiological parameters such as photosynthetic rate (PN, µmol(CO_2_)m^-2^s^-1^), transpiration rate (TR, mmol(H_2_O)m^-2^s^-1^), stomatal conductance (gs, mol(H_2_O)m^-2^s^-1^), internal CO_2_ concentration (Ci, µmol(CO_2_)m^-2^s^-1^), Intrinsic Water Use Efficiency (iWUE, PN/gs) and Ci/Ca (Internal CO_2_ concentration (Ci)/ Ambient CO_2_ concentration (Ca)). Chlorophyll content Index (CCI), was measured by Optical Science CCM-200 plus Chlorophyll Content Meter (CCM) by averaging the CCI of 10 leaf samples. The thermal images of the crop canopy were taken using a porTable thermal infrared camera Fluke TiX620 model and the obtained IR thermal images (.IRB files) were then analyzed using SmartView 4.3 software and average of 100 points in the crop canopy was taken to measure the canopy temperature (CT) **(Supplementary** Figure 3**, 4)**. The plants were harvested at physiological maturity, oven dried and biomass traits such as Total fresh weight (TFW), total dry weight (TDW), yield traits such as tiller weight (TW), tiller no., panicle weight (PW), panicle no. were recorded. Plant material was then powdered and N content of shoot were estimated following Kjeldahl’s method (Jones, 1988). The total N content or uptake (g plant^-1^) was calculated by multiplying the N concentration (%) obtained from the Kjeldahl’s method with the biomass (g plant-1) of the plant (Baiyin et al., 2021). The NUE of a plant was determined by dividing the total biomass of the plant by its total N uptake and is expressed as g dry weight g^-1^ N uptake (Baiyin *et al*., 2021). The N Utilization Efficiency (NUtE) of the plant was estimated by dividing the grain weight per plant by its total N uptake and is expressed as g dry weight g^-1^ N uptake (Moll et al., 1982).

### Annotation of SNP effects of the non-synonymous SNPs using RiceVarMap and IRRI SNP Seek database

The effects of the non-synonymous SNPs in each gene *OsAAP3, OsAAP5* and *OsAAP11* were retrieved from IRRI SNP-Seek Database (https://snpseek.irri.org/) and RiceVarMap (https://ricevarmap.ncpgr.cn) and visualized using MOTIF FINDER (https://www.genome.jp/tools/motif/) and PROTTER tools (https://wlab.ethz.ch/protter/start/)

### Statistical analyses

Two-way ANOVA was carried out in GraphPad Prism version 8 (la Jolla, California, USA) and adjusted P values and levels of significance were computed for variety and treatment effects. Sidak’s multiple comparisons test was performed and graphs were made using GraphPad Prism version 8 (la Jolla, California, USA). Principal Component Analysis (PCA) plots, Correlation matrix and Cluster dendrograms were made using R version 4.2.2 (R Foundation for Statistical Computing, Vienna, Austria).

## RESULTS

### Identification of *AAP3, AAP5 and AAP11* haplotypes from IRRI 3K SNP seek database and the physiological evaluation for response to nitrogen

The IRRI SNP-Seek system (http://www.oryzasnp.org/iric-portal/) provides the single nucleotide polymorphism (SNP) information of 20 million rice SNPs each from 3000 rice genomes project after aligning with the Nipponbare genome. Our objective was to correlate the sequence variation in selected *OsAAPs (OsAAP3, OsAAP5* and *OsAAP11)* with the N response and N use efficiency. The three selected AAPs are previously reported as negative regulators of rice NUE, yield and/or quality (Lu *et al*., 2018, Wang *et al*., 2019, Yang *et al.,* 2023). Haplotype analysis of *OsAAP3, OsAAP5* and *OsAAP11* genes was carried out in SNP seek database (https://snpseek.irri.org/) under the 3K filtered SNP set (Manseuto *et al*., 2017). In *OsAAP3*, one non-synonymous SNP was found in comparison to Nipponbare in the 3024 varieties considered, which led to 388 INDEL positions. In *OsAAP5*, one non-synonymous SNP was found in comparison to Nipponbare in the 3024 varieties considered, which led to 56 INDEL positions, while in OsAAP11, six non-synonymous SNP was found in comparison to Nipponbare in the 3024 varieties considered, which led to 11 INDEL positions. The genotypes of 3K germplasm Indica lines available at IARI were searched for these SNPs and lines with SNPs in *OsAAP3, OsAAP5 and OsAAP*11 and genotypes were selected for further study. Total 15 genotypes: one germplasm accession with SNPs similar to Nipponbare, 13 germplasm accessions with SNP in two or three of the selected AAPs, and MTU1010 were used for physiological evaluation **(Supplementary Table 1, Fig. 1a).**

### Effect of N treatments on observed morpho-physiological traits

The genotypes were grown in hydroponics under three different N treatments (HN: 7mM NO_3_^-^ and 0.5mM NH_4_^+^, HA: 7mM NH_4_^+^ and 0.5mM NO_3_^-^, LN: 0.24mM N) in hydroponics at National Phytotron facility, IARI, New Delhi **(Fig. 2).** The 2-Way ANOVA was performed for the following traits SDW, RDW, LA, PH, CCI, TRL, LRSA, LRV, LRL, TRV, TRSA, MRSA, MRL, MRV, RD, tips, forks, TN, shoot N%, NUE and most of them were significant w.r.to N level, variety and interaction **(Supplementary Table 2-8).** Sidak’s multiple comparisons test showed that treatment means were significantly different among various treatments across the traits. HN vs. LN treatment had the most significant difference for all observed morpho-physiological traits except for root traits such as LRSA and MRL. TN and LA showed better significance among all treatment mean comparisons. The phenotypic performance of the genotypes was compared with the high yielding check genotype MTU1010 **(Supplementary Table 3-8)**. Under HN treatment there was approximately 187% and under HA there was 174% higher SDW in comparison to LN. Similarly, HN showed 50% higher RDW than LN. Genotypes Local, NCS901 A and Bhainsa Mundariya showed highest SDW and RDW in all N treatments while ARC 10581 and Selhi showed lower biomass accumulation (**Fig. 3, Supplementary Table 2)**. In HN and HA, the leaf area was 2-3-fold higher than LN. Highest average leaf area was seen in Local, while Selhi showed lower LA. Local, NCS901 A and Bhainsa Mundariya showed highest leaf area in all N treatments **(Fig. 3(c), Supplementary Table 2)**. Highest average plant height was seen in NCS 901 A, while Dongrem and MTU1010 showed lower plant height. Under LN treatment there was approximately 75% and 78% reduction in plant height and leaf area respectively in comparison to HN treatment **(Fig. 3(d), Supplementary Table 2)**. Under HN and HA treatment there was 3-fold higher CCI in comparison to LN. Genotypes Local, NCS901 A, Bhainsa Mundariya and MTU1010 showed highest average CCI and Selhi recorded the lowest. ARC10799, Local, NCS901 A, Bhainsa Mundariya, Dongrem, NCS603 B, SXC216 and ARC18325 performed on par with MTU1010 **(Fig. 3(e), Supplementary Table 3).** Irrespective of genotype, nitrate nutrition (HN) significantly improved the root growth, root surface area and number of fine root hairs in rice. N deficiency also significantly promoted root proliferation. Genotypic variation in root traits were also significant. Root proliferation also has significant correlation with N uptake. Genotypes Local, NCS901 A and Bhainsa Mundariya showed highest TRL among all N treatments and performed better than MTU1010 based on Sidak’s multiple comparisons test comparing varietal means in TRL, TRSA and TRV. In LRL and LRSA, genotype local showed significantly higher values than MTU1010. Genotypes Local, NCS901 and Bhainsa Mundariya were having statistically significant robust root system which is evident visually in **Fig. 4**. The treatment effects on RD were significant, with HA treatment resulting in higher RD. Genotypes with robust root system Local, NCS901 and Bhainsa Mundariya also showed higher RD. Genotypes Dongrem, Selhi, Zinya Kolamba showed significantly lower and genotype Local showed significantly higher root diameter than MTU1010. Root tips and forks were also highest in Local and significantly higher than that of MTU1010 **(Fig. 3, 5, 6; Supplementary Table 4).** Under HN and HA treatment there was approximately 2-6-fold higher TN in comparison to LN. Genotypes Local, Bhainsa Mundariya and NCS 901 showed highest average TN. Genotypes like ARC10799, Bhu Busi, Dongrem, and ARC18325 performed on par with MTU1010. Under HN and HA treatment there was approximately 2-3-fold higher N concentration in comparison to LN Genotypes Bhainsa Mundariya, Selhi and Local showed highest average shoot N concentration and all genotypes were significantly different from MTU1010 **(Fig 6, Supplementary Table 3).** Total shoot N% and NUE were calculated for all the genotypes which were significantly different among the varieties, treatments and interact and a significant trend was observed which is represented as heat map in **Supplementary figure 1a-b**.

**Fig 3.**
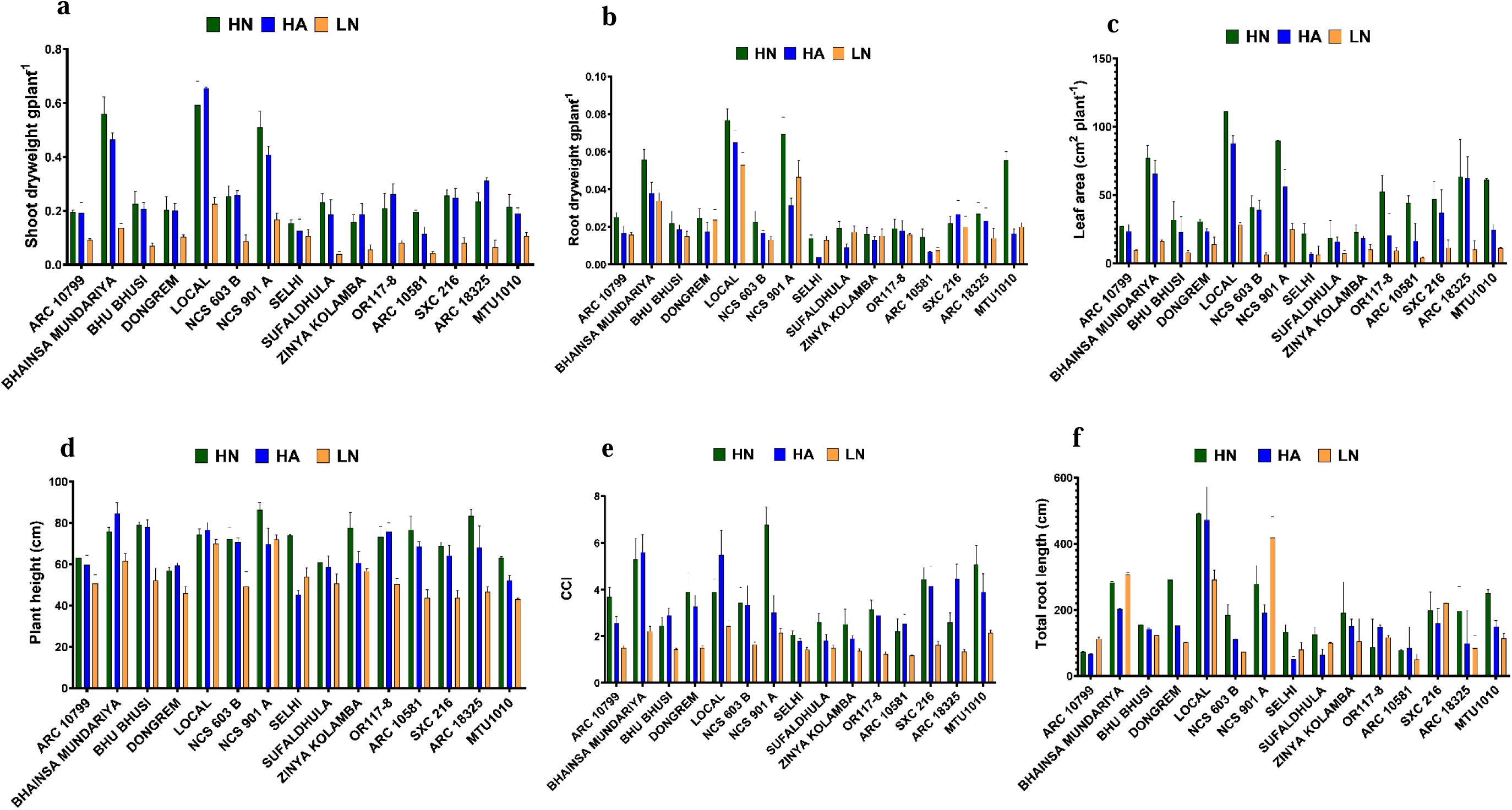
Effect of high nitrate (HN), high ammonium (HA) and nitrogen deficient (LN) conditions on a) shoot dry weight (SDW) b) root dry weight (RDW) c) leaf area (LA) d) plant height (PH) e) chlorophyll content index (CCI) f) total root length (TRL) of rice genotypes belonging to different AAP3, AAP5 and AAP11 haplotypes in hydroponics (HN: 7mM NO_3_^-^ and 0.5mM NH_4_^+^, HA: 7mM NH_4_^+^ and 0.5mM NO_3_^-^, LN: 0.24mM N). Values are mean of three biological replicates (n=3)

**Fig 4.**
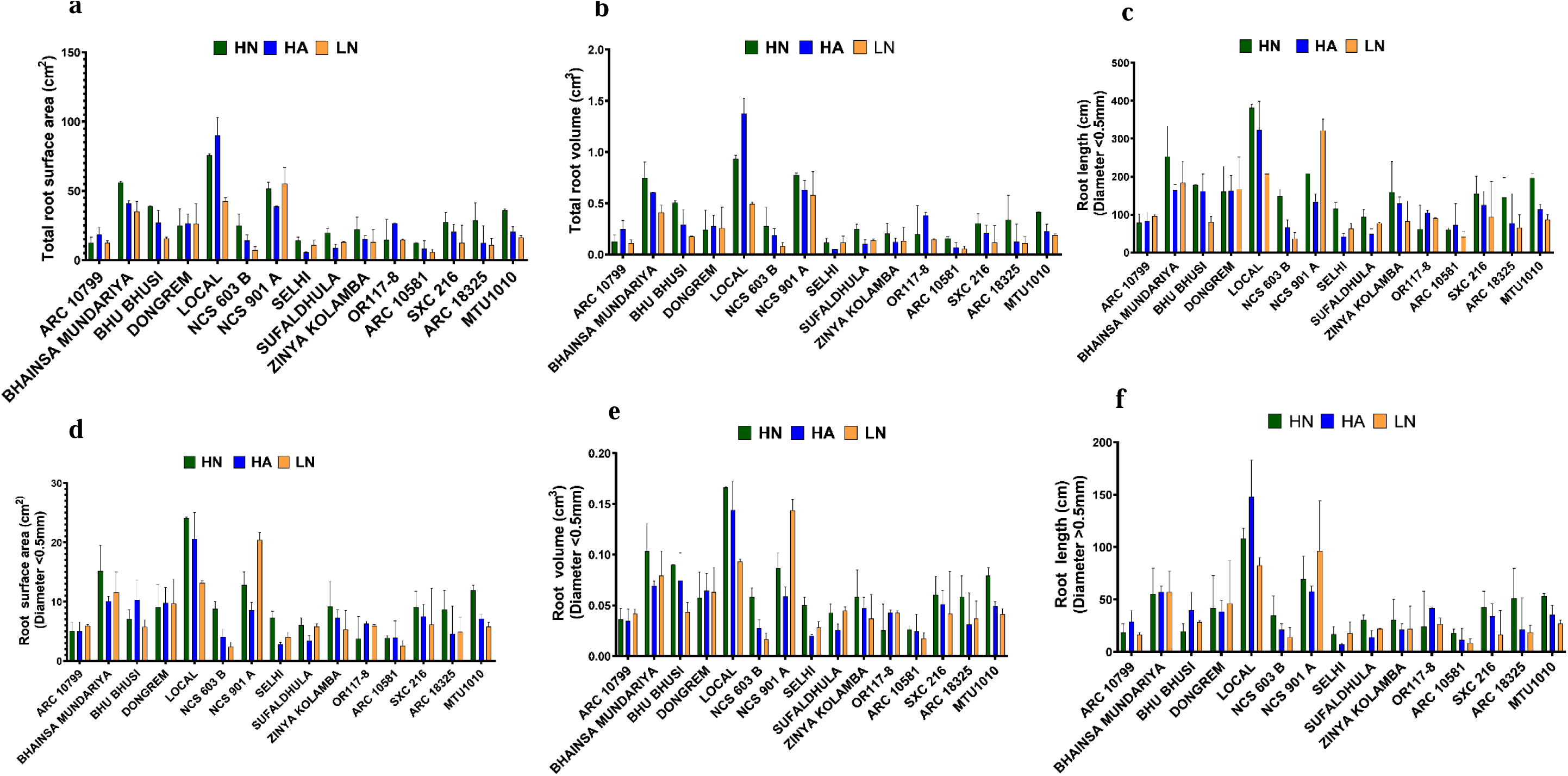
Effect of high nitrate (HN), high ammonium (HA) and nitrogen deficient (LN) conditions on a) total root surface area (TRSA) b) total root volume c) length of lateral root (Diameter≤0.5mm) d) surface area of lateral root (LRSA) e) volume of lateral root (LRV) f) length of main root (Diameter>0.5mm) (MRL) of rice genotypes belonging to different AAP3, AAP5 and AAP11 haplotypes in hydroponics (HN: 7mM NO_3_^-^ and 0.5mM NH_4_^+^, HA: 7mM NH_4_^+^ and 0.5mM NO_3_^-^, LN: 0.24mM N). Values are mean of three biological replicates (n=3).

**Fig 5.**
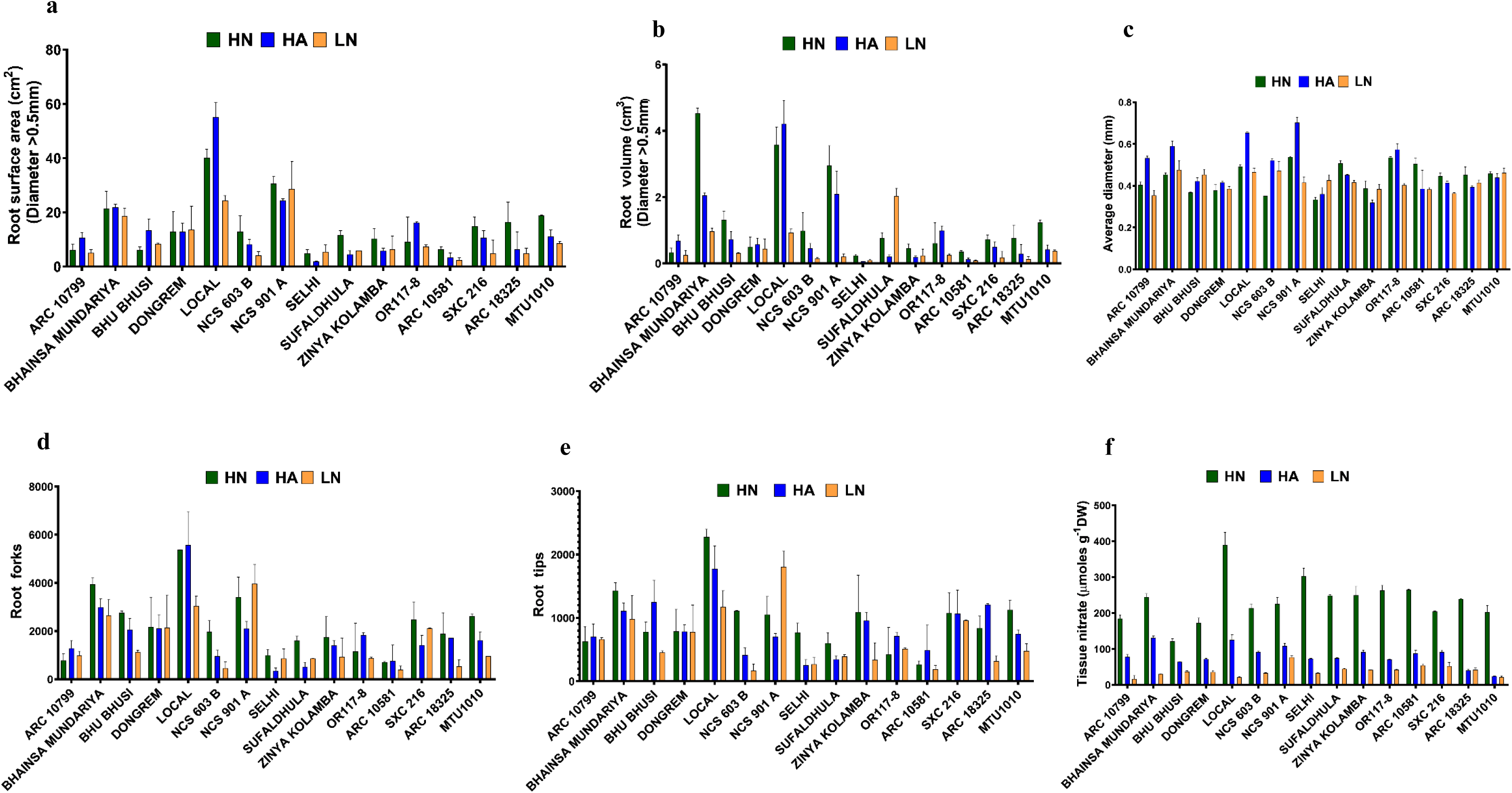
Effect of high nitrate (HN), high ammonium (HA) and nitrogen deficient (LN) conditions on a) surface area of main root (Diameter>0.5mm) (MRSA) b)volume of main root (Diameter>0.5mm) (MRV) c) root diameter (RD) d) root tips (Tips) e) root forks (Forks) f) tissue NO_3_^-^ (TN) of rice genotypes belonging to different AAP3, AAP5 and AAP11 haplotypes in hydroponics (HN: 7mM NO_3_^-^ and 0.5mM NH_4_^+^, HA: 7mM NH_4_^+^ and 0.5mM NO_3_^-^, LN: 0.24mM N). Values are mean of three biological replicates (n=3)

**Fig 6.**
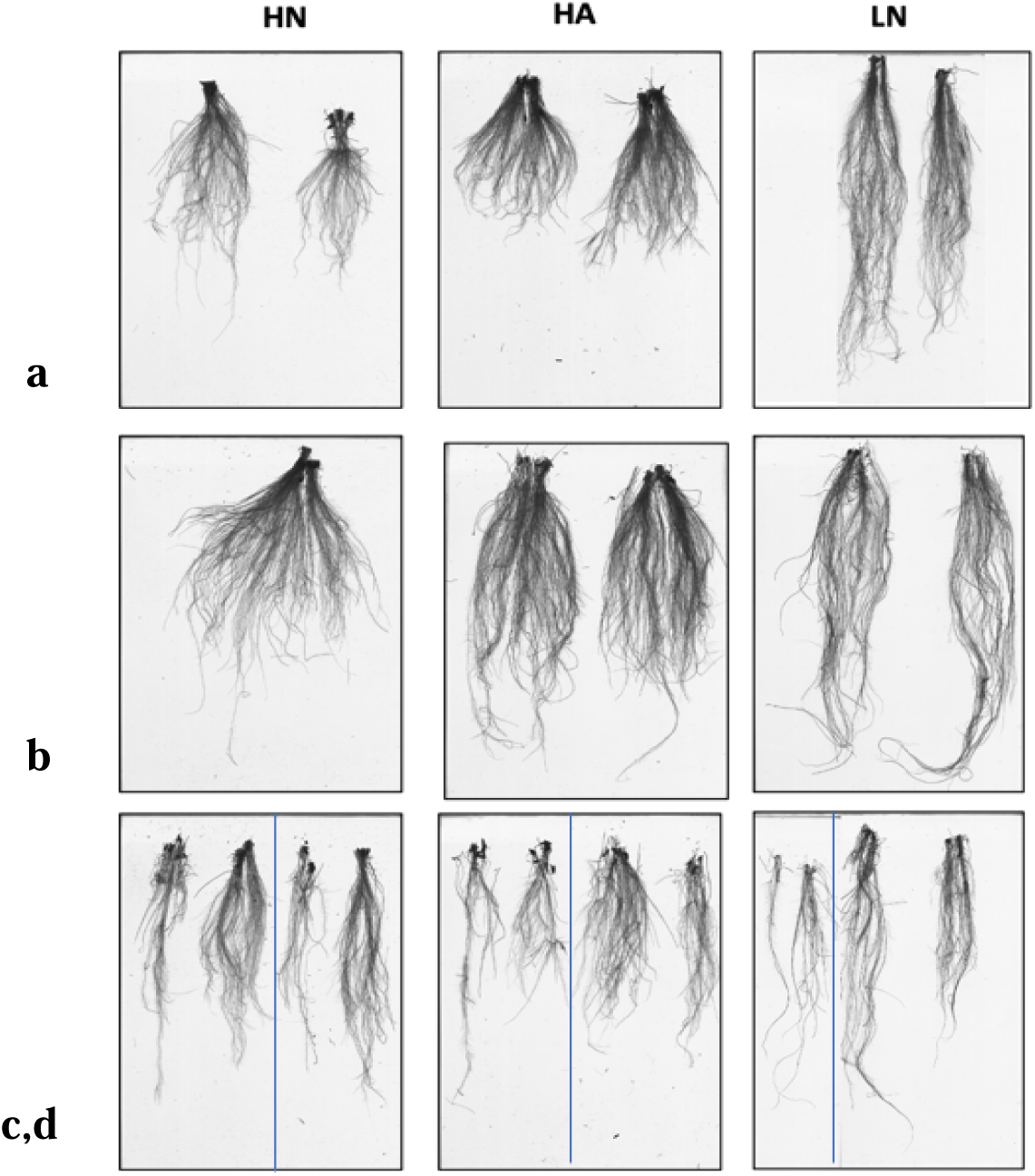
Comparison of root growth of rice genotypes of rice genotypes belonging to different AAP3, AAP5 and AAP11 haplotypes in hydroponics (HN: 7mM NO_3_^-^ and 0.5mM NH_4_^+^, HA: 7mM NH_4_^+^ and 0.5mM NO_3_^-^, LN: 0.24mM N). The images of highly contrasting four genotypes are included: a) BHAINSA MUNDARIYA::(IRGC 60893-1) b) LOCAL::(IRGC 53300-1) c) ARC 18325::IRGC 42399-1) d) MTU1010

### Correlation of the different seedling traits and N treatments in hydroponic culture

Under HN treatment, TN and PH showed no significant correlation with any other traits. CCI was positively correlated with LA, SDW, RDW, MRV, MRL, MRSA and negatively correlated with TN. All the root traits (MRSA, MRL, MRV, TRV, TRL, RD, LRSA, LRV, LRL, TRSA, Tips, Forks), plant biomass (RDW, SDW) and shoot trait such as LA showed significant positive correlation among each other **(Fig.7a)**

**Fig 7.**
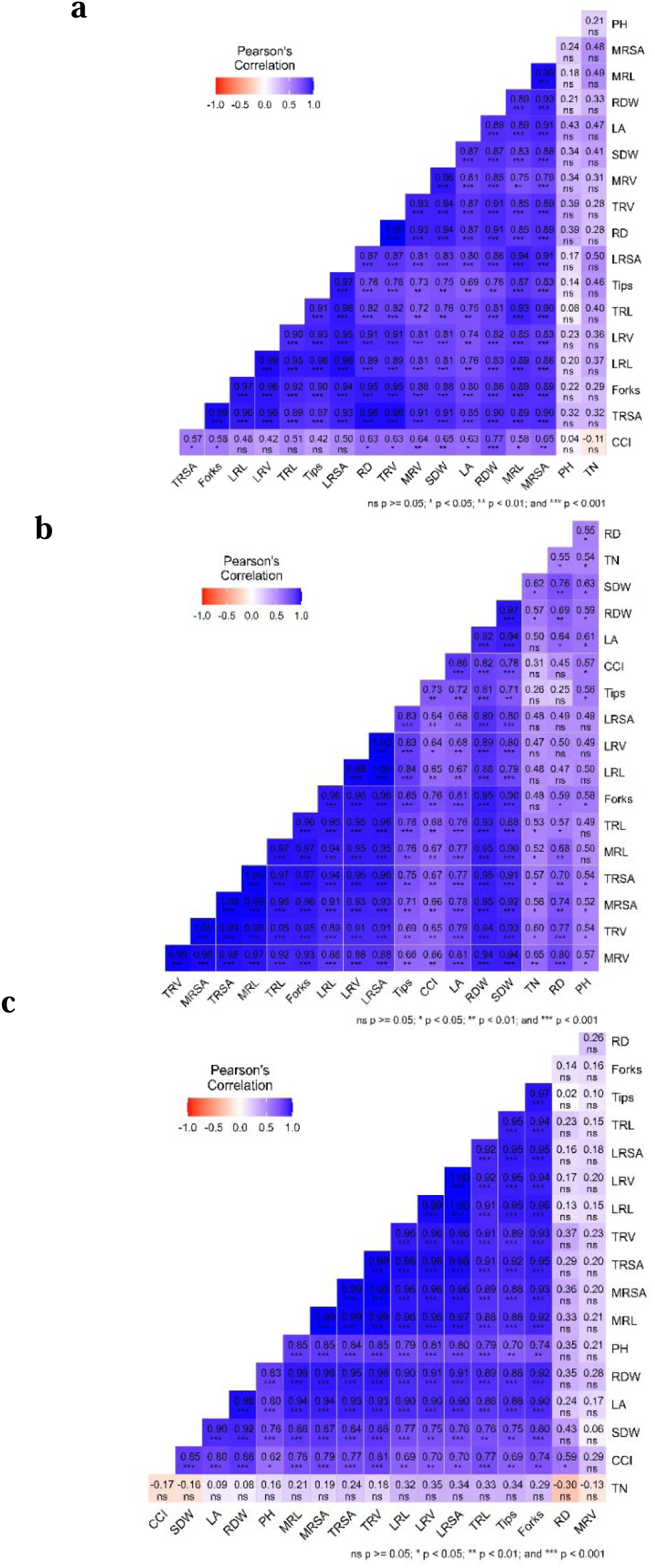
Correlation plot of the different seedling traits in rice genotypes belonging to different AAP3, AAP5 and AAP11 haplotypes in hydroponics grown under a) high nitrate (HN) b) high ammonium (HA) c) and nitrogen deficient (LN) conditions in hydroponic culture. (HN: 7mM NO_3_^-^ and 0.5mM NH_4_^+^, HA: 7mM NH_4_^+^ and 0.5mM NO_3_^-^, LN: 0.24mM N)

Under HA treatment, TN showed no significant correlation with LA, CCI, tips, LRSA, LRV, LRL and forks. RD showed no significant correlation with LRSA, LRV, LRL, tips and CCI. PH had no significant correlation with TRL, LRL, LRV, LRSA, MRL. All other root traits (TRL, LRSA, LRV, LRL, TRV, TRSA, MRSA, MRL, Tips, forks), plant biomass (RDW, SDW), shoot traits (LA), CCI were positively correlated among each other **(Fig.7b)**

Under LN treatment, TN had negative correlation with CCI, SDW, RD and MRV. MRV and RD had no significant correlation with any other traits. All other root traits (TRL, LRSA, LRV, LRL, TRV, TRSA, MRSA, MRL, Tips, forks), plant biomass (RDW, SDW), shoot traits (LA, PH), CCI were positively correlated among each other **(Fig.7c)**

### Principal Component Analysis (PCA) and cluster dendrogram for the observed morpho-physiological traits in the rice seedlings grown in hydroponics

Principal component analysis of 18 traits viz., plant biomass (SDW, RDW), shoot traits (PH, LA), root traits (TRL, TRSA, RD, TRV, LRL, LRV, LRSA, MRL, MRV, MRSA, Tips, Forks), Tissue nitrate content (TN), Chlorophyll Content Index (CCI) of 15 Indica rice accessions grown up to seedling stage in hydroponics to identify traits contributing the most to the genotypic variability in response to HN, HA and LN treatments. Total variability among the genotypes were contributed by PC1 (77.3%) and PC2 (7.2%) in HN; PC1 (79.7%) and PC2 (7.2%) in HA; PC1 (75.6%) and PC2 (10.6%) in LN. The following traits, TRSA, RD, TRV, TRL, LRSA, MRL, LRL, LRV, MRSA, MRV, SDW, RDW, Tips and Forks contribute significantly in HN treatment; TRSA, TRV, MRV, MRSA, SDW, RDW, LRV, TRL, MRL, LRL, LRSA, forks in HA treatment, TRSA, MRSA, MRL, TRV, RDW, LRV, TRL, LRL, LRSA, SDW, CCI, LA, Tips, Forks in LN treatment. The length of the arrows indicates the significance of the traits’ contribution towards the PCA components. **(Fig.8a-c)**

To compare different physiological traits between the rice genotypes and to understand the patterns or similarity among the lines under HN, HA and LN, clustering of genotypes was done. Clustering analysis groups genotypes showing similar physiological traits together. Clustering separated the genotypes into two groups. The second smaller group includes three genotypes, that outperformed others in several physiological traits. Clearly the genotypes were most efficient because they showed robust biomass and root traits. The first group is sub-divided into two groups, the first among them included 7 accessions while the second group contains 5 genotypes including MTU1010. The less efficient genotypes were grouped together based physiological traits suggesting the possibility of improving NUE of these lines by physiological and molecular approaches **(Fig. 8).**

**Fig 8.**
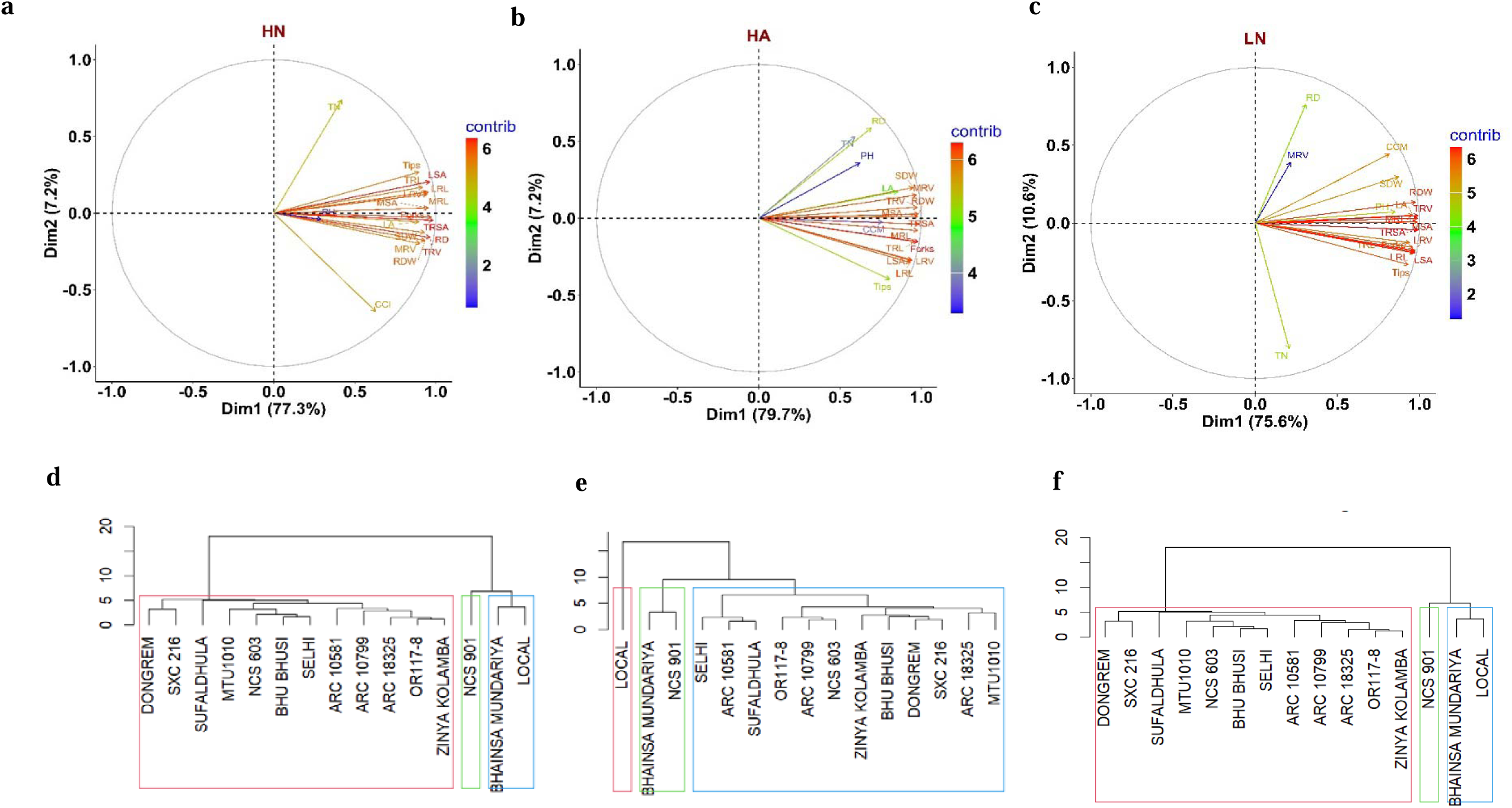
Principal Component Analysis (PCA) plot (a, b, c) and cluster dendrogram (d, e, f) of the different seedling traits in rice genotypes grown under high nitrate (HN) (a, d), high ammonium (HA) (b, e) and nitrogen deficient (LN) (c, f) conditions in hydroponic culture. hydroponics (HN: 7mM NO_3_^-^ and 0.5mM NH_4_^+^, HA: 7mM NH_4_^+^ and 0.5mM NO_3_^-^, LN: 0.24mM N). The length of the arrows indicates the significance of the traits’ contribution towards the PCA components. The traits along the same direction are positively correlated.

### Genotype ranking based on the most contributing traits found across all N treatments

The normalized mean data of all the observed morpho-physiological traits were extracted for the most contributing traits in PC1 and PC2 across all treatments i.e. TRSA, RDW, RD, TN, CCI, Tips and Forks in HN; TRSA, RDW, RD, TN, Forks and Tips in HA; TRSA, TRV, MRSA, RD, TN, CCI in LN **(Supplementary material 1-3)** and ranked based on the normalized mean scores for each genotype and the haplotypes performing on par with MTU1010 and least performing haplotypes in all the treatments were chosen for N assimilation enzyme assays and further proceeded for field evaluation of photosynthetic and yield traits **(Fig. 1b, Supplementary material 4)**

### Effect of N treatments on N assimilation enzyme activity (NR, GS, GOGAT, GDH) and total soluble protein content (TSP) for the selected genotypes

The treatment, varietal and interaction effects were significantly different in all N assimilation enzyme activity in roots and shoots except for root GDH enzyme activity. Sidak’s multiple comparison test showed that the treatment means were significantly different among HN vs LN and HA vs LN for all except GOGAT and GDH activity in roots. The HA vs. HN treatment means were also not significantly different in NR, GS, GDH and TSP activity in roots **(Supplementary Table 9)**. Leaf NR and root NR activity was reduced by 60-70% and 54-58% by N deficiency treatment w.r.to HN and HA treatments. The lowest average leaf NR activity was recorded in OR117-8 followed by MTU1010 and the highest in SXC 216. ARC 10799 and ARC 10581 had higher leaf NR activity in HN and LN whereas SXC 216 in HN and HA. ARC 18325 showed higher leaf NR activity in HN whereas it performed poor in LN showing 86% and 120% reduction w.r.to HN and HA. Contradicting to this MTU1010 showed 4-fold increase in NR activity in leaves under LN w.r.to HN and 68% increase w.r.to HA. Similarly, OR117-8 showed 10% increase in NR activity under LN w.r.to HN. The genotypes ARC 10581 had the highest average root NR activity and ARC 18325 had the least. SXC 216, OR117-8 had similar activity to MTU1010. MTU1010 had the highest root NR activity in HA followed by SXC 216. **(Fig. 9, Supplementary Table 9).** Leaf GS activity and root GS activity was reduced by 16%-29% and 37-40% under HN and HA treatments w.r.to LN. Genotypes ARC 10581 and ARC18325 showed highest average GS activity in in both leaves and roots. But, ARC 18325 though recording the highest leaf GS activity in HA treatment reduced to about 43% in LN conditions. ARC 10581 showed nearly 4 fold increase in LN treatment w.r.to HN. Similar increase was observed in SXC 216, ARC 18325 and MTU1010 root which showed 4-5-fold increase in GS activity in roots in LN treatment w.r.to HN. SXC 216 showed highest GS activity in leaves at HN treatment. OR 117-8 showed decrease in root GS activity of about 34% in Low N conditions w.r.to HN **(Fig 9c, 9d, Supplementary Table 5).** HN and HA treatments had 21% and 31% lower GOGAT activity in leaves and 10% and 77% lower root GOGAT activity in comparison to LN. Genotypes ARC 10581 showed highest GOGAT activity in both leaves and roots. OR 117-8 had 70 % increase in the leaf GOGAT activity in LN w.r.to HN. ARC 18325 and MTU1010 had similar leaf GOGAT activity whereas SXC 216 had the least reduction of about 7% GOGAT in LN w.r.to HN. It showed nearly 4-fold increase in GOGAT root activity under LN conditions w.r.to HN and HA. OR117-8 had the least GOGAT activity in roots similar to MTU1010. **(Fig. 9e, 9f, Supplementary Table 9).** Under HN treatment there was approximately 8% and under HA there was 6% higher leaf GDH activity in comparison to LN. Genotypes ARC10799 showed highest GDH activity and performed significantly better than MTU1010. ARC 10581 had the highest GDH activity in HA treatment and also showed about 4 fold increase in LN w.r.to HN. ARC 18325 showed 5-6-fold increase in leaf GDH activity in LN w.r.to HN and HA. There was approximately 17% in HN and under HA there was 6% lower root GDH activity in comparison to LN. **(Fig. 9g, 9h, Supplementary Table 9**). Genotype ARC 10581 showed highest average root GDH activity and performed significantly better than MTU1010. It increased about 98% in LN treatment w.r.to HN. Approximately, 29-34% and 37-40% of the protein content were reduced under HA and HN treatments in comparison to LN. Every genotype performed on par with MTU1010 except ARC 10581 which showed the least average TSP content in leaves and roots. SXC 216 and OR 117-8 showed the highest average TSP in leaves and root respectively. ARC 10799 had the least amount of reduction in TSP in HA and LN conditions w.r.to HN in leaves and roots. ARC 10581 showed significant increase in TSP in LN wr.to HN in roots. Root TSP was the highest and least in OR117-8 and ARC 18325 in HA. **(Fig 9i, 9j, Supplementary Table 9**).

**Fig 9.**
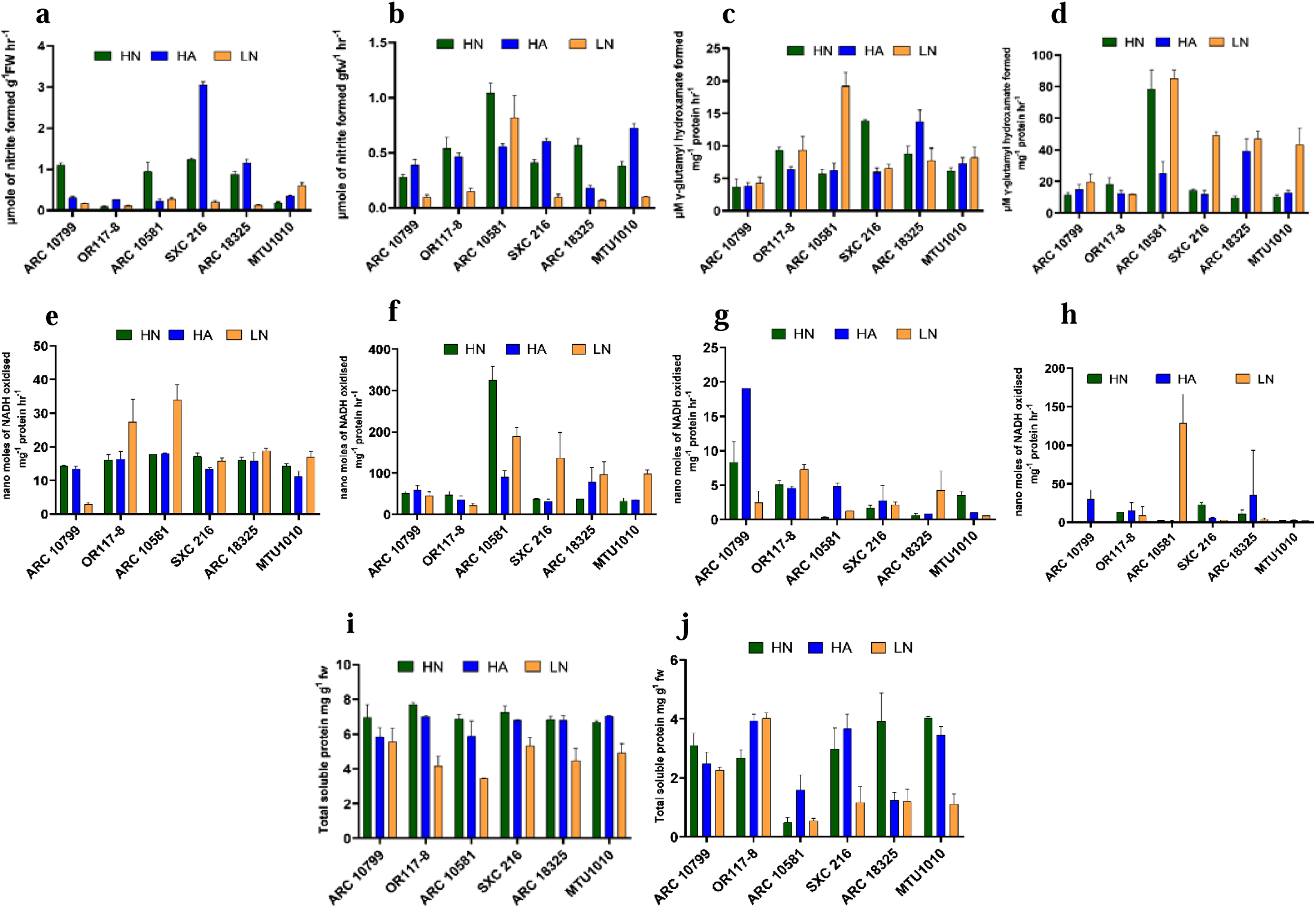
Effect of high nitrate (HN), high ammonium (HA) and nitrogen deficient (LN) conditions on a) nitrate reductase activity in shoots (NR) b) nitrate reductase activity in roots (NR)) c) glutamine synthetase in shoots (GS) d) glutamine synthetase in roots (GS) e) glutamate synthase activity in shoots (GOGAT) f) glutamate synthase activity in roots (GOGAT) g) glutamate dehydrogenase in shoots (GDH) h) glutamate dehydrogenase in roots i) shoot protein content (TSP) j) root protein content (TSP) of rice genotypes belonging to different AAP3, AAP5 and AAP11 haplotypes in hydroponics (HN: 7mM NO_3_^-^ and 0.5mM NH_4_^+^, HA: 7mM NH_4_^+^ and 0.5mM NO_3_, LN: 0.24mM N). Values are mean of three biological replicates (n=3).

### Field evaluation of the selected genotypes to assess the effect of N120 and N0 treatments on the photosynthetic and yield parameters

The selected genotypes **(Supplementary** Fig. 2) were grown in field in Kharif season (June-October 2024) under field conditions with recommended 120 Kg ha^-1^ (N+) and without N fertilizer application (N^-^) at Division of Plant Physiology, ICAR-IARI, New Delhi.

### Effect of N120 and N0 treatments on NUE and yield

Based on 2 Way ANOVA test it is evident that the genotypic variation is significant in all traits except TFW, grain N% and CCI2. The treatment effects are significant in TDW, TW, PW, CCI1, CCI2, PH, CT, PN, PN/Ci, Shoot N%, Total N uptake, and NUE. The varietal and treatment interaction is highly significant except TDW, Tiller No., Panicle No., Shoot N% and Grain N%, **(Supplementary Table 9)**. TFW had nearly 80% reduction in low N conditions and the highest and least values in N120 were recorded in ARC 10581 and MTU1010 respectively. Though ARC 10581 had the highest TFW in N120 it had the least TFW in N0 conditions with 94% reduction. ARC 10799 had the highest TFW in N0 treatment. Similarly, TDW had severe impact in N0 conditions due to 78% reduction **(Fig. 10a, Supplementary Table 9)**. ARC 10581 and ARC 10799 had the highest TDW in N120 and N0 respectively and the least in N120 was observed in MTU1010. Though ARC 10581 performed the best in N120 it had the least TDW in N0 conditions with 94% reduction. OR 117-8 had the least reduction in N0 treatment (59%) **(Fig. 10b, Supplementary Table 9)**. Tiller No., Panicle No. and grain weight are highly influenced by genotypic variation rather than treatment effects **(Fig. 10c, 10f, 10g, Supplementary Table 9)**. TW faced 83% reduction in low N treatment. TW was the highest and least in ARC 18325 and MTU1010 in N120 and in ARC 10581 and MTU1010 in N0 conditions. The highest significant reduction of about 83% in N0 treatment was observed in ARC 18325 and OR 117-8 **(Fig. 10d, Supplementary Table 9)**. Similarly, OR 117-8 had a 100% significant reduction in PW in N0 **(Fig. 10e, Supplementary Table 9)**. CCI1 and CCI2 had 17% and 50% reduction in N0. CCI1 was the highest in ARC 18325 in both conditions and the least in OR 117-8 in N sufficient treatment and in ARC 10581 in N0 treatment. SXC 216 maintained same level of CCI1 in both treatments and OR 117-8 had 9% increase in N0. MTU1010 faced severe reduction of about 37% in N deficiency **(Fig. 11a, Supplementary Table 9)**. CCI2 was observed the highest in ARC 18325 and the least in MTU1010 in N120 and the same was observed in MTU1010 and OR 117-8 in N0 conditions. 45-80% of reduction was observed in OR 117-8, ARC 10799, SXC 216 and ARC 18325 whereas MTU1010 had the least reduction of about 7% in low N conditions **(Fig. 11b, Supplementary Table 9)**. CT was increased about 1.5.% in N scarce conditions. MTU1010 maintained the least CT in both conditions. OR 117-8 had 0.2% reduction in CT in N0. ARC 10581 had the highest percentage increase in N0 of about 5% in N0 conditions followed by ARC 18325 and ARC 10799. SXC 216 maintained almost same levels of CT in both conditions **(Fig. 11c, Supplementary Table 9)**. PH had 28% reduction in N deficiency. The highest and least PH was observed in ARC 18325 and MTU1010 respectively in N120 and the same was observed in ARC 10581 and MTU1010 in N0 conditions. ARC 10799, ARC 10581, SXC 216 had only about 20-25% reduction in N0 whereas ARC 18325 had 43% reduction in N0 conditions. MTU1010 though recorded in least PH in N120 it maintained the PH in N0 conditions with only 5% reduction. PH had 28% reduction in N deficiency. The highest and least PH was observed in ARC 18325 and MTU1010 respectively in N120 and the same was observed in ARC 10581 and MTU1010 in SXC 216 in N0 conditions. ARC 10799, ARC 10581, SXC 216 had only about 20-25% reduction in N0 whereas ARC 18325 had 43% reduction in N0 conditions. MTU1010 though recorded in least PH in N120 it maintained the PH in N0 conditions with only 5% reduction **(Fig. 11d, Supplementary Table 9)**. PN increased about 20% in N0 conditions in which SXC 216 and OR 117-8 had about 84% and 76% increase in PN respectively in No conditions. MTU1010 had the highest PN in N120 conditions but faced reduction of about 30% in N deficient conditions followed by ARC 10799 which recorded the lowest in both conditions. ARC 10581 and ARC 18325 showed 27% and 47% increase in PN in low N conditions **(Fig. 12a, Supplementary Table 9)**. gs had 0.26% increase in N0 conditions. The highest and least was recorded in MTU1010 and ARC 10581 in N120 conditions and the same was observed in OR117-8 and ARC 10799 in N0 conditions. MTU1010 showed the highest reduction of about 70% in N0 conditions whereas OR 117-8 showed about 3 fold increase in N0 conditions followed by SXC 216 and ARC 10581 which showed about 2 fold increase **(Fig. 12b, Supplementary Table 9)**. Ci was the highest and the least in OR117-8 and ARC 10581 in both treatments. Ci increased about 30% higher in ARC 10581 whereas ARC 18325 and SXC 216 had 3-8% increase in Ci in low N. ARC 10799 followed by MTU1010 had the highest reduction in Ci in N0 treatment of about 20% and 13% respectively **(Fig. 12c, Supplementary Table 9)**. TR increased about 4% in N0 treatment. OR 117-8 had the highest TR in N0 treatment with a 84% increase w.r.to N120 followed by ARC 10581 and SXC 216 which had 65% and 45% increase in N0 treatment. MTU1010 and SXC 216 recorded the highest and least TR in N120 treatment. **(Fig. 12d, T Supplementary Table 9)**. ARC 10581 and MTU1010 had the highest and least iwUE in N120 treatment and ARC 10799 and OR 117-8 in N0 treatment. MTU1010 showed 145% increase and ARC 10799 had 84% increase in iWUE in N0. OR 117-8, ARC 10581 and SXC 216 showed nearly 15-45% of reduction in iWUE in N0 conditions **(Fig. 12e, Supplementary Table 9)**. 80% reduction in PN/Ci was observed in N0 conditions. The highest and least were observed in MTU1010 and ARC 10799 in N120 and in ARC 18325 and ARC 10799 in N0 conditions. SXC 216, OR 117-8, ARC 18325 showed 42-56% increase in PN/Ci in N0 conditions and MTU1010 had the highest reduction of about 18% in N0 conditions. **(Fig. 12f, Supplementary Table 9)**. Ci/Ca increased about 1-31% in ARC 18325, SXC 216, OR 117-8 and ARC 10581 and reduced to about 13% and 21% in MTU1010 and ARC 107999 respectively in N scarce conditions. The highest and least were recorded in MTU1010 and ARC 10581 in N120 treatment. **(Fig. 12g, Supplementary Table 9)**. N % in shoots reduced about 18% in N0 treatment. Shoot N% was the highest and least in ARC 10581 and OR 117-9 in N120 and in ARC 10799 and OR 117-8 in N0 treatment. ARC 18325 faced a significant reduction of 42% under N deficiency **(Fig. 12h, Supplementary Table 9)**. Grain N% is non-significant for varietal, treatment and interaction effects **(Fig. 12i, Supplementary Table 9).** 80% reduction in Total N uptake was observed in N0 treatment and the highest significant reduction was observed in ARC 18325. Though SXC 216 showed the least N uptake in N120 conditions it showed nearly 2-fold increase in N0 conditions. ARC 10799, ARC 10581 and MTU1010 performed on par under optimum N supply and showed 20-50% reduction under low N conditions. OR 117-8 though performing better than other cultivars in N120 faced a reduction of 93% under scarce N supply **(Fig. 12j, Supplementary Table 9)** NUE was significantly higher in SXC 216 under low N condition whereas all other cultivars except ARC 18325 showed 20-40% increase under low N supply **(Fig. 12k, Supplementary Table 9).** NUtE was significantly higher of about 10 fold in SXC 216 under low N conditions followed by ARC 1058.

**Fig 10.**
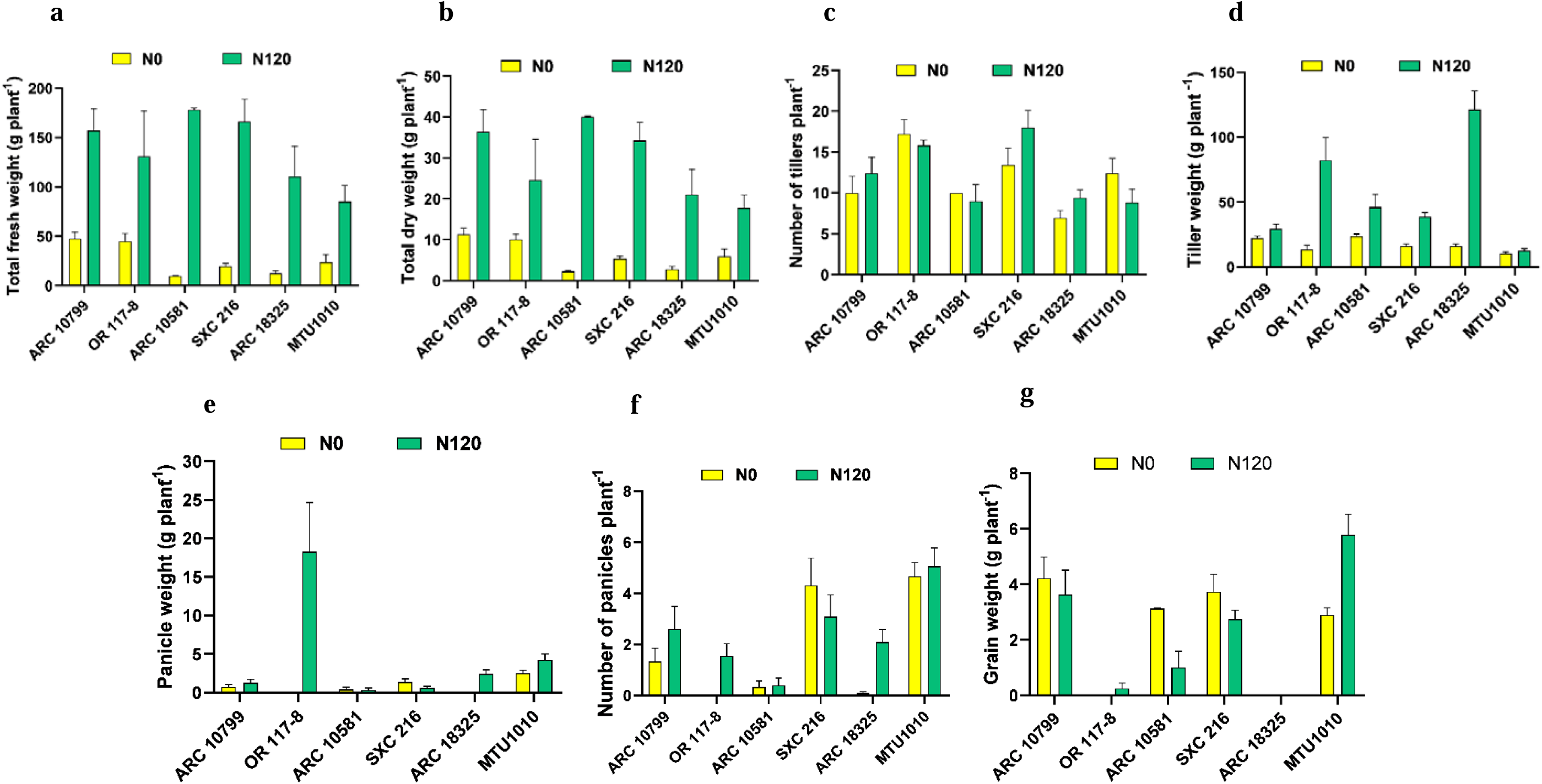
Effect of high nitrogen (120 kg ha^−1^ applied N: N120), and nitrogen deficient (no applied N: N0) field conditions on biomass and yield traits a) total fresh weight (TFW) b) total dry weight (TDW) c) number of tillers (Tiller No.) d) tiller weight (TW) e) panicle weight (PW) f) number of panicles (Panicle No.) g) grain weight (GW) of rice genotypes belonging to different AAP3, AAP5 and AAP11 haplotypes. Values are mean of three biological replicates (n=3).

**Fig. 11.**
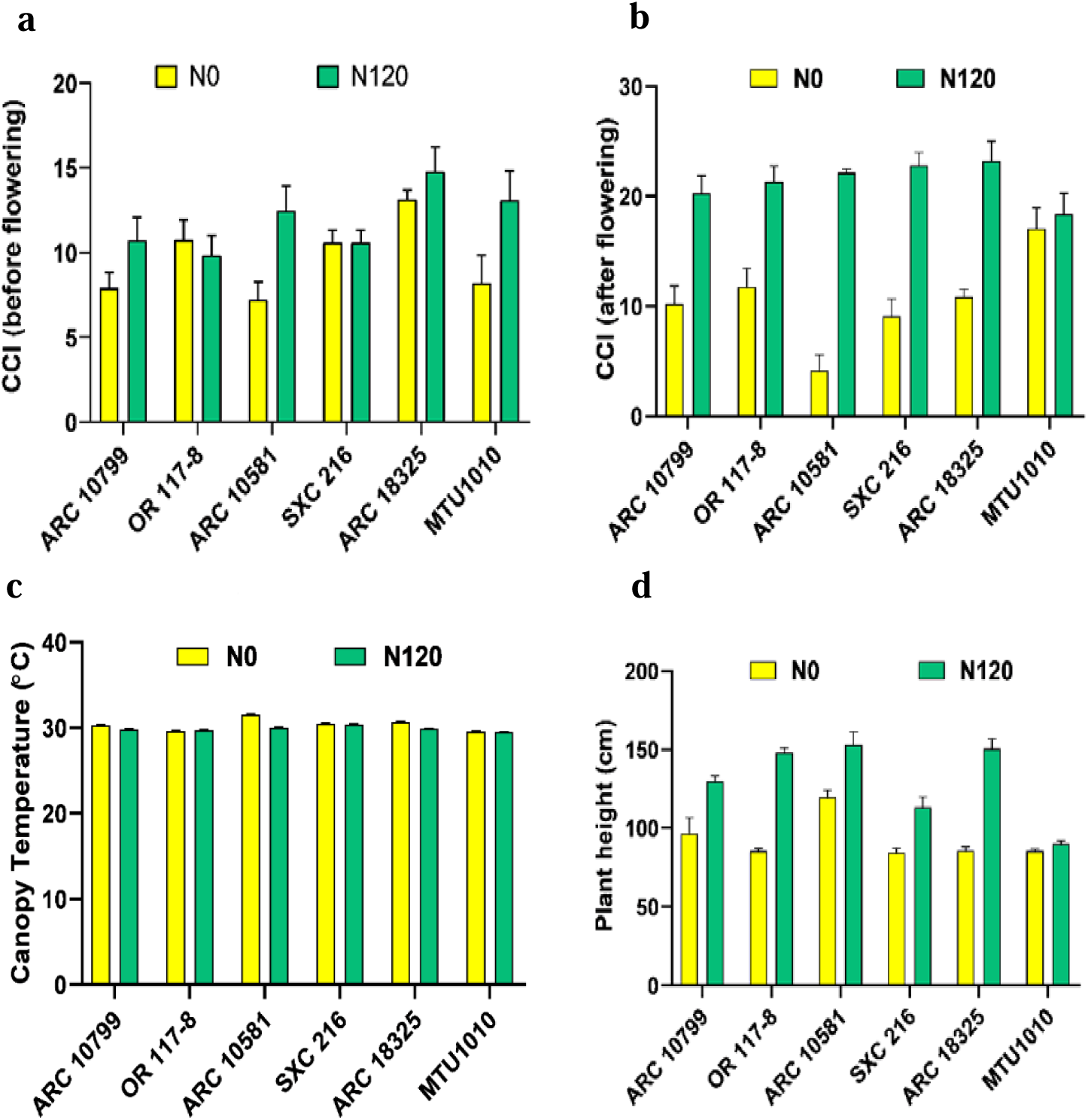
Effect of high nitrogen (120 kg ha^−1^ applied N: N120), and nitrogen deficient (no applied N: N0) field conditions on a) CCI before flowering b) CCI after flowering c) plant height (PH) d) canopy temperature (CT) of rice genotypes belonging to different AAP3, AAP5 and AAP11 haplotypes. Values are mean of three biological replicates (n=3).

**Fig. 12.**
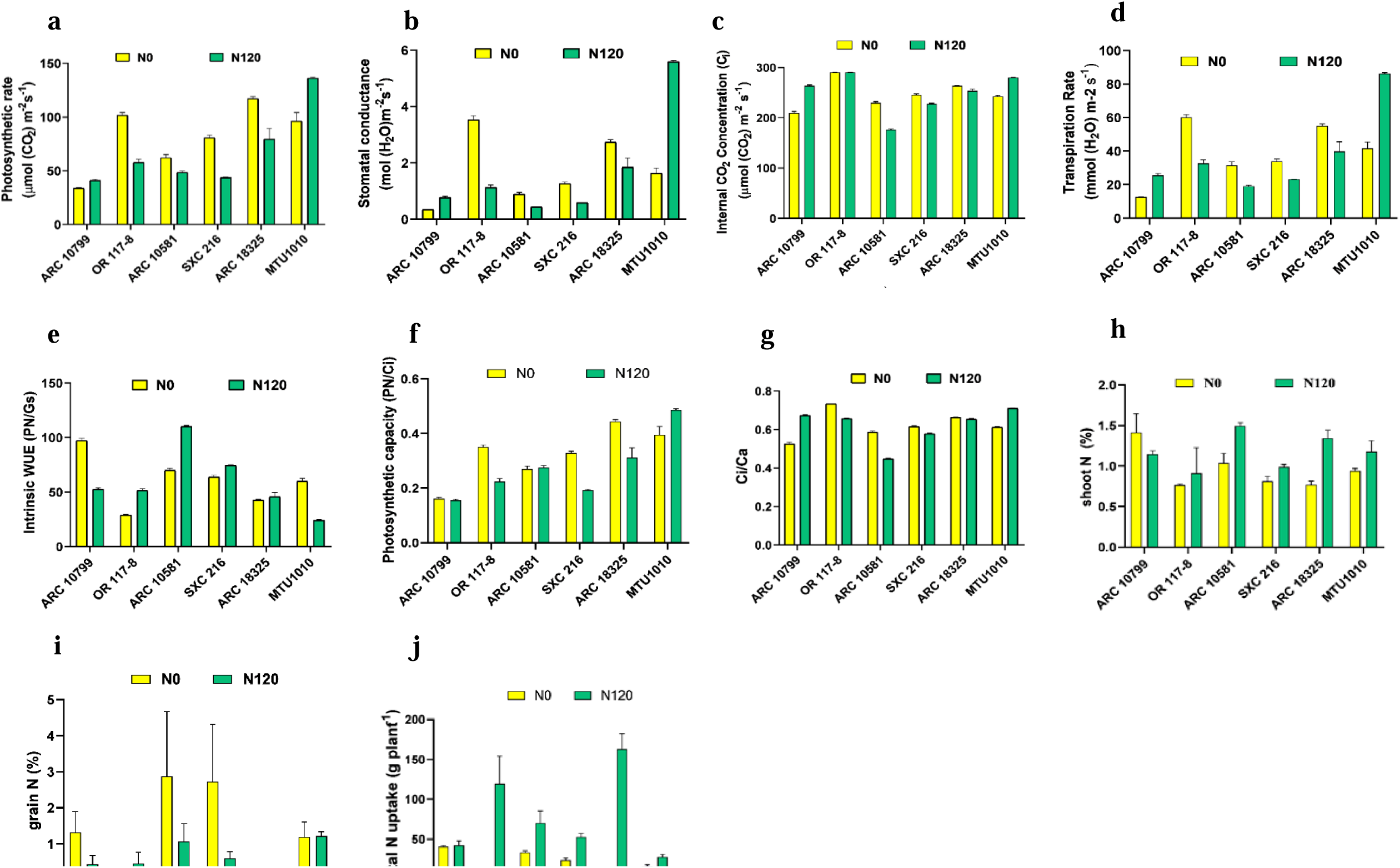
Effect of high nitrogen (120 kg ha^−1^ applied N: N120), and nitrogen deficient (no applied N: N0) field conditions on a) Photosynthetic rate (Pn) b) Stomatal conductance (gs) c) Internal CO_2_ Concentration (Ci) d) Transpiration rate (TR) e) Intrinsic Water Use Efficiency (WUE) f) Pn/Ci g) Ci/Ca h) shoot N% i) grain N% j) Total N uptake k) NUE l) NUtE of rice genotypes belonging to different AAP3, AAP5 and AAP11 haplotypes. Values are mean of three biological replicates (n=3).

ARC 18325 had ill filled grains under both N conditions and OR 11-8 faced the same under low N availability **(Fig. 12l, Supplementary Table 9).** Based on Sidak’s multiple comparison test comparing the treatment means, it is evident that N120 vs. N0 treatment is highly significant in TFW, TDW, TW, CCI2, PH, CT, Ci, TR, iWUE, Ci/Ca in the genotype ARC 10799. OR 117-8 had high significant differences in the traits such as TW, PW, CCI2, PH, PN,. Gs, TR, iWUE, PN/Ci, Ci/Ca. ARC 10581 had significant treatment effects on TFW, TDW, CCI1, CCI2, PH, CT, Ci, TR, iWUE and Ci/Ca. In SXC 216 it was observed in TFW, TDW, TW, CCI2, PH, CT, PN, gs, Ci, TR, iWUE, PN/Ci, Ci/Ca, NUE and NUtE. The treatment means comparison were significant for the following traits in ARC 18325 such as TFW, TDW, TW, CCI2, PH, CT, PN, gs, Ci, TR, PN/Ci, shoot N%, and Total N uptake. The treatment effects were not significant in most traits for MTU1010 except CCI1, PN, gs, Ci, TR, iWUE, PN/Ci, Ci/Ca and TNC **(Supplementary Table 9)**

### Annotating the SNP effects of the non-synonymous mutations in OsAAP3, OsAAP5 and OsAAP11

The effects of the non-synonymous SNPs in each gene *OsAAP3, OsAAP5* and *OsAAP11* were retrieved from IRRI SNP-Seek Database (https://snpseek.irri.org/) and RiceVarMap (https://ricevarmap.ncpgr.cn/) **(Table 1).** The frequency of the non-synonymous SNP (T➔ C) located in chromosome 6 and position 21191370 in the *OsAAP3* gene causes a missense variation leading to an amino acid change from valine to alanine in position 379 which is found in the functional transmembrane domain of the protein in the cytoplasmic side **(Fig 13a)**. SIFT and Polyphen-2 scores were 0.51 and -0.148 respectively which predict that the SNP effect is benign and tolerated **(Table 1).** The reference allele is present in 95.80% of the Indica population and the SNP is found in 4.20% of the Indica varieties. By comparing the polymorphic variations (only SNPs) of the selected genotypes and Nipponbare in the RiceVarMap it was found that the genotypes ARC 10799, ARC 10581 had this non-synonymous SNP whereas ARC 18325, SXC 216 and OR 117-8 had the primary allele (T) in this position. For the gene OsAAP5, the reference allele (T) is the secondary allele in Indica group and is present only in 2.50% of the population whereas the non-synonymous SNP (G) is present in 97.30% of the Indica population. The SNP causes a change in the amino acid from glutamate to alanine at position 350 in the protein sequence which is present in the functional transmembrane domain in the cytoplasmic region **(Fig 13b)** and the polyphen2 and SIFT scores indicate the SNP to be possibly damaging **(Table 1).** Through putative genotype-phenotype associations found in external GWAS sources in RiceVarMap it was found that this SNP (vg0138118647) is strongly associated with yield with significant Likelihood Ratio p-value (LR-P) value of 7.03E-19 across all sub-populations which is far below the strict Bonferroni threshold (5* 10^-8) which indicates strong association with yield (Xie et al., 2015). This SNP was observed in ARC 10799, ARC 10581, SXC 216 and OR117-8. The gene OsAAP11 has 6 non-synonymous SNP positions which are mostly tolerated except the SNP vg1104799528 (T➔C) which is probably damaging as detected by polyphen2 scores and this SNP causes an amino acid change from methionine to threonine at 36 position which is near the membrane spanning region of the protein **(Table 1).** All the 6 SNPs are found at or near the membrane spanning domain of the trans-membrane helices and these 6 SNPs were present in ARC 10799, ARC 10581 and SXC 216. OR 117-8 and ARC 18325 lacked these SNPs in OsAAP11 **(Table 1**, **Fig. 13c)**

**Fig 13.**
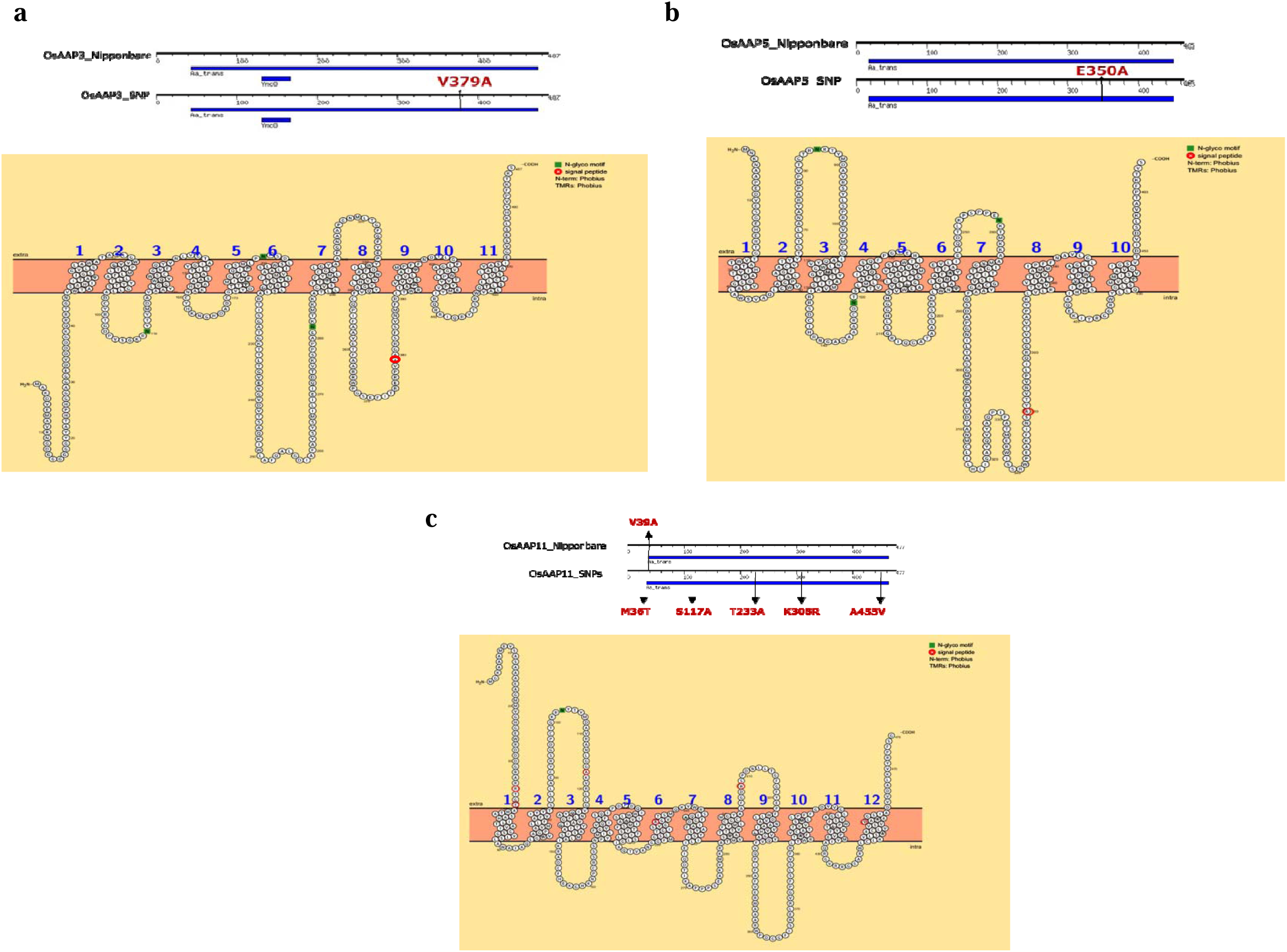
Visualization of the position of non-synonymous SNPs in the amino-acid sequences of a) AAP3 b) AAP5 and c) AAP11 using Motif Finder and Protter

## Discussion

Haplotype-based breeding is an important approach for the generation of high performing, custom-made rice genotypes and is a way out in meeting future food and nutritional demands (Abbai et al., 2019). We identified AAP3, AAP5 and AAP11 haplotypes from IRRI 3K SNP seek database to understand the variation in physiological response of these plants to N doses. Total 15 genotypes: one germplasm accession with SNPs similar to Nipponbare, 13 germplasm accessions with SNP in two or three of the selected AAPs, and a high yielding cultivar, MTU1010 were used for physiological evaluation. Genotypes were grown in hydroponics under three different N treatments, HN:7mM Nitrate:0.5mM Ammonium, HA:7mM Ammonium:0.5mM Nitrate, LN:0.24mM Ammonium Nitrate ((Jagadhesan et al., 2020).

Shoot biomass and root biomass accumulation of rice genotypes were significantly different, with regard to N level, variety and their interaction. Genotypes Local, NCS901 and Bhainsa Mundariya showed highest shoot biomass accumulation in all N treatments and performed better than MTU1010. N deficiency in rice lead to yellowing of lower leaves, reduced the leaf area and shoot growth. The reduction in leaf area and biomass is more evident in low NUE genotype in seedling stage and mature plants (Sathee et al., 2019). Response to long-term N deficiency is reflected as change in physiology and biochemistry of shoot and root tissues. Including reduction of growth and rate of photosynthesis, the remobilization of N from old, mature tissues to actively growing ones and the accumulation of abundant photo-damage-protecting anthocyanin pigments (Jagadhesan et al., 2020).

Application of N fertilizer increases the chlorophyll content (Pramanik and Bera, 2013). In rice, chlorophyll content is widely used to ascertain N demand and to increase grain yield and also as an indicator of NUE (Huang et al., 2008). Jagadhesan et al., 2020 found considerable reduction in chlorophyll content of the rice crop over two seasons subjected to N starvation. N availability also influences the composition of component chlorophyll pigments and carotenoids, and their ratios. Significant decline in chlorophyll and carotenoid content under low N conditions was also found in wheat genotypes (Nouriyani et al., 2012).

Local, NCS901 and Bhainsa Mundariya showed highest leaf area, shoot biomass and root traits in all N treatments and performed better than MTU1010. These lines can be used as donors for NUE associated traits in breeding programs. Clustering analysis groups genotypes showing similar physiological traits together. Clustering separated the genotypes into two groups. The second smaller group includes three genotypes, that outperformed others in several physiological traits. Clearly the genotypes were most efficient because they showed robust biomass and root traits. The first group is sub-divided into two groups, the first among them included 7 accessions while the second group contains 5 genotypes including MTU1010. The less efficient genotypes were grouped together based physiological traits suggesting the possibility of improving NUE of these lines by physiological and molecular approaches. Thus, the genotypes which were performing on par with MTU1010 and the least performing genotypes in the seedling stage were chosen based on their ranking in their phenotypic performance in the most contributing traits across all N regimes and further validated by performing N assimilation assays and field evaluation for NUE and yield traits.

Nitrogen use efficiency (NUE) is the product of N uptake efficiency (NUpE) and N utilization efficiency (NUtE), which is the optimal combination between N assimilation efficiency (NAE) and N remobilization efficiency (Chardon et al., 2010). It is evident that genes involved in root architecture, N uptake, assimilation, N-storage and re-translocation as well as genes involved in regulation of these processes, such as transcription factors, have a critical impact on NUE (Xu et al., 2012). NUtE is determine by the activity of component enzymes as described in earlier sections. Enzyme nitrate reductase (NR) induced by nitrate availability and is the rate limiting enzyme in N assimilation pathway (Vincentz et al., 1993). In the current study, also we found huge variation in NR activity between two N+ and N-treatments. Our current and earlier data (Jagadhesan et al., 2020) hints that in low NUE lines, NR is having low affinity for nitrate and hence the threshold level is high (Vijayalaxmi et al., 2015). Hakeem et al., (2012) also found that high NUE varieties displayed higher NR activity and consistently maintained it even with variable N levels. A study on wheat cultivars exhibiting contrasting NUE reported GS activity correlates with N remobilization and filling of grains (Kichey et al., 2007). In rice, *OsGS1.1* was found to be crucial for the remobilization of glutamine via phloem for the enhanced growth rate and grain filling (Obara et al., 2001, Sathee et al., 2021). Thus, it may be inferred that high NUE genotypes can utilize remobilized ammonia as alternate N source with help of more GS1 activity in N deficient conditions.

As discussed in earlier sections, amino acid transporters including AAPs have multiple functions and more profoundly, AAP’s role in modulating growth and development in rice is well established (Zhao et al., 2012). *OsAAP1* and *OsAAP4* controls tillering and yield positively, while *OsAAP3*, *OsAAP5*, *OsAAP7* etc regulate rice tillering negatively (Ji et al., 2020, Fang et al., 2021, Jin et al., 2024). The negatively regulation of axillary bud growth and tillering in *OsAAP3*, *OsAAP5*, *OsAAP7* is mainly due to transport of basic amino acids Arg and Lys and by crosstalk with N, auxin, and cytokinin signaling mechanisms. Mutation in *OsAAP6*, *OsAAP10* and *OsAAP11* improved grain quality in rice (Peng et al., 2014, Wang et al., 2020c). The amylose contents of *OsAAP6* and *OsAAP10* mutants were low (Peng et al., 2014, Wang et al., 2020) while in *OsAAP11* mutant higher quality was achieved by low concentrations of grain amino acids, low grain protein and high viscosity of cooked rice (Yang et al., 2023). Mutants of *OsLHT1* also display higher grain protein, lower amylose along with lower grain yield (Guo et al., 2020). Amino acid transporter-like gene 13 (*OsATL13*) transports phenylalanine (Phe) and methionine (Met), and improves axillary bud growth and grain rice yield, while capable of modulating grain amylose and protein, positively.

NUE is a complex trait critical for improving rice productivity under variable N supply. Amino acid permeases such as OsAAP3, OsAAP5, and OsAAP11 have been shown to function as negative regulators of NUE in Japonica rice (Lu et al., 2018; Wang et al., 2019), yet their roles in Indica rice remain to be fully elucidated. In this study, we evaluated selected Indica genotypes under contrasting N regimes—seedlings were tested under high nitrate (HN), high ammonium (HA), and low N (LN) conditions, and field performance was assessed under N120 (optimum N) and N0 (low N). Our aim was to relate performance differences to the SNP status in these three genes, thereby clarifying their influence on N uptake, assimilation, yield potential, and N stress adaptation. ARC 10799 had non-synonymous SNP in OsAAP3, OsAAP5 and OsAAP11. ARC 10799 showed better NR activity in seedling stage in all treatments and recorded the highest average GDH activity in leaves significantly in HA treatment followed by enhanced TFW, TDW, TW in N scarce conditions in field maintaining moderate number of panicles, higher PW in in both optimum and scarce N supply along with moderate reduction in NUE and NUtE under low N availability. It also had lower reduction in PH and Total N uptake in N0 conditions but the least PN and PN/Ci in both treatments and decreased CCI2, Ci/Ca and gs in low N supply along with increased iWUE and CT. This shows that ARC 10799 has enhanced N uptake with improved nitrogen metabolism under stress in seedling stage. Field evaluation indicates that it has stronger biomass production with maintenance of productive tillering capacity under N stress with efficient use of water. Despite strong N assimilation under N stress, it represented poor photosynthetic activity and stomatal performance reflecting its adaptive response to stress conditions. This confirms that the SNPs in OsAAP3, OsAAP5, and OsAAP11 positively influence N assimilation and biomass production under N stress but does not have high photosynthetic NUE. ARC 10581 had non-synonymous SNP in OsAAP3, OsAAP5 and OsAAP11. ARC 10581 had significantly higher NR activity, GS activity, GOGAT activity in both leaves and roots across all treatments with significant GDH activity in HA treatment in leaves and highest average activity in roots. ARC 10581 had highest biomass accumulation (TFW, TDW), iWUE in N120 conditions and increased growth stability (PH), CCI1, PN, PN/Ci, gs, TR in N0 conditions along with a maintenance of panicle number and increased panicle weight under low N supply. It also showed partial increase in NUE and high-fold hike of NUtE with moderate reduction in total N uptake under N deficiency. This suggests that ARC 10581 performed the best in terms of N uptake, N assimilation and N remobilization and N cycling under N stress in seedling stage and maintained the highest biomass in optimum N conditions. Under low N conditions, the higher retention of chlorophyll content, growth stability and photosynthetic rate and a partial increase in panicle weight and NUE alongwith shoot up of NUtE suggest that it maintained better physiological efficiency under low N through improved stress adaptation which becomes trade-off to biomass accumulation under N stress to enhance the reproductive potential of the plant. Thus, non-synonymous SNPs in OsAAP3, OsAAP5 and OsAAP11 in common enhances the stress resilience in genotypes ARC 10581 and ARC 10799 despite different growth and yield strategies which could be influenced by other genotypic features. SXC 216 had non synonymous SNPs in OsAAP5 and OsAAP11. SXC 216 showed highest average leaf NR activity, GS activity in HN and the highest average TSP content in leaves. It showed 4-fold higher GOGAT activity in LN conditions in leaves. Under field evaluation, there was reduction in tiller weight in N0 along with maintenance of CT in both N treatments, least reduction in PH, increase in TR, Ci/Ca, gs, PN and highest reduction in iWUE, in low N supply. It performed extra-ordinarily showing significant high-fold increase under low N treatment in terms of PW, NUE, NUtE and total N uptake. This shows that SXC 216 shows better N reduction capacity via improved N uptake and assimilation under high N stress and efficient N recycling and stress recovery in low N along with excellent N partitioning into proteins. For the stabilization of growth and yield under low N supply, the plant has prioritized photosynthetic capacity via increased stomatal regulation and transpiration rate which comes at the cost of higher water loss which significantly improved the yield and N use and utilization efficiency of the plant. Though the non-synonymous SNPs in OsAAP5 and OsAAP11 improved the N assimilation and N remobilization under stress, the lack of significant N response and biomass accumulation in optimum N conditions might be due to the absence of OsAAP3 SNPs which may regulate better N uptake and biomass accumulation in N optimum conditions as observed in ARC 10799 and ARC 10581. OR 117-8 had non synonymous SNPs only in OsAAP5. It showed the least average NR activity in leaves and GOGAT activity in roots but it showed 70% increase in GOGAT activity in LN conditions w.r.to HN in leaves and had the highest average TSP content in roots. OR 117-8 though having higher TW in optimum N conditions it performed the least in terms of TW and PW under low N. It had cooler canopy (CT), higher PN, gs, PN/Ci, Ci/Ca, least iWUE in N0 treatment and least shoot N% in both treatments (Fig 14a). Data suggests an N induced regulation of physiological parameters (Fig 14a). The changes in NUE parameters was at least in part, related to variation in biomass, plant height, photosynthesis and pigment content related parameters (Fig 14 b). The comparatively dissimilar trends in shown by different genotypes (Fig 14 a) suggests possibility of high or low correlation trait-wise, offering an opportunity to identify significant contrasts in a divergent set of AAP haplotypes suggesting their probable role in yield determination and N stress adaptation in rice.

**Fig. 14.**
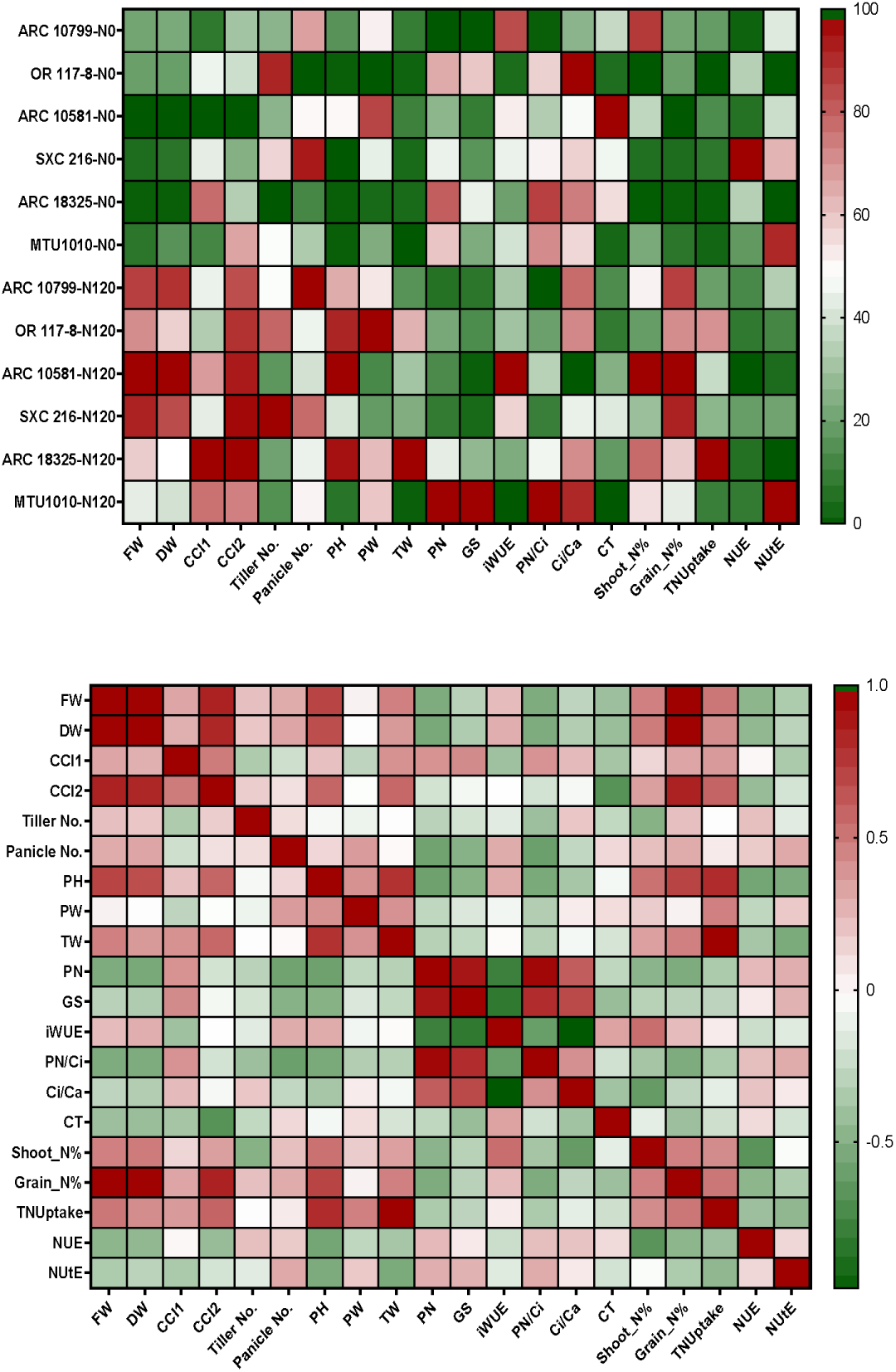
Heat maps representing (a) normalized values of physiological parameters showing the effect of high nitrogen (120 kg ha^−1^ applied N: N120), and nitrogen deficient (no applied N: N0) field conditions in rice genotypes belonging to different AAP3, AAP5 and AAP11 haplotypes (b) Pearson correlation matrix showing relationship between physiological and NUE parameters.

Nearly 100% reduction in total N uptake and NutE confirms that SNPs in OsAAP5 alone enhances physiological adaptation to stress but not enough for stress resilience due to inefficient conversion to biomass or reproductive output under stress conditions. We can interpret that OR 117-8 did not perform well under optimum conditions also owing to the absence of OsAAP3 SNPs which may partially contribute to better N uptake under N supply. Also, the poor NR activity in seedling stage supports this statement whereas the higher GOGAT activity and TSP content compensates for the poor N uptake and improves N remobilization due to the presence of OsAAP5 SNPs. ARC 18325 had no non-synonymous SNPs and was similar to Japonica Nipponbare cultivar. ARC 18325 showed poor NR activity in LN in leaves and the least average NR activity in roots. But it showed highest average GS activity in leaves and roots and better leaf GDH activity under LN treatment. It also showed least root TSP content under HA. ARC 18325 had higher, CCI1, PH and CCI2 under N120. It also showed higher PN, Ci, PN/Ci and the least TW, 100% reduction in panicles and their weight in low N supply. ARC 18325 had nearly 100% reduction in total N uptake under low N supply along with poor grain filling, NUtE and reduced NUE. ARC 18325 despite having poor NR activity and low TSP content in roots due to lack of efficient N uptake, higher GS and GDH activity compensated for this loss by metabolizing the internally available N more efficiently as a stress recovery response. But under field evaluation, this did not translate to effective biomass accumulation and yield under stress conditions. This confirms the role of the non-synonymous SNPs in OsAAP3, OsAAP5 and OsAAP11 in which non-synonymous SNPs in OsAAP3 may lead to enhance N uptake in seedling stage under different N regimes and in field conditions under optimum and low N levels whereas the non-synonymous SNPs in OsAAP5 is required for the better stress adaptation by enhancing physiological responses such as PN, gs, PN/Ci under N stress but alone it does not sustain good yield performance under N deficient conditions. The non-synonymous SNPs in OsAAP11 is crucial to convert the stress adaptation in to higher yield via better N remobilization to reproductive organs under N stress. OR 117-8 lacked OsAAP5 and OsAAP11. MTU1010 was the high yielding check genotype which was grown along with the selected genotypes. It showed the highest PN, Gs and PN/Ci under optimum N conditions but significant reduction in iWUe and PN/Ci along with low tiller weight under both N regimes and partial decrease in NutE and total N uptake under low N supply. MTU1010’s performance resembles that of genotypes lacking SNPs in OsAAP3, OsAAP5, and OsAAP11 (like ARC 18325 and OR 117-8) — suggesting that it might lack all three SNPs. However, its strong growth under high N points to an efficient NUE under both conditions, potentially through alternative NUE pathways not linked to OsAAP3, OsAAP5, or OsAAP11. Interestingly, the trend observed in the NUE calculated in seedling stage was similar to those proven in field evaluation and this proves that in general, genotypes with non-synonymous mutation in AAP3, AAP5 and AAP11 in comparison to Japonica showed better growth and N content. Indica rice can thus be improved by knocking out editing the potential target genes which are responsible for the negative regulation of NUE growth. The previous reports and current findings opens new avenues and insights for improving rice yield, quality and NUE by changes in AAP haplotypes that occur naturally or created by precise genome editing.

## Acknowledgement

The authors are thankful to the ICAR-Indian Agricultural Research Institute for funding and for providing the necessary facilities. HSi acknowledges ICAR for the fellowship support received during the study.

## Author contributions

HSi conducted the hydroponic experiment, collected data and determined all NUE related data. KR conducted the field experiment and determined yield parameters. RP helped in the determination of photosynthetic and fluorescence traits and thermal imaging. LS, SKJ, RE and VC provided resources and finalised the experiments. HSi and LS finalized figures, analysed data and wrote the first draft. LS conceived the idea and revised the manuscript.

