## Supplementary material for "Association of haplotypes of AAP family amino acid transporters with nitrogen response and nitrogen use efficiency in rice grown under hydroponics and field conditions": 3.Supplementary table1, 7-9

**Supplementary Table 1. Details of Indica rice genotypes belonging to different AAP haplotypes and check genotype MTU1010 selected for physiological evaluation of 30 days old seedlings grown in hydroponics under three different N treatments** **(HN: 7mM NO_3_^-^ and 0.5mM NH_4_^+^, HA: 7mM NH_4_^+^ and 0.5mM NO_3_^-^, LN: 0.24mM N)**

| S.No. | Name: Accession | IRIS ID | Box Code | Subpopulation | Country |
| --- | --- | --- | --- | --- | --- |
| 1 | ARC 10799::IRGC 12631-1 | IRIS 313-9313 | AE78 | indx | India |
| 2 | BHAINSA MUNDARIYA::IRGC 60893-1 | IRIS 313-11596 | AO84 | indx | India |
| 3 | BHU BHUSI::IRGC 70803-1 | IRIS 313-11823 | AR32 | ind2 | India |
| 4 | DONGREM::IRGC 6688-1 | IRIS 313-10544 | AY25 | ind2 | India |
| 5 | LOCAL::IRGC 53300-1 | IRIS 313-11478 | AN59 | ind2 | India |
| 6 | NCS 603 B::IRGC 62377-1 | IRIS 313-11643 | AP38 | ind1B | India |
| 7 | NCS 901 A::IRGC 62568-1 | IRIS 313-11650 | AP45 | ind2 | India |
| 8 | SELHI::IRGC 52760-1 | IRIS 313-11460 | AN41 | ind2 | India |
| 9 | SUFALDHULA::IRGC 46698-2 | IRIS 313-11372 | BH14 | ind2 | India |
| 10 | ZINYA KOLAMBA::IRGC 52402-1 | IRIS 313-11453 | AN34 | ind2 | India |
| 11 | ARC 10581::IRGC 12514-1 | IRIS 313-10671 | AZ58 | indx | India |
| 12 | SXC 216::IRGC 35175-1 | IRIS 313-11176 | BH4 | indx | India |
| 13 | OR 117-8::IRGC 39680-2 | IRIS 313-11242 | BF70 | ind2 | India |
| 14 | ARC 18325::IRGC 42399-1 | IRIS 313-11286 | BG21 | indx | India |
| 15 | MTU1010 | High yielding check genotype | | | |

**Supplementary Table 7. Details of the Sidak’s multiple comparison test for the comparison of treatment means (HN vs. HA, HN vs. LN, HA vs. LN and the 2-Way ANOVA test for the varietal, treatment and interaction significances and their percentage of total variation for the morpho-physiological traits 1) Shoot Dry Weight (SDW) 2) Root Dry Weight (RDW) 3) Leaf Area (LA) 4) Plant Height (PH) 5) Chlorophyll Content Index (CCI) 6) Total Root Length (TRL) 7) Total Root Surface Area (TRSA) 8) Total Root Volume 9) Length Of Lateral Root (Diameter≤0.5mm) 10) Surface Area Of Lateral Root (LRSA) 11) Volume Of Lateral Root (LRV) l2) Length Of Main Root (Diameter>0.5mm) (MRL) 13) Surface Area Of Main Root (Diameter>0.5mm) (MRSA) 14)Volume Of Main Root (Diameter>0.5mm) (MRV) 15) Root Diameter (RD) 16) Root tips (Tips) 17) Root forks (Forks) 18)Tissue NO_3_^-^ (TN)**

| S. No. | Traits | Sidak’s multiple comparison test (Adjusted P value, significance) | | | 2-Way ANOVA  (Percentage of total variation, P value, significance) | | |
| --- | --- | --- | --- | --- | --- | --- | --- |
|  |  | HN vs. HA | HN vs. LN | HA vs. LN | Genotype | Treatment | Interaction |
| 1. | SDW | 0.6680 (ns) | <0.0001 (****) | <0.0001 (****) | 46.25 <0.0001 (****) | 28.92 <0.0001 (****) | 11.21 <0.0001 (****) |
| 2. | RDW | <0.0001 (****) | <0.0001 (****) | 0.9993 (ns) | 65.04 0.0002 (***) | 7.368 <0.0001 (****) | 9.513 0.0001 (****) |
| 3. | LA | 0.0001 (***) | <0.0001 (****) | <0.0001 (****) | 42.45 <0.0001 (****) | 32.96 <0.0001 (****) | 14.01 0.0117 (*) |
| 4. | PH | 0.0003 (***) | <0.0001 (****) | <0.0001 (****) | 27.74 <0.0001 (****) | 36.85 <0.0001 (****) | 16.58 0.0001 (***) |
| 5. | CCI | 0.2743 (ns) | <0.0001 (****) | <0.0001 (****) | 19.22 <0.0001 (****) | 21.21 <0.0001 (****) | 11.23 <0.0001 (****) |
| 6. | TRL | 0.0025 (**) | 0.0052 (**) | 0.9919 (ns) | 66.92 <0.0001 (****) | 4.431 0.0011 (**) | 16.19 0.0134 (*) |
| 7. | TRSA | 0.0727 (ns) | 0.0001 (***) | 0.0890 (ns) | 70.57 <0.0001 (****) | 5.701 0.0002 (***) | 11.36 0.1200 (ns) |
| 8. | TRV | 0.2895 (ns) | <0.0001 (****) | 0.0003 (***) | 71.61 <0.0001 (****) | 6.454 <0.0001 (****) | 14.00 0.0009 (***) |
| 9. | LRL | 0.0362 (*) | 0.0096 (**) | 0.9446 (ns) | 56.44 <0.0001 (****) | 5.623 0.0068 (**) | 15.29 0.3949 (ns) |
| 10. | LRSA | 0.0814 (ns) | 0.0565 (ns) | 0.9981 (ns) | 59.40 <0.0001 (***) | 3.494 0.0327 (*) | 15.84 0.2903 (ns) |
| 11. | LRV | 0.0354 (ns) | 0.0419 (*) | 0.9998 (ns) | 60.70 <0.0001 (****) | 3.912 0.0171 (*) | 15.65 0.2292 (ns) |
| 12. | MRL | 0.9359 (ns) | 0.2735 (ns) | 0.5915 (ns) | 70.17 <0.0001 (****) | 1.906 0.2419 (ns) | 11.90 0.3446 (ns) |
| 13. | MRSA | 0.8295 (ns) | 0.0037 (**) | 0.0307 (*) | 71.68 <0.0001 (****) | 3.651 0.0032 (**) | 12.12 0.0923 (ns) |
| 14. | MRV | 0.0009 (***) | <0.0001 (****) | <0.0001 (****) | 55.51 <0.0001 (****) | 10.00 <0.0001 (****) | 28.46 <0.0001 (****) |
| 15. | RD | 0.0002 (***) | 0.0385 (*) | <0.0001 (****) | 43.49 <0.0001 (****) | 8.811 <0.0001 (****) | 39.85 <0.0001 (****) |
| 16. | TIPS | 0.4234 (ns) | 0.0010 (***) | 0.0483 (*) | 49.73 <0.0001 (****) | 6.917 0.0012 (**) | 23.37 0.0288 (*) |
| 17. | FORKS | 0.0784 (ns) | 0.0010 (**) | 0.3148 (ns) | 64.76 <0.0001 (****) | 5.851 0.0014 (**) | 12.09 0.3566 (ns) |
| 18. | TN | 0.0001 (***) | <0.0001 (****) | <0.0001 (****) | 7.420 <0.0001 (****) | 81.76 <0.0001 (****) | 9.675 <0.0001 (****) |

**Supplementary Table 8. Details of the Sidak’s multiple comparison test for the comparison of treatment means (HN vs. HA, HN vs. LN, HA vs. LN) and the 2-Way ANOVA test for the varietal, treatment and interaction significances and their percentage of total variation for the N assimilation enzymes (Nitrate reductase (NR), Glutamine Oxoglutarate Aminotransferase/Glutamate Synthase (GOGAT), Glutamine synthetase (GS), Glutamate dehydrogenase (GDH) and the total soluble protein content (TSP) in both shoots and roots**

| S. No. | Traits | Sidak’s multiple comparison test (Adjusted P value, significance) | | | 2-Way ANOVA  (Percentage of total variation, P value, significance) | | |
| --- | --- | --- | --- | --- | --- | --- | --- |
|  |  | HN vs. HA | HN vs. LN | HA vs. LN | Genotype | Treatment | Interaction |
| 1. | Leaf- NR | 0.0001 (***) | <0.0001 (****) | <0.0001 (****) | 36.35 <0.0001 (****) | 15.30 <0.0001 (****) | 47.70 <0.0001 (****) |
| 2. | Root – NR | 0.0899 (ns) | <0.0001 (****) | <0.0001 (****) | 44.29 <0.0001 (****) | 25.03 <0.0001 (****) | 27.07 <0.0001 (****) |
| 3. | GS – leaf | 0.2584 (ns) | 0.0076 (**) | <0.0001 (****) | 29.18 <0.0001 (****) | 4.344 <0.0001 (****) | 60.31 <0.0001 (****) |
| 4. | GS – root | 0.0537 (ns) | <0.0001 (****) | <0.0001 (****) | 50.15 <0.0001 (****) | 18.86 <0.0001 (****) | 27.61 <0.0001 (****) |
| 5. | GOGAT – leaf | 0.3020 (ns) | 0.0002 (***) | <0.0001 (****) | 41.98 <0.0001 (****) | 9.260 <0.0001 (****) | 40.69 <0.0001 (****) |
| 6. | GOGAT – root | 0.0001 (***) | 0.4910 (ns) | <0.0001 (****) | 55.02 <0.0001 (****) | 6.056 <0.0001 (****) | 33.40 <0.0001 (****) |
| 7. | GDH – leaf | 0.0006 (***) | 0.9292 (ns) | 0.0002 (***) | 47.08 <0.0001 (****) | 6.585 <0.0001 (****) | 42.68 <0.0001 (****) |
| 8. | GDH – root | 0.8088 (ns) | 0.1629 (ns) | 0.5024 (ns) | 17.71 0.0125 (*) | 3.868 <0.1521 (ns) | 61.85 <0.0001 (****) |
| 9. | TSP – leaf | 0.0070 (**) | <0.0001 (****) | <0.0001 (****) | 7.536 0.0004 (***) | 71.45 <0.0001 (****) | 11.84 0.0003 (***) |
| 10. | TSP- root | 0.6864 (ns) | <0.0001 (****) | <0.0001 (****) | 41.02 <0.0001 (****) | 16.01 <0.0001 (****) | 35.83 <0.0001 (****) |

**Supplementary Table 9. Details of the Sidak’s multiple comparison test for the comparison of treatment means and the 2-Way ANOVA test for the varietal, treatment and interaction significances and their percentage of total variation for the morpho-physiological traits observed in field conditions 1) Total fresh weight (TFW) 2) Total Dry Weight (TDW) 3) Number of tillers (Tiller No.) 4) Tiller weight (TW) 5) No. of Panicles (Panicle No.) 6) Panicle weight (PW) 7)Grain weight (GW) 8) CCI before flowering (CCI1) 9) CCI after flowering (CCI2) 10) Plant height (PH) 11) Canopy temperature (CT) 12) Photosynthetic rate (Pn) 13) Stomatal conductance (gs) 14) Internal CO_2_ concentration (Ci) 15) Transpiration rate (TR) 16) intrinsic Water Use Efficiency (iWUE) 17) Pn/Ci 18) Ci/Ca 19) Shoot N% 20) Grain N% 21) Total N uptake 22) NUE 23) NUtE**

| S No. | Traits | Sidak’s multiple comparison test (Significance)  (N0 vs. N120) | | | | | | 2-Way ANOVA  (Percentage of total variation, significance) | | |
| --- | --- | --- | --- | --- | --- | --- | --- | --- | --- | --- |
|  |  | ARC 10799 | OR 117-8 | ARC 10581 | SXC 216 | ARC 18325 | MTU1010 | Genotype | Treatment | Interaction |
| 1. | TFW | *** | ns | *** | **** | ** | ns | 7.627 (ns) | 58.66 (ns) | 5.866 (****) |
| 2. | TDW | *** | ns | *** | **** | ** | ns | 10.89 (*) | 54.93 (****) | 7.325 (ns) |
| 3. | Tiller No. | ns | ns | ns | ns | ns | ns | 43.78 (****) | 0.3171 (ns) | 9.264 (ns) |
| 4. | TW | ns | **** | ns | ns | **** | ns | 21.44 (****) | 9.407 (****) | 18.21 (****) |
| 5. | Panicle No. | ns | ns | ns | ns | ns | ns | 14.07 (****) | 0.6014 (ns) | 1.517 (ns) |
| 6. | PW | ns | **** | ns | ns | ns | ns | 5.214 (****) | 1.905 (***) | 6.175 (****) |
| 7. | GW | ns | ns | ns | ns | ns | ns | 24.35 (**) | 0.4642 (ns) | 3.839 (ns) |
| 8. | CCI1 | ns | ns | * | ns | ns | * | 12.22 (**) | 7.002 (**) | 7.107 (ns) |
| 9. | CCI2 | *** | *** | ** | **** | **** | ns | 2.167 (ns) | 39.37 (****) | 10.03 (**) |
| 10. | PH | **** | **** | * | *** | **** | ns | 24.59 (****) | 41.98 (****) | 15.57 (****) |
| 11. | CT | **** | ns | **** | ns | **** | ns | 18.77 (****) | 5.923 (****) | 7.190 (****) |
| 12. | Pn | ns | **** | ns | **** | **** | **** | 67.51 (****) | 4.814 (****) | 21.39 (****) |
| 13. | Gs | ns | **** | ns | ** | **** | **** | 54.61 (****) | 0.0002 (ns) | 42.68 (****) |
| 14. | Ci | **** | ns | **** | **** | ** | **** | 68.65 (****) | 0.06 (ns) | 29.96 (****) |
| 15. | TR | ** | **** | ** | * | *** | **** | 59.44 (****) | 0.1194 (ns) | 35.15 (****) |
| 16. | iWUE | **** | **** | **** | *** | ns | **** | 59.92 (****) | 0.01970 (ns) | 37.91 (****) |
| 17. | Pn/Ci | ns | **** | ns | **** | **** | *** | 70.38 (****) | 5.619 (****) | 16.26 (****) |
| 18. | Ci/Ca | **** | **** | **** | **** | ns | **** | 57.86 (****) | 0.04 (ns) | 41.10 (****) |
| 19. | Shoot N% | ns | ns | ns | ns | * | ns | 25.47 (**) | 11.23 (**) | 16.28 (ns) |
| 20. | Grain N% | ns | ns | ns | ns | ns | ns | 25.08 (ns) | 12.59 (ns) | 16.92 (ns) |
| 21. | Total N uptake | ns | *** | ns | ns | ***** | ns | 15.80 (****) | 11.16 (****) | 20.03 (****) |
| 22. | NUE | ns | ns | ns | *** | * | ns | 11.43 (**) | 5.668 (***) | 9.443 (**) |
| 23. | NUtE | ns | ns | ns | ns | ns | ns | 25.25 (**) | 0.0001743 (ns) | 8.118 (ns) |

**( P value style - p value < 0.0001 (****), 0.001-0.005(***), <0.005(**), <0.05(*), >0.05 (ns))**
