## Supplementary material for "Association of haplotypes of AAP family amino acid transporters with nitrogen response and nitrogen use efficiency in rice grown under hydroponics and field conditions": 4.1Supplementary figure 1-5

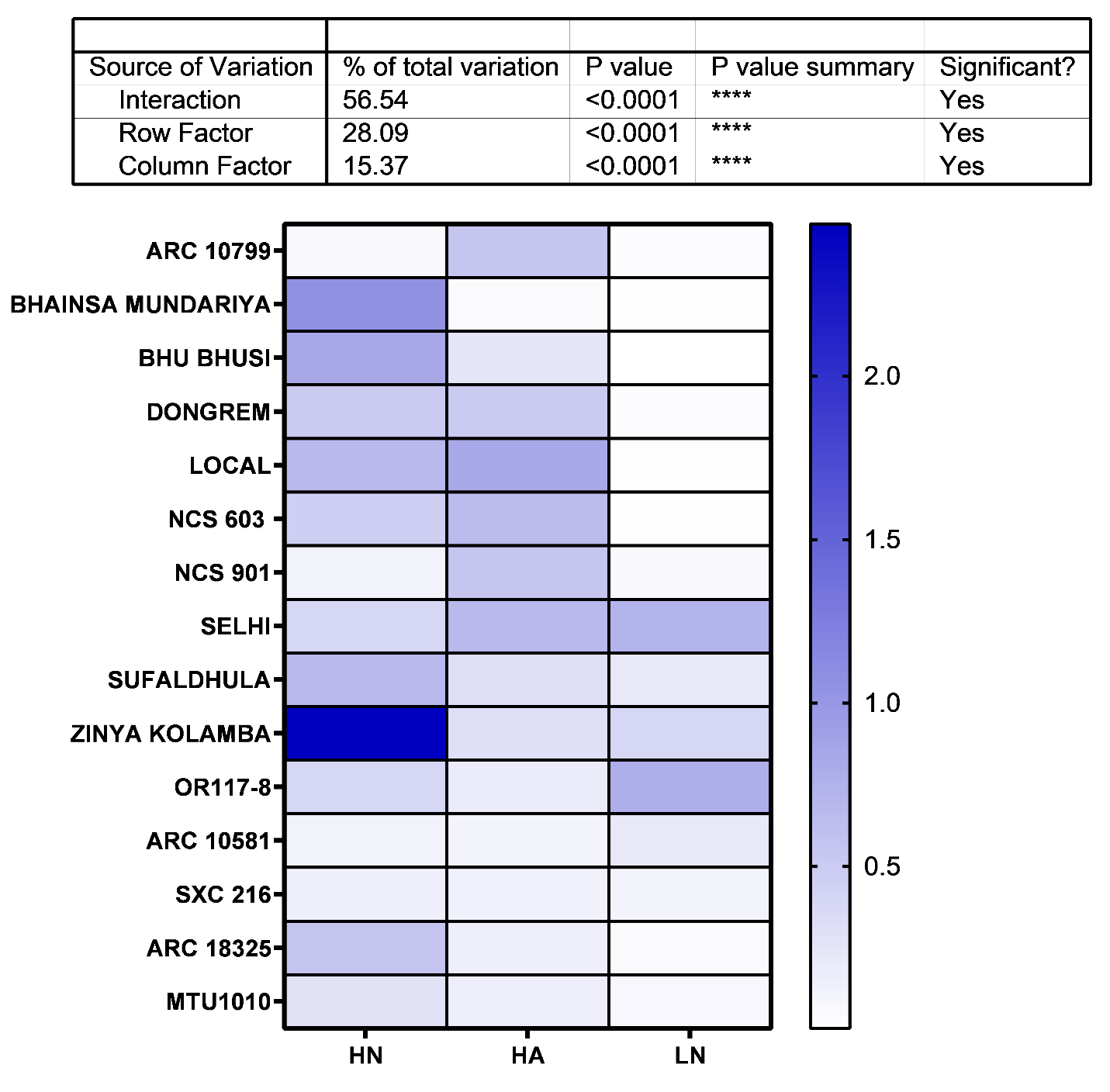
**Supplementary Figure 1. Effect of high nitrate (HN), high ammonium (HA) and nitrogen deficient (LN) conditions on A) total shoot N accumulation B) NUE of rice seedlings grown under hydroponic culture. (HN-7mM+Nitrate:0.5mM Ammonium, HA-7mM Ammonium+0.5mM Nitrate, LN-0.24mM Ammonium Nitrate)**

**A**

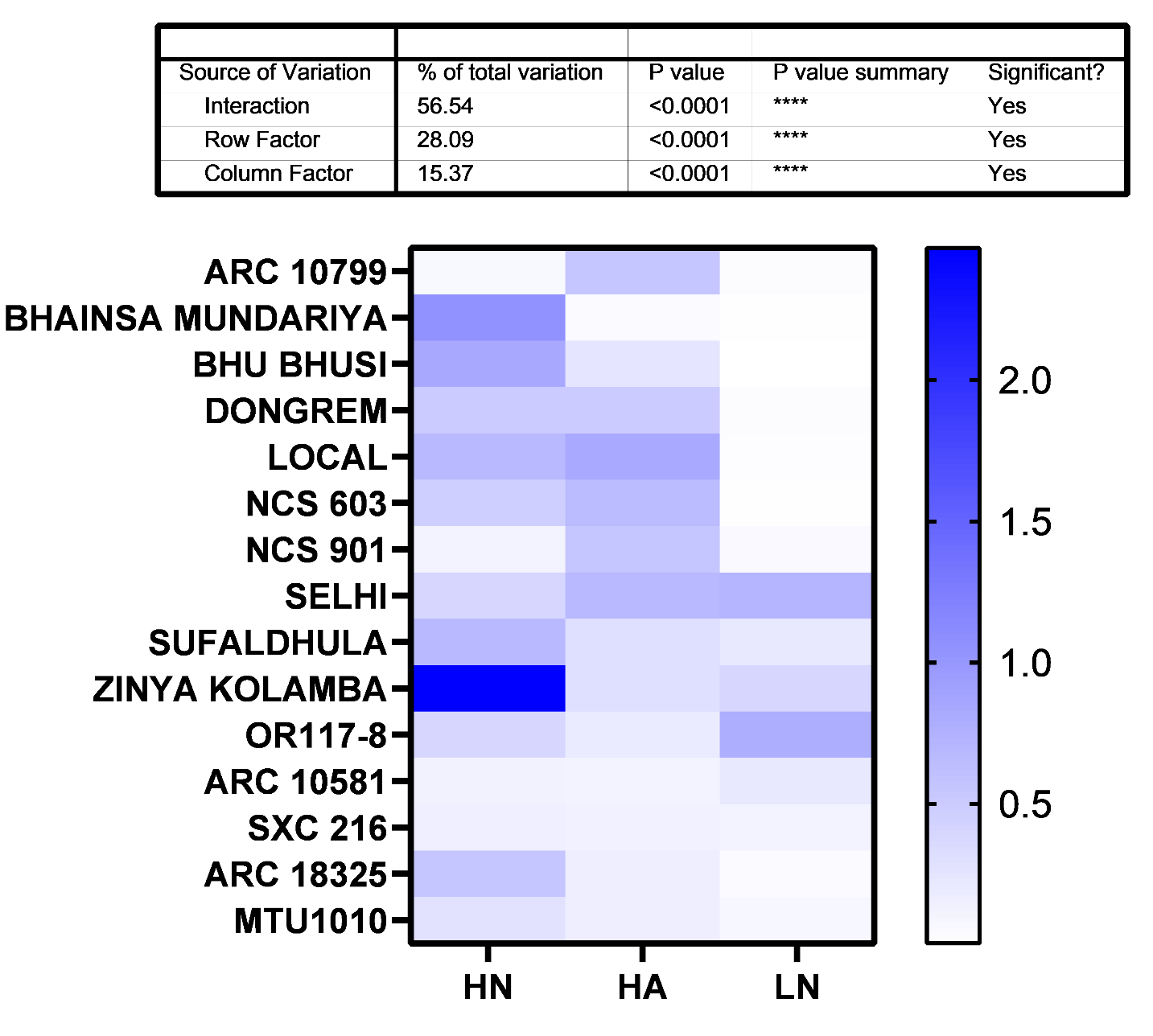

**B**

**Supplementary Figure 2. Growth of rice haplotypes in field under high N (N120) and low N (N0) conditions for the evaluation of morpho-physiological traits**

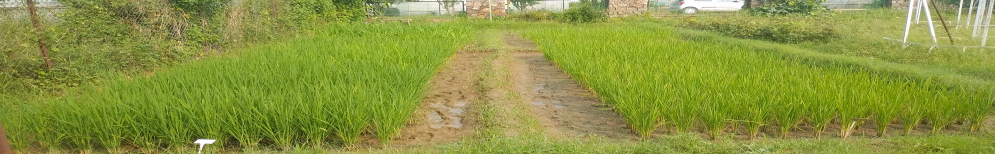

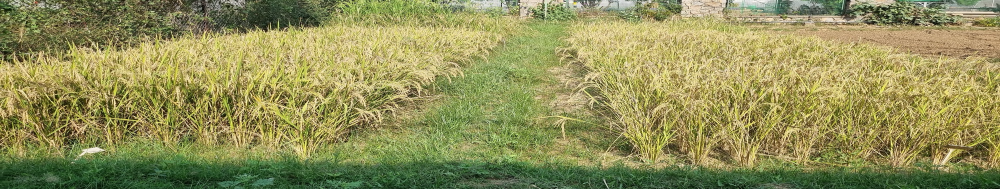

N+

N-

38 DAT

93 DAT

**Supplementary Figure 3 Thermal images and the normal images of the selected genotypes in field under N120 and N0 conditions which were analysed in Smartview 4.3 software**

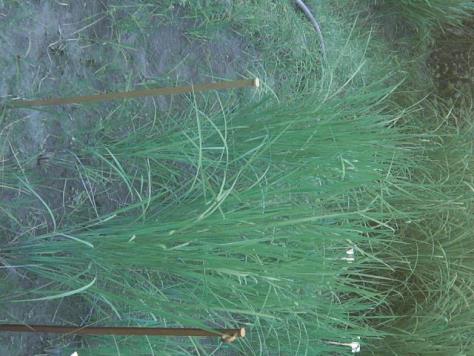

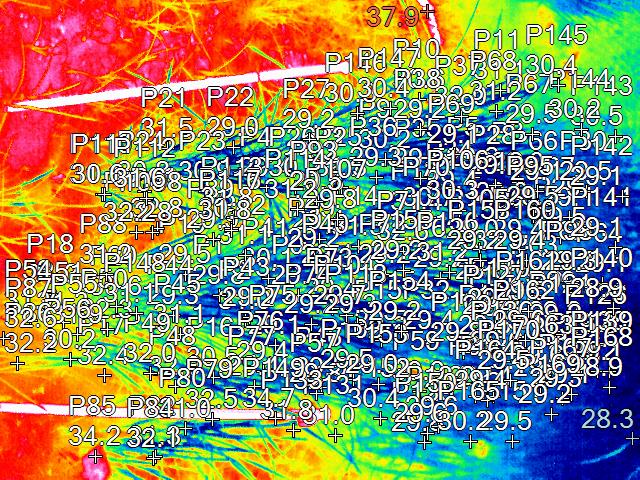

**ARC 10799 (N-)**

**ARC 10799(N-)**

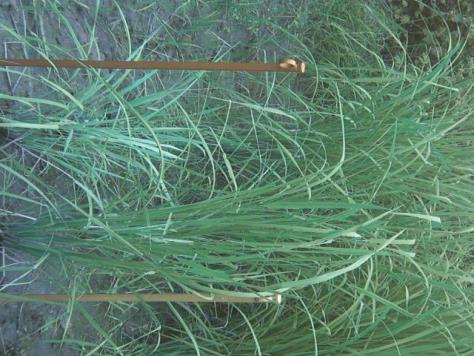

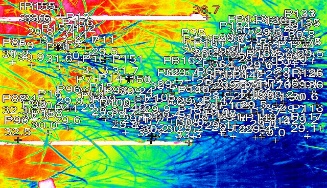

**ARC 10799 (N+)**

**ARC 10799 (N+)**

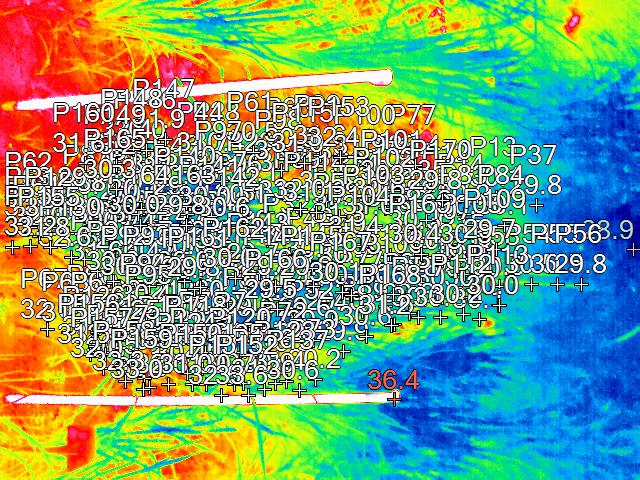

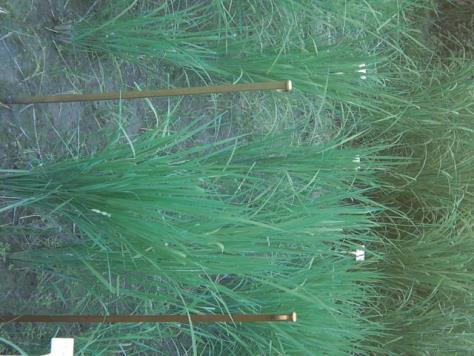

**ARC 18325 (N-)**

**ARC 18325 (N-)**

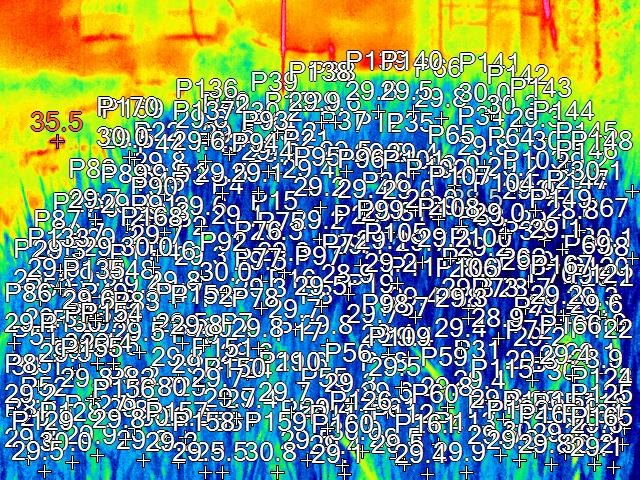

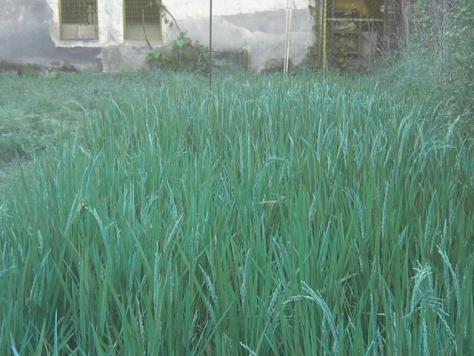

**MTU1010 (N+)**

**MTU1010 (N+)**

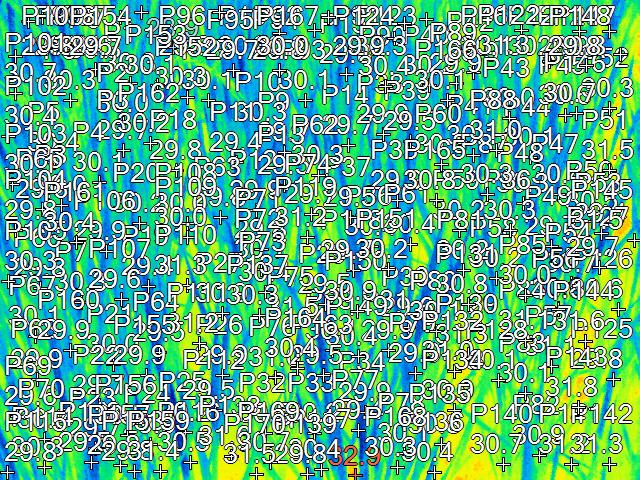

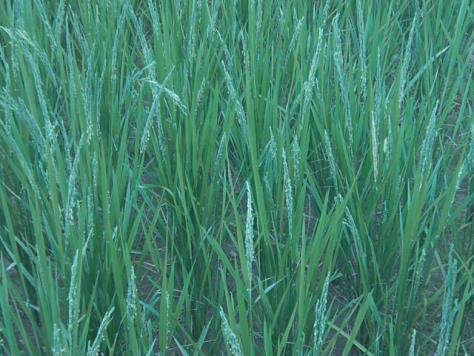

**MTU1010 (N-)**

**MTU1010 (N-)**

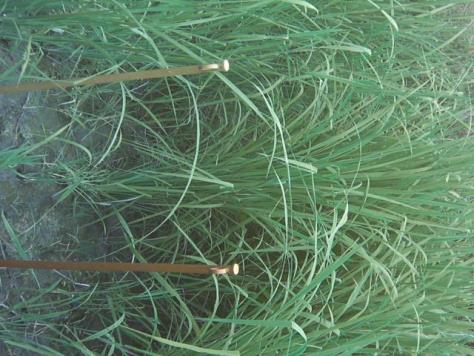

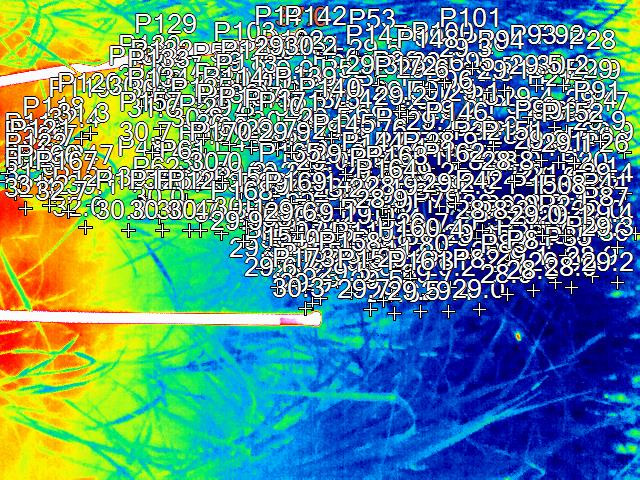

**OR 117-8 (+N)**

**OR 117-8 (+N)**

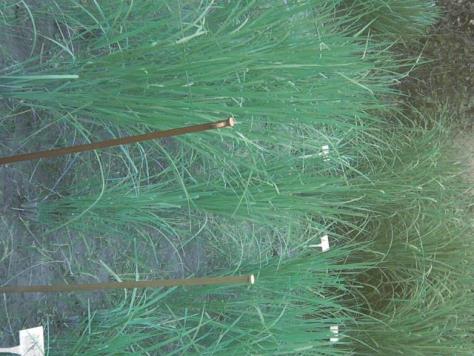

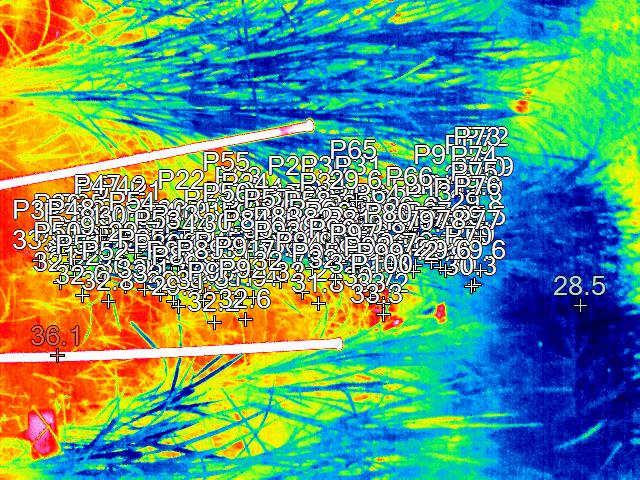

**OR 117-8 (-N)**

**OR 117-8 (-N)**

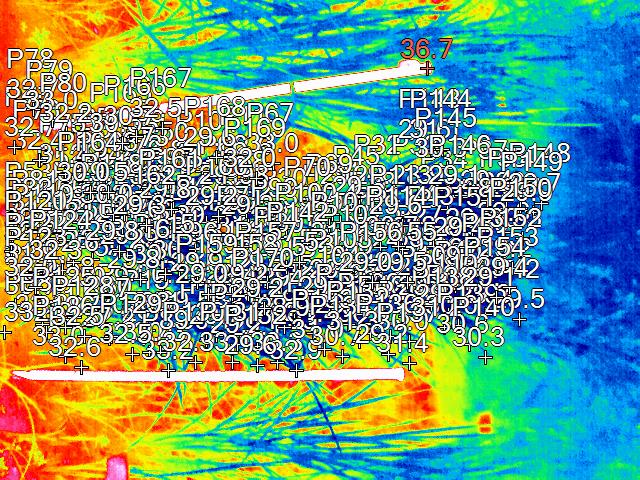

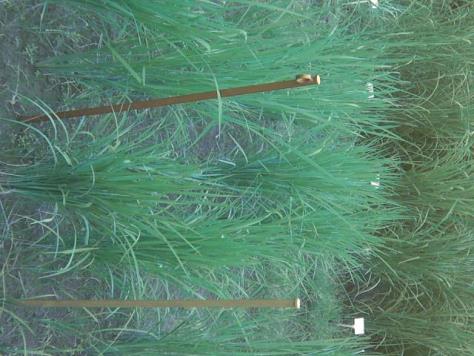

**SXC 216 (-N)**

**SXC 216 (-N)**

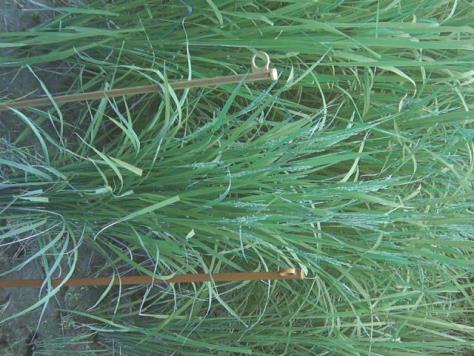

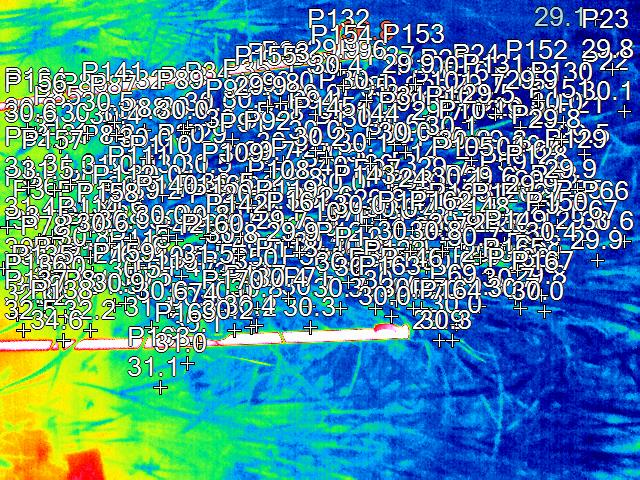

**SXC 216 (+N)**

**SXC 216 (+N)**

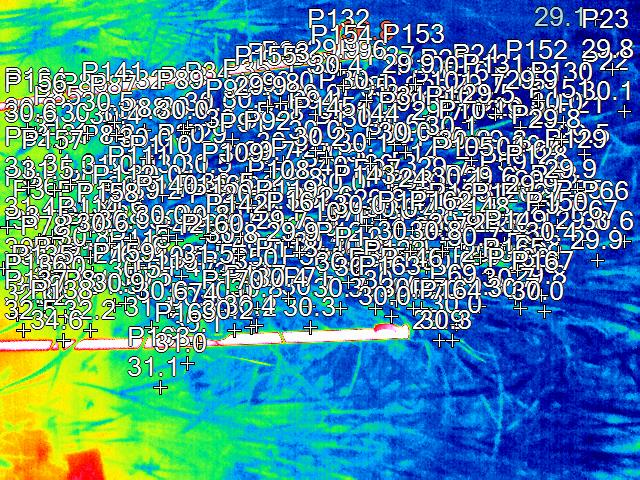

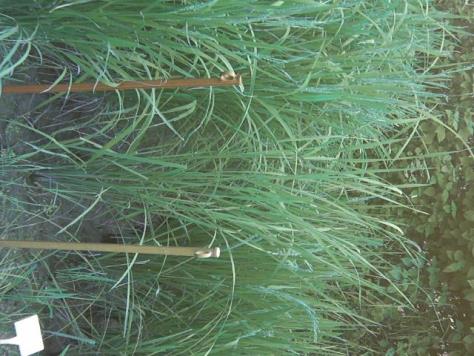

**ARC 18325(+N) (+N)**

**ARC 18325 (+N)**

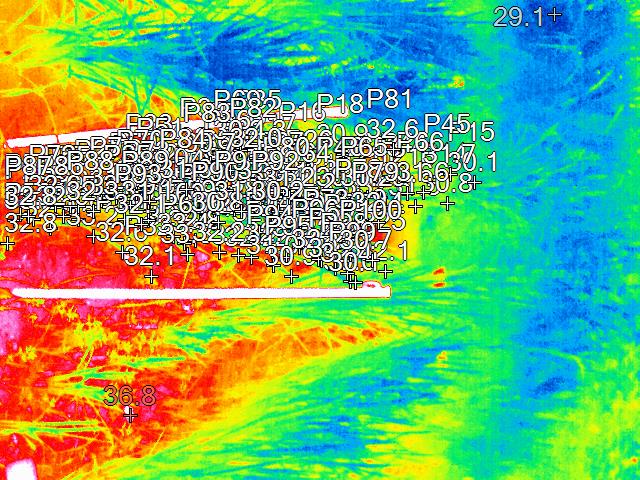

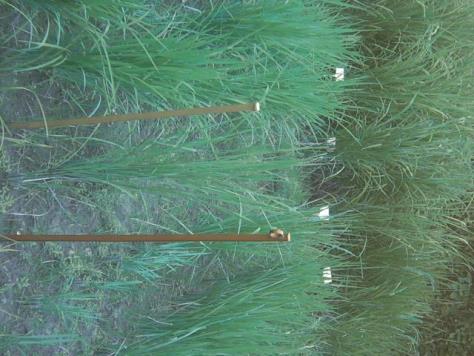

**ARC 10581 (-N)**

**ARC 10581 (-N)**

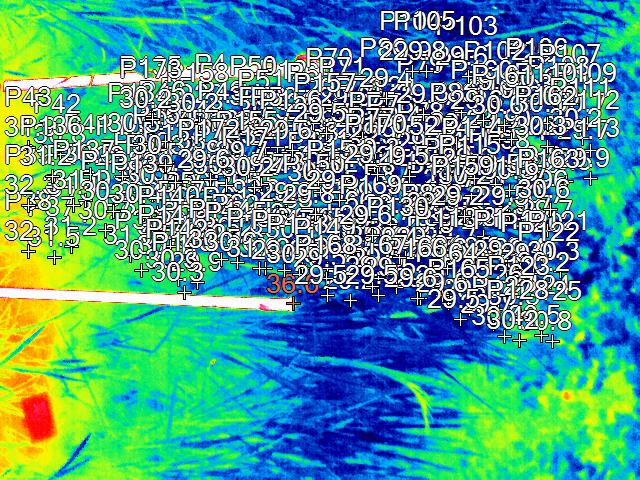

**ARC 10581 (+N)**

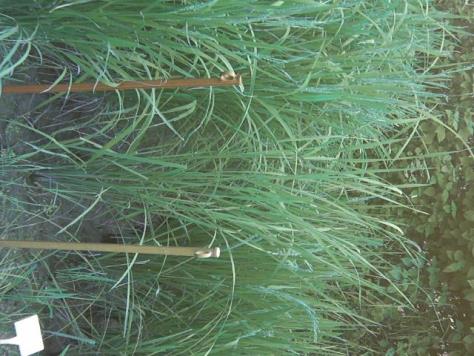

**ARC 10581 (+N)**

**Supplementary Figure 4. Comparison of rice haplotypes grown in field under N120 and N0 treatments (IRG-55: ARC 10799; IRG 242: SXC 216; IRG 318: OR117-8; IRG 383:ARC 18325; IRG 127: ARC 10581; MTU1010)**

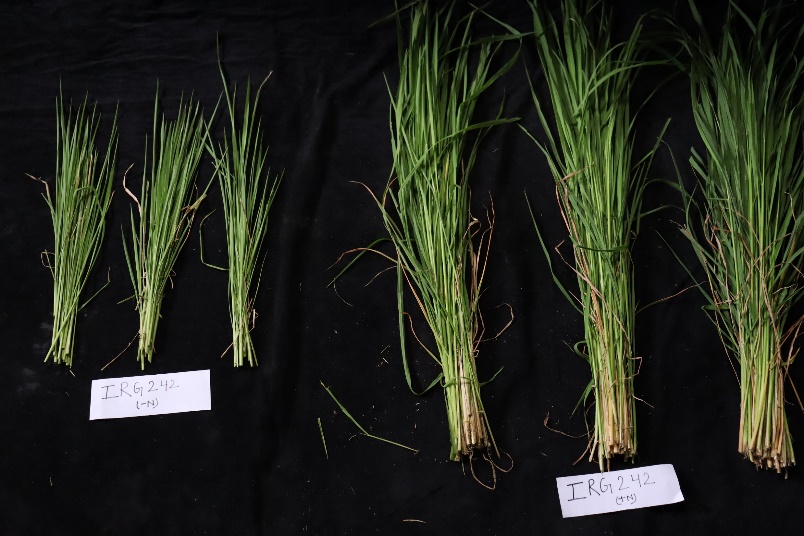

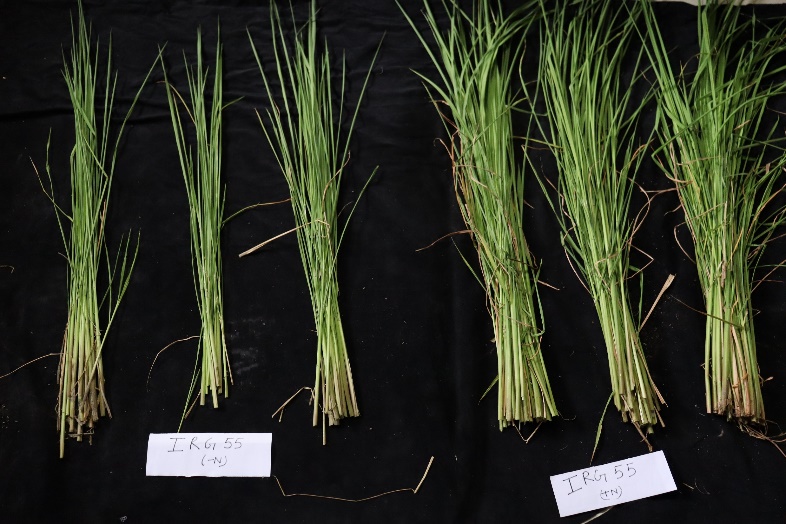

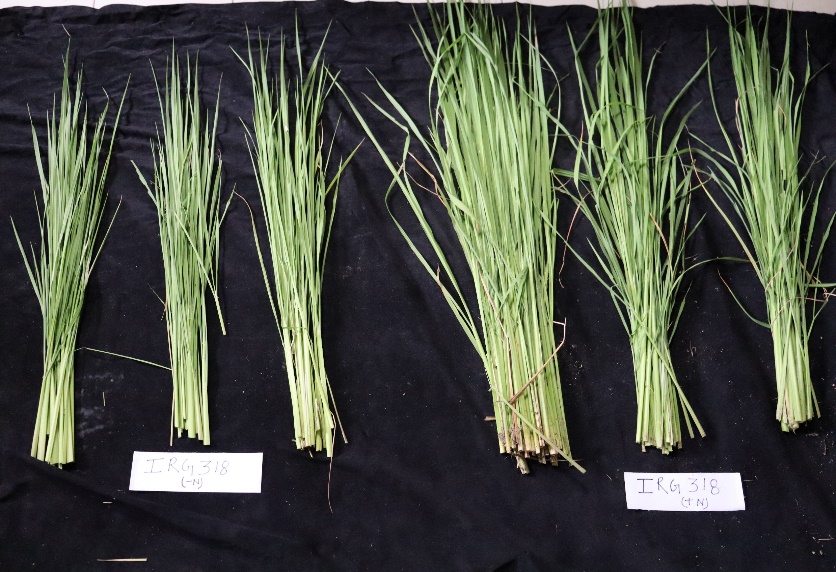

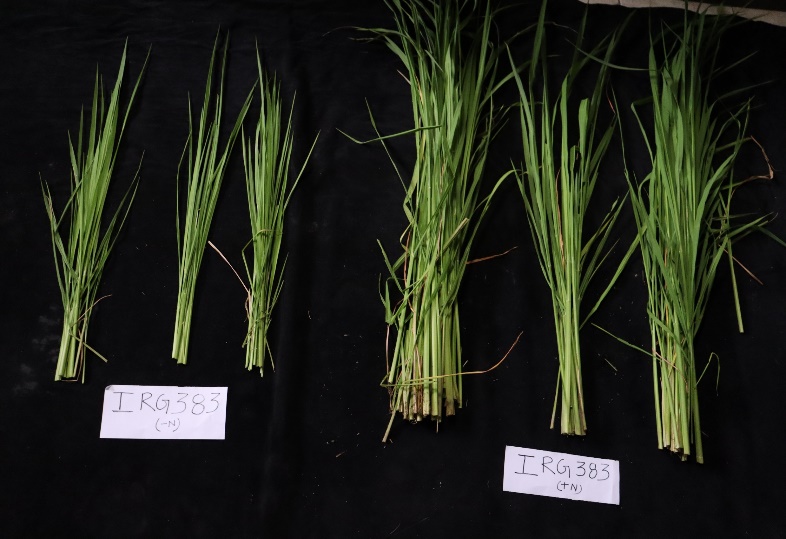

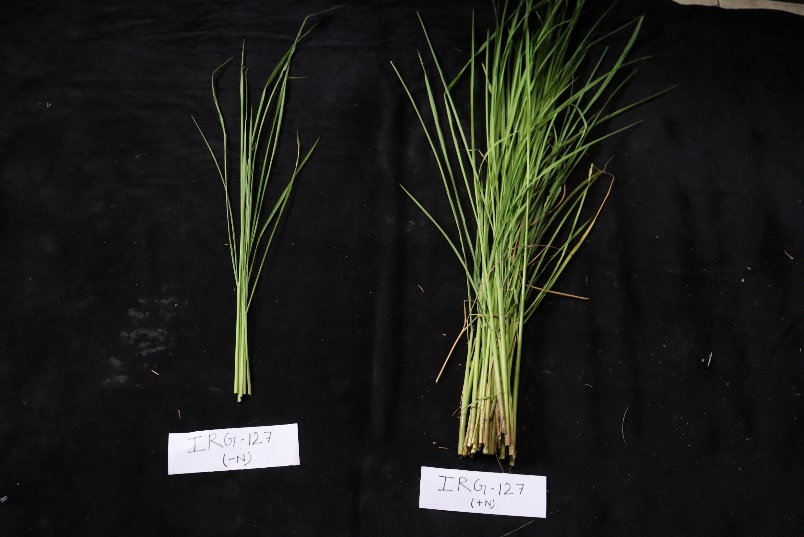

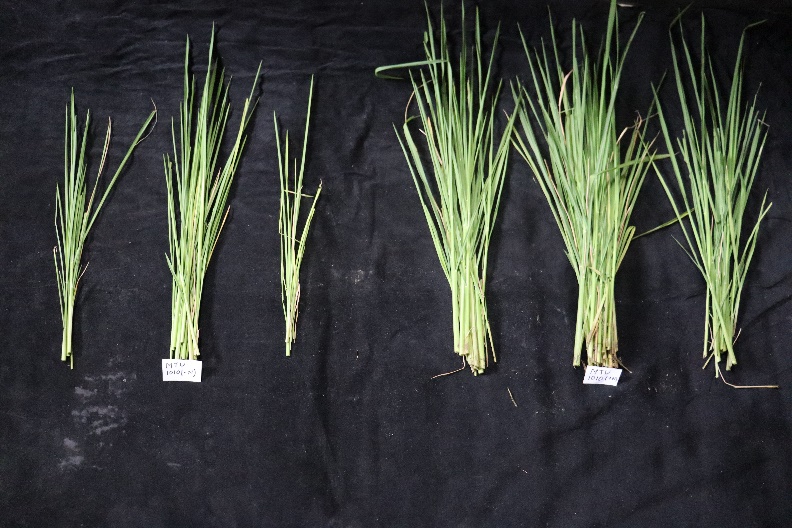

**Supplementary Figure 5. 3 P values of Pearson correlation matrix showing relationship between physiological and NUE parameters in rice genotypes belonging to different AAP3, AAP5 and AAP11 haplotypes and grown in high nitrogen (120 kg ha−1 applied N: N120), and nitrogen deficient (no applied N: N0) field conditions**

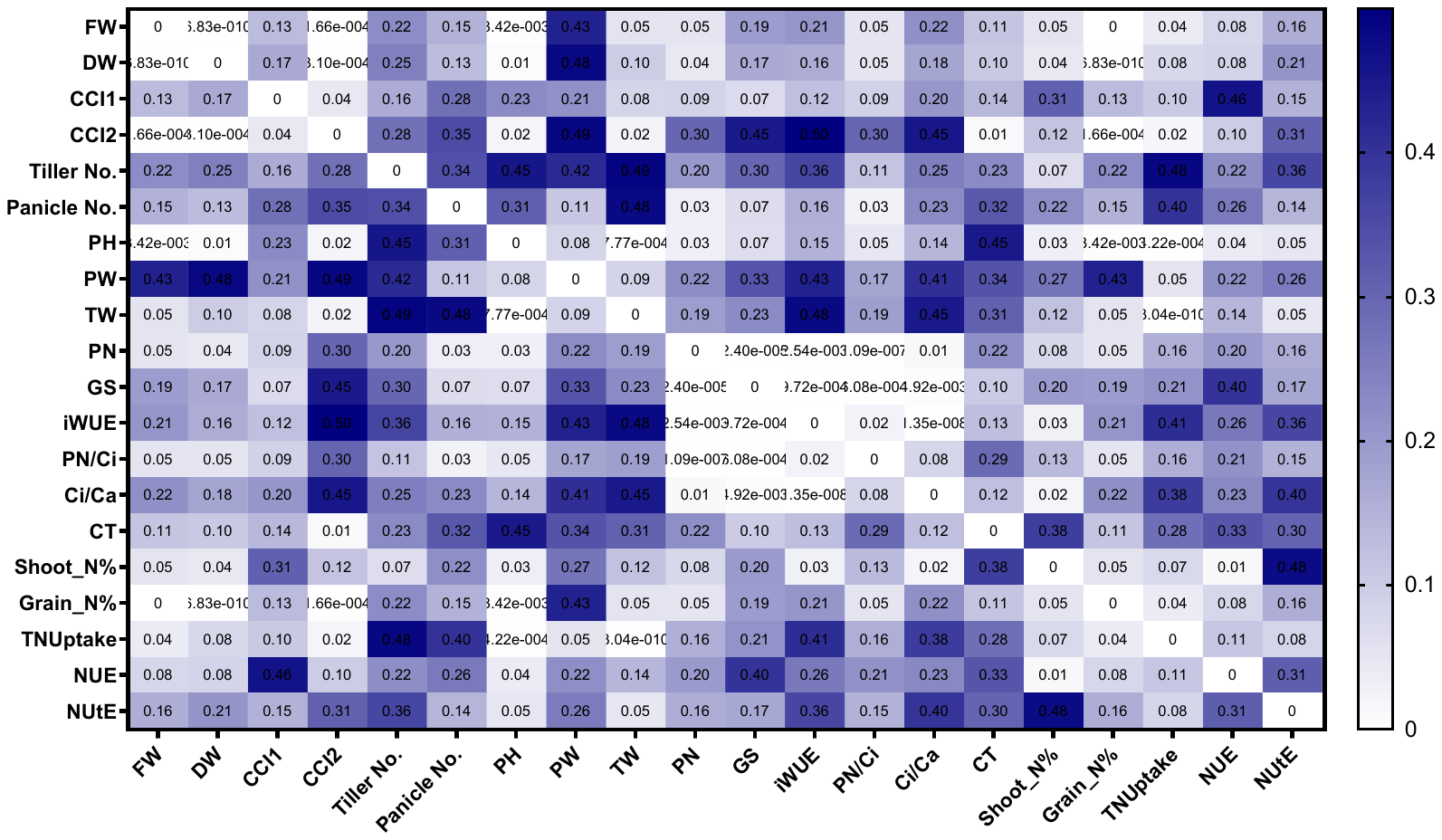
