## Supplementary material for "Association of haplotypes of AAP family amino acid transporters with nitrogen response and nitrogen use efficiency in rice grown under hydroponics and field conditions": 4.3supplementray table 2-6

**Supplementary table 2. Sidak’s multiple comparisons test on the effect of high nitrate (HN), high ammonium (HA) and nitrogen deficient (LN) conditions on (a) root biomass (RDW) and (b) shoot biomass (SDW) of rice seedlings grown under hydroponic culture.**

| Sidak's multiple comparisons test | Significant? | Summary | Adjusted P Value | Sidak's multiple comparisons test | Significant? | Summary | Adjusted P Value |
| --- | --- | --- | --- | --- | --- | --- | --- |
| RDW (g) |  |  |  | SDW (g) |  |  |  |
| ARC 10799 |  |  |  | ARC 10799 | |  |  |
| HA vs. HN | No | ns | 0.4692 | HA vs. HN | No | ns | >0.9999 |
| LN vs. HN | No | ns | 0.3719 | LN vs. HN | No | ns | 0.0864 |
| LN vs. HA | No | ns | 0.9981 | LN vs. HA | No | ns | 0.0956 |
| BHAINSA MUNDARIYA |  |  |  | BHAINSA MUNDARIYA | | |  |
| HA vs. HN | Yes | * | 0.0203 | HA vs. HN | No | ns | 0.1345 |
| LN vs. HN | Yes | ** | 0.003 | LN vs. HN | Yes | **** | <0.0001 |
| LN vs. HA | No | ns | 0.9005 | LN vs. HA | Yes | **** | <0.0001 |
| BHU BHUSI |  |  |  | BHU BHUSI | |  |  |
| HA vs. HN | No | ns | 0.9545 | HA vs. HN | No | ns | 0.9597 |
| LN vs. HN | No | ns | 0.6537 | LN vs. HN | Yes | ** | 0.0028 |
| LN vs. HA | No | ns | 0.9162 | LN vs. HA | Yes | * | 0.0116 |
| DONGREM |  |  |  | DONGREM | |  |  |
| HA vs. HN | No | ns | 0.6272 | HA vs. HN | No | ns | 0.9998 |
| LN vs. HN | No | ns | 0.9992 | LN vs. HN | No | ns | 0.0981 |
| LN vs. HA | No | ns | 0.7057 | LN vs. HA | No | ns | 0.1151 |
| LOCAL |  |  |  | LOCAL |  |  |  |
| HA vs. HN | No | ns | 0.1984 | HA vs. HN | No | ns | 0.4571 |
| LN vs. HN | Yes | ** | 0.001 | LN vs. HN | Yes | **** | <0.0001 |
| LN vs. HA | No | ns | 0.1834 | LN vs. HA | Yes | **** | <0.0001 |
| NCS 603 |  |  |  | NCS 603 |  |  |  |
| HA vs. HN | No | ns | 0.731 | HA vs. HN | No | ns | 0.9989 |
| LN vs. HN | No | ns | 0.3719 | LN vs. HN | Yes | ** | 0.0016 |
| LN vs. HA | No | ns | 0.9304 | LN vs. HA | Yes | ** | 0.001 |
| NCS 901 |  |  |  | NCS 901 |  |  |  |
| HA vs. HN | Yes | **** | <0.0001 | HA vs. HN | No | ns | 0.0933 |
| LN vs. HN | Yes | ** | 0.0017 | LN vs. HN | Yes | **** | <0.0001 |
| LN vs. HA | No | ns | 0.0637 | LN vs. HA | Yes | **** | <0.0001 |
| SELHI |  |  |  | SELHI |  |  |  |
| HA vs. HN | No | ns | 0.3274 | HA vs. HN | No | ns | 0.9018 |
| LN vs. HN | No | ns | 0.9981 | LN vs. HN | No | ns | 0.6574 |
| LN vs. HA | No | ns | 0.4193 | LN vs. HA | No | ns | 0.9649 |
| SUFALDHULA |  |  |  | SUFALDHULA | |  |  |
| HA vs. HN | No | ns | 0.2862 | HA vs. HN | No | ns | 0.7183 |
| LN vs. HN | No | ns | 0.9799 | LN vs. HN | Yes | *** | 0.0002 |
| LN vs. HA | No | ns | 0.4949 | LN vs. HA | Yes | ** | 0.0059 |
| ZINYA KOLAMBA |  |  |  | ZINYA KOLAMBA | |  |  |
| HA vs. HN | No | ns | 0.9432 | HA vs. HN | No | ns | 0.904 |
| LN vs. HN | No | ns | 0.9981 | LN vs. HN | No | ns | 0.0864 |
| LN vs. HA | No | ns | 0.9799 | LN vs. HA | Yes | * | 0.017 |
| OR117-8 |  |  |  | OR117-8 |  |  |  |
| HA vs. HN | No | ns | 0.9964 | HA vs. HN | No | ns | 0.617 |
| LN vs. HN | No | ns | 0.9432 | LN vs. HN | Yes | * | 0.0202 |
| LN vs. HA | No | ns | 0.9857 | LN vs. HA | Yes | *** | 0.0005 |
| ARC 10581 |  |  |  | ARC 10581 | |  |  |
| HA vs. HN | No | ns | 0.5474 | HA vs. HN | No | ns | 0.2309 |
| LN vs. HN | No | ns | 0.6272 | LN vs. HN | Yes | ** | 0.0042 |
| LN vs. HA | No | ns | 0.9992 | LN vs. HA | No | ns | 0.3421 |
| SXC 216 |  |  |  | SXC 216 |  |  |  |
| HA vs. HN | No | ns | 0.8245 | HA vs. HN | No | ns | 0.9979 |
| LN vs. HN | No | ns | 0.9903 | LN vs. HN | Yes | *** | 0.0008 |
| LN vs. HA | No | ns | 0.6537 | LN vs. HA | Yes | ** | 0.0014 |
| ARC 18325 |  |  |  | ARC 18325 | |  |  |
| HA vs. HN | No | ns | 0.9162 | HA vs. HN | No | ns | 0.279 |
| LN vs. HN | No | ns | 0.132 | LN vs. HN | Yes | ** | 0.0011 |
| LN vs. HA | No | ns | 0.3952 | LN vs. HA | Yes | **** | <0.0001 |
| MTU1010 |  |  |  | MTU1010 |  |  |  |
| HA vs. HN | Yes | **** | <0.0001 | HA vs. HN | No | ns | 0.9148 |
| LN vs. HN | Yes | **** | <0.0001 | LN vs. HN | No | ns | 0.0574 |
| LN vs. HA | No | ns | 0.9304 | LN vs. HA | No | ns | 0.2147 |

**Supplementary table 3. Sidak’s multiple comparisons test on the effect of high nitrate (HN), high ammonium (HA) and nitrogen deficient (LN) conditions on (a) leaf area (LA) (b) plant height (PH) of rice seedlings grown under hydroponic culture.**

| a) LA (cm^2^) | | | | b) PH (cm) | | | |
| --- | --- | --- | --- | --- | --- | --- | --- |
| Sidak's multiple comparisons test | Significant? | Summary | Adjusted P Value | Sidak's multiple comparisons test | Significant? | Summary | Adjusted P Value |
| ARC 10799 | |  |  | ARC 10799 | |  |  |
| HA vs. HN | No | ns | 0.9883 | HA vs. HN | No | ns | 0.9213 |
| LN vs. HN | No | ns | 0.4204 | LN vs. HN | No | ns | 0.1109 |
| LN vs. HA | No | ns | 0.6108 | LN vs. HA | No | ns | 0.3386 |
| BHAINSA MUNDARIYA | | |  | BHAINSA MUNDARIYA | | |  |
| HA vs. HN | No | ns | 0.7405 | HA vs. HN | No | ns | 0.3386 |
| LN vs. HN | Yes | **** | <0.0001 | LN vs. HN | No | ns | 0.0564 |
| LN vs. HA | Yes | *** | 0.0006 | LN vs. HA | Yes | *** | 0.0005 |
| BHU BHUSI | |  |  | BHU BHUSI | |  |  |
| HA vs. HN | No | ns | 0.8535 | HA vs. HN | No | ns | 0.9975 |
| LN vs. HN | No | ns | 0.163 | LN vs. HN | Yes | **** | <0.0001 |
| LN vs. HA | No | ns | 0.5382 | LN vs. HA | Yes | **** | <0.0001 |
| DONGREM | |  |  | DONGREM | |  |  |
| HA vs. HN | No | ns | 0.9182 | HA vs. HN | No | ns | 0.9708 |
| LN vs. HN | No | ns | 0.4482 | LN vs. HN | No | ns | 0.1804 |
| LN vs. HA | No | ns | 0.8215 | LN vs. HA | No | ns | 0.0746 |
| LOCAL |  |  |  | LOCAL |  |  |  |
| HA vs. HN | No | ns | 0.1857 | HA vs. HN | No | ns | 0.9708 |
| LN vs. HN | Yes | **** | <0.0001 | LN vs. HN | No | ns | 0.8445 |
| LN vs. HA | Yes | **** | <0.0001 | LN vs. HA | No | ns | 0.5932 |
| NCS 603 |  |  |  | NCS 603 |  |  |  |
| HA vs. HN | No | ns | 0.999 | HA vs. HN | No | ns | 0.9919 |
| LN vs. HN | Yes | * | 0.0238 | LN vs. HN | Yes | *** | 0.0006 |
| LN vs. HA | Yes | * | 0.0328 | LN vs. HA | Yes | ** | 0.0014 |
| NCS 901 |  |  |  | NCS 901 |  |  |  |
| HA vs. HN | Yes | * | 0.0271 | HA vs. HN | Yes | * | 0.0167 |
| LN vs. HN | Yes | **** | <0.0001 | LN vs. HN | Yes | * | 0.0489 |
| LN vs. HA | Yes | * | 0.0451 | LN vs. HA | No | ns | 0.9708 |
| SELHI |  |  |  | SELHI |  |  |  |
| HA vs. HN | No | ns | 0.535 | HA vs. HN | Yes | **** | <0.0001 |
| LN vs. HN | No | ns | 0.5239 | LN vs. HN | Yes | ** | 0.0029 |
| LN vs. HA | No | ns | >0.9999 | LN vs. HA | No | ns | 0.3712 |
| SUFALDHULA | |  |  | SUFALDHULA | |  |  |
| HA vs. HN | No | ns | 0.996 | HA vs. HN | No | ns | 0.9708 |
| LN vs. HN | No | ns | 0.7522 | LN vs. HN | No | ns | 0.2257 |
| LN vs. HA | No | ns | 0.8651 | LN vs. HA | No | ns | 0.4412 |
| ZINYA KOLAMBA | |  |  | ZINYA KOLAMBA | |  |  |
| HA vs. HN | No | ns | 0.9809 | HA vs. HN | Yes | * | 0.0142 |
| LN vs. HN | No | ns | 0.6891 | LN vs. HN | Yes | ** | 0.0017 |
| LN vs. HA | No | ns | 0.8858 | LN vs. HA | No | ns | 0.873 |
| OR117-8 |  |  |  | OR117-8 |  |  |  |
| HA vs. HN | Yes | * | 0.0383 | HA vs. HN | No | ns | 0.9708 |
| LN vs. HN | Yes | ** | 0.0034 | LN vs. HN | Yes | *** | 0.0005 |
| LN vs. HA | No | ns | 0.7592 | LN vs. HA | Yes | *** | 0.0001 |
| ARC 10581 | |  |  | ARC 10581 | |  |  |
| HA vs. HN | No | ns | 0.086 | HA vs. HN | No | ns | 0.4412 |
| LN vs. HN | Yes | ** | 0.0071 | LN vs. HN | Yes | **** | <0.0001 |
| LN vs. HA | No | ns | 0.7058 | LN vs. HA | Yes | *** | 0.0002 |
| SXC 216 |  |  |  | SXC 216 |  |  |  |
| HA vs. HN | No | ns | 0.8264 | HA vs. HN | No | ns | 0.8135 |
| LN vs. HN | Yes | * | 0.019 | LN vs. HN | Yes | *** | 0.0001 |
| LN vs. HA | No | ns | 0.1227 | LN vs. HA | Yes | ** | 0.002 |
| ARC 18325 | |  |  | ARC 18325 | |  |  |
| HA vs. HN | No | ns | 0.9998 | HA vs. HN | Yes | * | 0.0313 |
| LN vs. HN | Yes | *** | 0.0003 | LN vs. HN | Yes | **** | <0.0001 |
| LN vs. HA | Yes | *** | 0.0003 | LN vs. HA | Yes | ** | 0.0014 |
| MTU1010 |  |  |  | MTU1010 |  |  |  |
| HA vs. HN | Yes | * | 0.015 | HA vs. HN | No | ns | 0.1804 |
| LN vs. HN | Yes | *** | 0.0006 | LN vs. HN | Yes | ** | 0.0029 |
| LN vs. HA | No | ns | 0.6293 | LN vs. HA | No | ns | 0.3386 |

**Supplementary table 4. Sidak’s multiple comparison test on the effect of high nitrate (HN), high ammonium (HA) and nitrogen deficient (LN) conditions on chlorophyll content index (CCI) and tissue NO_3_^-^ (TN)**

| Sidak's multiple comparisons test | Significant? | Summary | Adjusted P Value | Sidak's multiple comparisons test | Significant? | Summary | Adjusted P Value |
| --- | --- | --- | --- | --- | --- | --- | --- |
| CCI | | | | Tissue NO_3_^-^ (TN) |  |  |  |
| ARC 10799 |  |  |  |  |  |  |  |
| HA vs. HN | No | ns | 0.2941 | ARC 10799 |  |  |  |
| LN vs. HN | Yes | ** | 0.0053 | HA vs. HN | Yes | **** | <0.0001 |
| LN vs. HA | No | ns | 0.3233 | LN vs. HN | Yes | **** | <0.0001 |
|  |  |  |  | LN vs. HA | Yes | *** | 0.0002 |
| BHAINSA MUNDARIYA | | |  |  |  |  |  |
| HA vs. HN | No | ns | 0.9626 | BHAINSA MUNDARIYA |  |  |  |
| LN vs. HN | Yes | **** | <0.0001 | HA vs. HN | Yes | **** | <0.0001 |
| LN vs. HA | Yes | **** | <0.0001 | LN vs. HN | Yes | **** | <0.0001 |
|  |  |  |  | LN vs. HA | Yes | **** | <0.0001 |
| BHU BHUSI | |  |  |  |  |  |  |
| HA vs. HN | No | ns | 0.9029 | BHU BHUSI |  |  |  |
| LN vs. HN | No | ns | 0.3754 | HA vs. HN | Yes | *** | 0.0005 |
| LN vs. HA | No | ns | 0.1124 | LN vs. HN | Yes | **** | <0.0001 |
|  |  |  |  | LN vs. HA | No | ns | 0.1697 |
| DONGREM | |  |  |  |  |  |  |
| HA vs. HN | No | ns | 0.759 | DONGREM |  |  |  |
| LN vs. HN | Yes | ** | 0.002 | HA vs. HN | Yes | **** | <0.0001 |
| LN vs. HA | Yes | * | 0.0324 | LN vs. HN | Yes | **** | <0.0001 |
|  |  |  |  | LN vs. HA | Yes | * | 0.0417 |
| LOCAL |  |  |  |  |  |  |  |
| HA vs. HN | No | ns | 0.0636 | LOCAL |  |  |  |
| LN vs. HN | No | ns | 0.1033 | HA vs. HN | Yes | **** | <0.0001 |
| LN vs. HA | Yes | **** | <0.0001 | LN vs. HN | Yes | **** | <0.0001 |
|  |  |  |  | LN vs. HA | Yes | **** | <0.0001 |
| NCS 603 |  |  |  |  |  |  |  |
| HA vs. HN | No | ns | 0.9985 | NCS 603 |  |  |  |
| LN vs. HN | Yes | * | 0.0324 | HA vs. HN | Yes | **** | <0.0001 |
| LN vs. HA | Yes | * | 0.048 | LN vs. HN | Yes | **** | <0.0001 |
|  |  |  |  | LN vs. HA | Yes | *** | 0.0004 |
| NCS 901 |  |  |  |  |  |  |  |
| HA vs. HN | Yes | **** | <0.0001 | NCS 901 |  |  |  |
| LN vs. HN | Yes | **** | <0.0001 | HA vs. HN | Yes | **** | <0.0001 |
| LN vs. HA | No | ns | 0.503 | LN vs. HN | Yes | **** | <0.0001 |
|  |  |  |  | LN vs. HA | No | ns | 0.0875 |
| SELHI |  |  |  |  |  |  |  |
| HA vs. HN | No | ns | 0.9744 | SELHI |  |  |  |
| LN vs. HN | No | ns | 0.7361 | HA vs. HN | Yes | **** | <0.0001 |
| LN vs. HA | No | ns | 0.9306 | LN vs. HN | Yes | **** | <0.0001 |
|  |  |  |  | LN vs. HA | Yes | * | 0.0193 |
| SUFALDHULA | |  |  |  |  |  |  |
| HA vs. HN | No | ns | 0.6022 | SUFALDHULA |  |  |  |
| LN vs. HN | No | ns | 0.3134 | HA vs. HN | Yes | **** | <0.0001 |
| LN vs. HA | No | ns | 0.958 | LN vs. HN | Yes | **** | <0.0001 |
|  |  |  |  | LN vs. HA | No | ns | 0.1313 |
| ZINYA KOLAMBA | |  |  |  |  |  |  |
| HA vs. HN | No | ns | 0.759 | ZINYA KOLAMBA |  |  |  |
| LN vs. HN | No | ns | 0.2666 | HA vs. HN | Yes | **** | <0.0001 |
| LN vs. HA | No | ns | 0.8233 | LN vs. HN | Yes | **** | <0.0001 |
|  |  |  |  | LN vs. HA | Yes | ** | 0.0026 |
| OR117-8 |  |  |  |  |  |  |  |
| HA vs. HN | No | ns | 0.9808 | OR117-8 |  |  |  |
| LN vs. HN | Yes | * | 0.0204 | HA vs. HN | Yes | **** | <0.0001 |
| LN vs. HA | No | ns | 0.0528 | LN vs. HN | Yes | **** | <0.0001 |
|  |  |  |  | LN vs. HA | No | ns | 0.1348 |
| ARC 10581 | |  |  |  |  |  |  |
| HA vs. HN | No | ns | 0.9532 | ARC 10581 |  |  |  |
| LN vs. HN | No | ns | 0.3647 | HA vs. HN | Yes | **** | <0.0001 |
| LN vs. HA | No | ns | 0.1492 | LN vs. HN | Yes | **** | <0.0001 |
|  |  |  |  | LN vs. HA | No | ns | 0.0617 |
| SXC 216 |  |  |  |  |  |  |  |
| HA vs. HN | No | ns | 0.9668 | SXC 216 |  |  |  |
| LN vs. HN | Yes | *** | 0.0002 | HA vs. HN | Yes | **** | <0.0001 |
| LN vs. HA | Yes | ** | 0.001 | LN vs. HN | Yes | **** | <0.0001 |
|  |  |  |  | LN vs. HA | Yes | * | 0.0243 |
| ARC 18325 | |  |  |  |  |  |  |
| HA vs. HN | Yes | * | 0.0239 | ARC 18325 |  |  |  |
| LN vs. HN | No | ns | 0.2018 | HA vs. HN | Yes | **** | <0.0001 |
| LN vs. HA | Yes | **** | <0.0001 | LN vs. HN | Yes | **** | <0.0001 |
|  |  |  |  | LN vs. HA | No | ns | 0.9983 |
| MTU1010 |  |  |  |  |  |  |  |
| HA vs. HN | No | ns | 0.2578 | MTU1010 |  |  |  |
| LN vs. HN | Yes | *** | 0.0001 | HA vs. HN | Yes | **** | <0.0001 |
| LN vs. HA | Yes | * | 0.0359 | LN vs. HN | Yes | **** | <0.0001 |
|  |  |  |  | LN vs. HA | No | ns | 0.9973 |

**Supplementary table 5. Sidak’s multiple comparison test on the effect of high nitrate (HN), high ammonium (HA) and nitrogen deficient (LN) conditions on root traits such as root average diameter (RD), total root length (TRL), total root surface area (TRSA), total root volume (TRV), main root length -diameter >0.5 mm (MRL), main root volume - diameter >0.5 mm (MRV), main root surface area - diameter >0.5 mm (MRSA), lateral roots length - diameter ≤0.5 mm (LRL), lateral roots volume -diameter ≤0.5 mm(LRV), lateral roots surface area -diameter ≤0.5 mm (LRSA), number of root forks and root tips of rice seedlings grown under hydroponic culture.**

| RD (mm) |  |  |  | TRL (cm) |  |  |  |
| --- | --- | --- | --- | --- | --- | --- | --- |
| Sidak's multiple comparisons test | Significant? | Summary | Adjusted P Value | Sidak's multiple comparisons test | Significant? | Summary | Adjusted P Value |
| ARC 10799 | |  |  | ARC 10799 | |  |  |
| HA vs. HN | Yes | ** | 0.0011 | HA vs. HN | No | ns | 0.9988 |
| LN vs. HN | No | ns | 0.3525 | LN vs. HN | No | ns | 0.8637 |
| LN vs. HA | Yes | **** | <0.0001 | LN vs. HA | No | ns | 0.7906 |
| BHAINSA MUNDARIYA | | |  | BHAINSA MUNDARIYA | | |  |
| HA vs. HN | Yes | *** | 0.0005 | HA vs. HN | No | ns | 0.3906 |
| LN vs. HN | No | ns | 0.8797 | LN vs. HN | No | ns | 0.9672 |
| LN vs. HA | Yes | ** | 0.0038 | LN vs. HA | No | ns | 0.1898 |
| BHU BHUSI | |  |  | BHU BHUSI | |  |  |
| HA vs. HN | No | ns | 0.2856 | HA vs. HN | No | ns | 0.9949 |
| LN vs. HN | Yes | * | 0.0375 | LN vs. HN | No | ns | 0.9207 |
| LN vs. HA | No | ns | 0.7233 | LN vs. HA | No | ns | 0.9793 |
| DONGREM | |  |  | DONGREM | |  |  |
| HA vs. HN | No | ns | 0.5835 | HA vs. HN | Yes | * | 0.0488 |
| LN vs. HN | No | ns | 0.9937 | LN vs. HN | Yes | ** | 0.0042 |
| LN vs. HA | No | ns | 0.7396 | LN vs. HA | No | ns | 0.7428 |
| LOCAL |  |  |  | LOCAL |  |  |  |
| HA vs. HN | Yes | **** | <0.0001 | HA vs. HN | No | ns | 0.9833 |
| LN vs. HN | No | ns | 0.8043 | LN vs. HN | Yes | ** | 0.0025 |
| LN vs. HA | Yes | **** | <0.0001 | LN vs. HA | Yes | ** | 0.0065 |
| NCS 603 |  |  |  | NCS 603 |  |  |  |
| HA vs. HN | Yes | **** | <0.0001 | HA vs. HN | No | ns | 0.4877 |
| LN vs. HN | Yes | ** | 0.002 | LN vs. HN | No | ns | 0.1454 |
| LN vs. HA | No | ns | 0.3696 | LN vs. HA | No | ns | 0.8636 |
| NCS 901 |  |  |  | NCS 901 |  |  |  |
| HA vs. HN | Yes | **** | <0.0001 | HA vs. HN | No | ns | 0.3294 |
| LN vs. HN | Yes | ** | 0.0017 | LN vs. HN | Yes | * | 0.0443 |
| LN vs. HA | Yes | **** | <0.0001 | LN vs. HA | Yes | *** | 0.0005 |
| SELHI |  |  |  | SELHI |  |  |  |
| HA vs. HN | No | ns | 0.8095 | HA vs. HN | No | ns | 0.3683 |
| LN vs. HN | Yes | * | 0.0222 | LN vs. HN | No | ns | 0.7234 |
| LN vs. HA | No | ns | 0.1479 | LN vs. HA | No | ns | 0.9295 |
| SUFALDHULA | |  |  | SUFALDHULA | |  |  |
| HA vs. HN | No | ns | 0.2938 | HA vs. HN | No | ns | 0.62 |
| LN vs. HN | Yes | * | 0.0287 | LN vs. HN | No | ns | 0.958 |
| LN vs. HA | No | ns | 0.6427 | LN vs. HA | No | ns | 0.8886 |
| ZINYA KOLAMBA | |  |  | ZINYA KOLAMBA | |  |  |
| HA vs. HN | No | ns | 0.133 | HA vs. HN | No | ns | 0.8521 |
| LN vs. HN | No | ns | >0.9999 | LN vs. HN | No | ns | 0.3334 |
| LN vs. HA | No | ns | 0.1498 | LN vs. HA | No | ns | 0.7954 |
| OR117-8 |  |  |  | OR117-8 |  |  |  |
| HA vs. HN | No | ns | 0.5631 | HA vs. HN | No | ns | 0.6245 |
| LN vs. HN | Yes | *** | 0.0007 | LN vs. HN | No | ns | 0.9265 |
| LN vs. HA | Yes | **** | <0.0001 | LN vs. HA | No | ns | 0.9317 |
| ARC 10581 | |  |  | ARC 10581 | |  |  |
| HA vs. HN | Yes | ** | 0.0024 | HA vs. HN | No | ns | 0.9994 |
| LN vs. HN | Yes | ** | 0.0018 | LN vs. HN | No | ns | 0.9386 |
| LN vs. HA | No | ns | 0.9997 | LN vs. HA | No | ns | 0.8997 |
| SXC 216 |  |  |  | SXC 216 |  |  |  |
| HA vs. HN | No | ns | 0.7301 | HA vs. HN | No | ns | 0.8701 |
| LN vs. HN | No | ns | 0.0515 | LN vs. HN | No | ns | 0.9669 |
| LN vs. HA | No | ns | 0.3482 | LN vs. HA | No | ns | 0.6176 |
| ARC 18325 | |  |  | ARC 18325 | |  |  |
| HA vs. HN | No | ns | 0.2282 | HA vs. HN | No | ns | 0.2377 |
| LN vs. HN | No | ns | 0.5835 | LN vs. HN | No | ns | 0.1432 |
| LN vs. HA | No | ns | 0.9028 | LN vs. HA | No | ns | 0.9915 |
| MTU1010 |  |  |  | MTU1010 |  |  |  |
| HA vs. HN | No | ns | 0.9108 | HA vs. HN | No | ns | 0.2063 |
| LN vs. HN | No | ns | 0.9994 | LN vs. HN | No | ns | 0.0524 |
| LN vs. HA | No | ns | 0.8639 | LN vs. HA | No | ns | 0.8966 |
| TRSA (cm^2^) |  |  |  | TRV (cm^3^) |  |  |  |
| Sidak's multiple comparisons test | Significant? | Summary | Adjusted P Value | Sidak's multiple comparisons test | Significant? | Summary | Adjusted P Value |
| ARC 10799 | |  |  | ARC 10799 | |  |  |
| HA vs. HN | No | ns | 0.8913 | HA vs. HN | No | ns | 0.6248 |
| LN vs. HN | No | ns | >0.9999 | LN vs. HN | No | ns | 0.9986 |
| LN vs. HA | No | ns | 0.8951 | LN vs. HA | No | ns | 0.5289 |
| BHAINSA MUNDARIYA | | |  | BHAINSA MUNDARIYA | | |  |
| HA vs. HN | No | ns | 0.3156 | HA vs. HN | No | ns | 0.4843 |
| LN vs. HN | No | ns | 0.0923 | LN vs. HN | Yes | ** | 0.01 |
| LN vs. HA | No | ns | 0.8992 | LN vs. HA | No | ns | 0.22 |
| BHU BHUSI | |  |  | BHU BHUSI | |  |  |
| HA vs. HN | No | ns | 0.5145 | HA vs. HN | No | ns | 0.1518 |
| LN vs. HN | No | ns | 0.0509 | LN vs. HN | Yes | * | 0.012 |
| LN vs. HA | No | ns | 0.5465 | LN vs. HA | No | ns | 0.6566 |
| DONGREM | |  |  | DONGREM | |  |  |
| HA vs. HN | No | ns | 0.9979 | HA vs. HN | No | ns | 0.987 |
| LN vs. HN | No | ns | 0.9981 | LN vs. HN | No | ns | 0.9988 |
| LN vs. HA | No | ns | >0.9999 | LN vs. HA | No | ns | 0.9977 |
| LOCAL |  |  |  | LOCAL |  |  |  |
| HA vs. HN | No | ns | 0.3714 | HA vs. HN | Yes | *** | 0.0006 |
| LN vs. HN | Yes | ** | 0.0031 | LN vs. HN | Yes | *** | 0.0006 |
| LN vs. HA | Yes | **** | <0.0001 | LN vs. HA | Yes | **** | <0.0001 |
| NCS 603 |  |  |  | NCS 603 |  |  |  |
| HA vs. HN | No | ns | 0.6091 | HA vs. HN | No | ns | 0.7918 |
| LN vs. HN | No | ns | 0.1929 | LN vs. HN | No | ns | 0.2216 |
| LN vs. HA | No | ns | 0.8433 | LN vs. HA | No | ns | 0.7193 |
| NCS 901 |  |  |  | NCS 901 |  |  |  |
| HA vs. HN | No | ns | 0.4534 | HA vs. HN | No | ns | 0.492 |
| LN vs. HN | No | ns | 0.9763 | LN vs. HN | No | ns | 0.232 |
| LN vs. HA | No | ns | 0.2498 | LN vs. HA | No | ns | 0.9532 |
| SELHI |  |  |  | SELHI |  |  |  |
| HA vs. HN | No | ns | 0.7622 | HA vs. HN | No | ns | 0.9 |
| LN vs. HN | No | ns | 0.984 | LN vs. HN | No | ns | >0.9999 |
| LN vs. HA | No | ns | 0.9241 | LN vs. HA | No | ns | 0.8953 |
| SUFALDHULA | |  |  | SUFALDHULA | |  |  |
| HA vs. HN | No | ns | 0.6038 | HA vs. HN | No | ns | 0.4514 |
| LN vs. HN | No | ns | 0.8627 | LN vs. HN | No | ns | 0.6592 |
| LN vs. HA | No | ns | 0.9657 | LN vs. HA | No | ns | 0.9855 |
| ZINYA KOLAMBA | |  |  | ZINYA KOLAMBA | |  |  |
| HA vs. HN | No | ns | 0.8508 | HA vs. HN | No | ns | 0.8286 |
| LN vs. HN | No | ns | 0.7233 | LN vs. HN | No | ns | 0.8711 |
| LN vs. HA | No | ns | 0.9949 | LN vs. HA | No | ns | 0.9997 |
| OR117-8 |  |  |  | OR117-8 |  |  |  |
| HA vs. HN | No | ns | 0.5244 | HA vs. HN | No | ns | 0.2685 |
| LN vs. HN | No | ns | >0.9999 | LN vs. HN | No | ns | 0.9605 |
| LN vs. HA | No | ns | 0.5377 | LN vs. HA | No | ns | 0.112 |
| ARC 10581 | |  |  | ARC 10581 | |  |  |
| HA vs. HN | No | ns | 0.9666 | HA vs. HN | No | ns | 0.8101 |
| LN vs. HN | No | ns | 0.8741 | LN vs. HN | No | ns | 0.7372 |
| LN vs. HA | No | ns | 0.9911 | LN vs. HA | No | ns | 0.9991 |
| SXC 216 |  |  |  | SXC 216 |  |  |  |
| HA vs. HN | No | ns | 0.861 | HA vs. HN | No | ns | 0.7987 |
| LN vs. HN | No | ns | 0.3448 | LN vs. HN | No | ns | 0.2593 |
| LN vs. HA | No | ns | 0.7971 | LN vs. HA | No | ns | 0.7658 |
| ARC 18325 | |  |  | ARC 18325 | |  |  |
| HA vs. HN | No | ns | 0.2516 | HA vs. HN | No | ns | 0.1545 |
| LN vs. HN | No | ns | 0.1864 | LN vs. HN | No | ns | 0.1222 |
| LN vs. HA | No | ns | 0.998 | LN vs. HA | No | ns | 0.9993 |
| MTU1010 |  |  |  | MTU1010 |  |  |  |
| HA vs. HN | No | ns | 0.3016 | HA vs. HN | No | ns | 0.2597 |
| LN vs. HN | No | ns | 0.1296 | LN vs. HN | No | ns | 0.1285 |
| LN vs. HA | No | ns | 0.9608 | LN vs. HA | No | ns | 0.9781 |
| LRL (cm) |  |  |  | LRSA (cm^2^) |  |  |  |
| Sidak's multiple comparisons test | Significant? | Summary | Adjusted P Value | Sidak's multiple comparisons test | Significant? | Summary | Adjusted P Value |
| ARC 10799 | |  |  | ARC 10799 | |  |  |
| HA vs. HN | No | ns | 0.9998 | HA vs. HN | No | ns | >0.9999 |
| LN vs. HN | No | ns | 0.9874 | LN vs. HN | No | ns | 0.9918 |
| LN vs. HA | No | ns | 0.9948 | LN vs. HA | No | ns | 0.9902 |
| BHAINSA MUNDARIYA | | |  | BHAINSA MUNDARIYA | | |  |
| HA vs. HN | No | ns | 0.3533 | HA vs. HN | No | ns | 0.3795 |
| LN vs. HN | No | ns | 0.5573 | LN vs. HN | No | ns | 0.6557 |
| LN vs. HA | No | ns | 0.9838 | LN vs. HA | No | ns | 0.965 |
| BHU BHUSI | |  |  | BHU BHUSI | |  |  |
| HA vs. HN | No | ns | 0.9887 | HA vs. HN | No | ns | 0.7377 |
| LN vs. HN | No | ns | 0.2789 | LN vs. HN | No | ns | 0.9721 |
| LN vs. HA | No | ns | 0.4398 | LN vs. HA | No | ns | 0.477 |
| DONGREM | |  |  | DONGREM | |  |  |
| HA vs. HN | No | ns | >0.9999 | HA vs. HN | No | ns | 0.9962 |
| LN vs. HN | No | ns | 0.9992 | LN vs. HN | No | ns | 0.9973 |
| LN vs. HA | No | ns | 0.9996 | LN vs. HA | No | ns | >0.9999 |
| LOCAL |  |  |  | LOCAL |  |  |  |
| HA vs. HN | No | ns | 0.689 | HA vs. HN | No | ns | 0.6918 |
| LN vs. HN | Yes | * | 0.0142 | LN vs. HN | Yes | ** | 0.0089 |
| LN vs. HA | No | ns | 0.1564 | LN vs. HA | No | ns | 0.108 |
| NCS 603 |  |  |  | NCS 603 |  |  |  |
| HA vs. HN | No | ns | 0.4145 | HA vs. HN | No | ns | 0.449 |
| LN vs. HN | No | ns | 0.1723 | LN vs. HN | No | ns | 0.1953 |
| LN vs. HA | No | ns | 0.9437 | LN vs. HA | No | ns | 0.9462 |
| NCS 901 |  |  |  | NCS 901 |  |  |  |
| HA vs. HN | No | ns | 0.4966 | HA vs. HN | No | ns | 0.5364 |
| LN vs. HN | No | ns | 0.1649 | LN vs. HN | No | ns | 0.0955 |
| LN vs. HA | Yes | ** | 0.0069 | LN vs. HA | Yes | ** | 0.0039 |
| SELHI |  |  |  | SELHI |  |  |  |
| HA vs. HN | No | ns | 0.5137 | HA vs. HN | No | ns | 0.4932 |
| LN vs. HN | No | ns | 0.7418 | LN vs. HN | No | ns | 0.7324 |
| LN vs. HA | No | ns | 0.981 | LN vs. HA | No | ns | 0.9783 |
| SUFALDHULA | |  |  | SUFALDHULA | |  |  |
| HA vs. HN | No | ns | 0.8319 | HA vs. HN | No | ns | 0.8431 |
| LN vs. HN | No | ns | 0.9888 | LN vs. HN | No | ns | >0.9999 |
| LN vs. HA | No | ns | 0.9509 | LN vs. HA | No | ns | 0.8714 |
| ZINYA KOLAMBA | |  |  | ZINYA KOLAMBA | |  |  |
| HA vs. HN | No | ns | 0.9354 | HA vs. HN | No | ns | 0.9331 |
| LN vs. HN | No | ns | 0.4712 | LN vs. HN | No | ns | 0.6036 |
| LN vs. HA | No | ns | 0.8147 | LN vs. HA | No | ns | 0.9134 |
| OR117-8 |  |  |  | OR117-8 |  |  |  |
| HA vs. HN | No | ns | 0.8465 | HA vs. HN | No | ns | 0.8444 |
| LN vs. HN | No | ns | 0.9494 | LN vs. HN | No | ns | 0.9 |
| LN vs. HA | No | ns | 0.9922 | LN vs. HA | No | ns | 0.9992 |
| ARC 10581 | |  |  | ARC 10581 | |  |  |
| HA vs. HN | No | ns | 0.9956 | HA vs. HN | No | ns | >0.9999 |
| LN vs. HN | No | ns | 0.9829 | LN vs. HN | No | ns | 0.9755 |
| LN vs. HA | No | ns | 0.9325 | LN vs. HA | No | ns | 0.9732 |
| SXC 216 |  |  |  | SXC 216 |  |  |  |
| HA vs. HN | No | ns | 0.9425 | HA vs. HN | No | ns | 0.9571 |
| LN vs. HN | No | ns | 0.6609 | LN vs. HN | No | ns | 0.7823 |
| LN vs. HA | No | ns | 0.9338 | LN vs. HA | No | ns | 0.9714 |
| ARC 18325 | |  |  | ARC 18325 | |  |  |
| HA vs. HN | No | ns | 0.5835 | HA vs. HN | No | ns | 0.5673 |
| LN vs. HN | No | ns | 0.4486 | LN vs. HN | No | ns | 0.6207 |
| LN vs. HA | No | ns | 0.9958 | LN vs. HA | No | ns | 0.9998 |
| MTU1010 |  |  |  | MTU1010 |  |  |  |
| HA vs. HN | No | ns | 0.4078 | HA vs. HN | No | ns | 0.432 |
| LN vs. HN | No | ns | 0.1851 | LN vs. HN | No | ns | 0.2276 |
| LN vs. HA | No | ns | 0.9572 | LN vs. HA | No | ns | 0.9731 |
| LRV (cm^3^) |  |  |  | MRL (cm) |  |  |  |
| Sidak's multiple comparisons test | Significant? | Summary | Adjusted P Value | Sidak's multiple comparisons test | Significant? | Summary | Adjusted P Value |
| ARC 10799 | |  |  | ARC 10799 | |  |  |
| HA vs. HN | No | ns | 0.9999 | HA vs. HN | No | ns | 0.9188 |
| LN vs. HN | No | ns | 0.9933 | LN vs. HN | No | ns | 0.9992 |
| LN vs. HA | No | ns | 0.9872 | LN vs. HA | No | ns | 0.868 |
| BHAINSA MUNDARIYA | | |  | BHAINSA MUNDARIYA | | |  |
| HA vs. HN | No | ns | 0.3812 | HA vs. HN | No | ns | 0.9996 |
| LN vs. HN | No | ns | 0.6718 | LN vs. HN | No | ns | 0.9996 |
| LN vs. HA | No | ns | 0.9599 | LN vs. HA | No | ns | >0.9999 |
| BHU BHUSI | |  |  | BHU BHUSI | |  |  |
| HA vs. HN | No | ns | 0.879 | HA vs. HN | No | ns | 0.5815 |
| LN vs. HN | No | ns | 0.1395 | LN vs. HN | No | ns | 0.9441 |
| LN vs. HA | No | ns | 0.454 | LN vs. HA | No | ns | 0.8853 |
| DONGREM | |  |  | DONGREM | |  |  |
| HA vs. HN | No | ns | 0.9876 | HA vs. HN | No | ns | 0.9969 |
| LN vs. HN | No | ns | 0.9935 | LN vs. HN | No | ns | 0.9915 |
| LN vs. HA | No | ns | 0.9999 | LN vs. HA | No | ns | 0.9604 |
| LOCAL |  |  |  | LOCAL |  |  |  |
| HA vs. HN | No | ns | 0.7117 | HA vs. HN | No | ns | 0.0796 |
| LN vs. HN | Yes | ** | 0.0081 | LN vs. HN | No | ns | 0.3929 |
| LN vs. HA | No | ns | 0.0936 | LN vs. HA | Yes | ** | 0.0016 |
| NCS 603 |  |  |  | NCS 603 |  |  |  |
| HA vs. HN | No | ns | 0.465 | HA vs. HN | No | ns | 0.8372 |
| LN vs. HN | No | ns | 0.2145 | LN vs. HN | No | ns | 0.5682 |
| LN vs. HA | No | ns | 0.9532 | LN vs. HA | No | ns | 0.9651 |
| NCS 901 |  |  |  | NCS 901 |  |  |  |
| HA vs. HN | No | ns | 0.5578 | HA vs. HN | No | ns | 0.8829 |
| LN vs. HN | Yes | * | 0.0487 | LN vs. HN | No | ns | 0.3466 |
| LN vs. HA | Yes | ** | 0.0018 | LN vs. HA | No | ns | 0.0965 |
| SELHI |  |  |  | SELHI |  |  |  |
| HA vs. HN | No | ns | 0.4846 | HA vs. HN | No | ns | 0.9349 |
| LN vs. HN | No | ns | 0.7199 | LN vs. HN | No | ns | >0.9999 |
| LN vs. HA | No | ns | 0.9794 | LN vs. HA | No | ns | 0.9146 |
| SUFALDHULA | |  |  | SUFALDHULA | |  |  |
| HA vs. HN | No | ns | 0.8492 | HA vs. HN | No | ns | 0.7322 |
| LN vs. HN | No | ns | 0.9996 | LN vs. HN | No | ns | 0.9526 |
| LN vs. HA | No | ns | 0.7986 | LN vs. HA | No | ns | 0.9562 |
| ZINYA KOLAMBA | |  |  | ZINYA KOLAMBA | |  |  |
| HA vs. HN | No | ns | 0.9495 | HA vs. HN | No | ns | 0.9351 |
| LN vs. HN | No | ns | 0.7352 | LN vs. HN | No | ns | 0.9453 |
| LN vs. HA | No | ns | 0.9603 | LN vs. HA | No | ns | >0.9999 |
| OR117-8 |  |  |  | OR117-8 |  |  |  |
| HA vs. HN | No | ns | 0.8392 | HA vs. HN | No | ns | 0.6806 |
| LN vs. HN | No | ns | 0.8465 | LN vs. HN | No | ns | 0.9983 |
| LN vs. HA | No | ns | >0.9999 | LN vs. HA | No | ns | 0.7762 |
| ARC 10581 | |  |  | ARC 10581 | |  |  |
| HA vs. HN | No | ns | 0.9997 | HA vs. HN | No | ns | 0.9775 |
| LN vs. HN | No | ns | 0.9734 | LN vs. HN | No | ns | 0.931 |
| LN vs. HA | No | ns | 0.9865 | LN vs. HA | No | ns | 0.9972 |
| SXC 216 |  |  |  | SXC 216 |  |  |  |
| HA vs. HN | No | ns | 0.9699 | HA vs. HN | No | ns | 0.9475 |
| LN vs. HN | No | ns | 0.8029 | LN vs. HN | No | ns | 0.3683 |
| LN vs. HA | No | ns | 0.9674 | LN vs. HA | No | ns | 0.6869 |
| ARC 18325 | |  |  | ARC 18325 | |  |  |
| HA vs. HN | No | ns | 0.5599 | HA vs. HN | No | ns | 0.2663 |
| LN vs. HN | No | ns | 0.7374 | LN vs. HN | No | ns | 0.21 |
| LN vs. HA | No | ns | 0.9909 | LN vs. HA | No | ns | 0.9989 |
| MTU1010 |  |  |  | MTU1010 |  |  |  |
| HA vs. HN | No | ns | 0.4918 | HA vs. HN | No | ns | 0.6716 |
| LN vs. HN | No | ns | 0.2926 | LN vs. HN | No | ns | 0.3666 |
| LN vs. HA | No | ns | 0.9814 | LN vs. HA | No | ns | 0.9533 |
| MRSA (cm^2^) |  |  |  | MRV (cm^3^) |  |  |  |
| Sidak's multiple comparisons test | Significant? | Summary | Adjusted P Value | Sidak's multiple comparisons test | Significant? | Summary | Adjusted P Value |
| ARC 10799 | |  |  | ARC 10799 | |  |  |
| HA vs. HN | No | ns | 0.8076 | HA vs. HN | No | ns | 0.7211 |
| LN vs. HN | No | ns | 0.995 | LN vs. HN | No | ns | 0.9956 |
| LN vs. HA | No | ns | 0.6717 | LN vs. HA | No | ns | 0.582 |
| BHAINSA MUNDARIYA | | |  | BHAINSA MUNDARIYA | | |  |
| HA vs. HN | No | ns | 0.9997 | HA vs. HN | Yes | **** | <0.0001 |
| LN vs. HN | No | ns | 0.9378 | LN vs. HN | Yes | **** | <0.0001 |
| LN vs. HA | No | ns | 0.9076 | LN vs. HA | Yes | * | 0.0186 |
| BHU BHUSI | |  |  | BHU BHUSI | |  |  |
| HA vs. HN | No | ns | 0.4903 | HA vs. HN | No | ns | 0.3427 |
| LN vs. HN | No | ns | 0.9704 | LN vs. HN | Yes | * | 0.0326 |
| LN vs. HA | No | ns | 0.7555 | LN vs. HA | No | ns | 0.614 |
| DONGREM | |  |  | DONGREM | |  |  |
| HA vs. HN | No | ns | >0.9999 | HA vs. HN | No | ns | 0.995 |
| LN vs. HN | No | ns | 0.9993 | LN vs. HN | No | ns | 0.999 |
| LN vs. HA | No | ns | 0.9988 | LN vs. HA | No | ns | 0.9811 |
| LOCAL |  |  |  | LOCAL |  |  |  |
| HA vs. HN | Yes | * | 0.0291 | HA vs. HN | No | ns | 0.2734 |
| LN vs. HN | Yes | * | 0.0197 | LN vs. HN | Yes | **** | <0.0001 |
| LN vs. HA | Yes | **** | <0.0001 | LN vs. HA | Yes | **** | <0.0001 |
| NCS 603 |  |  |  | NCS 603 |  |  |  |
| HA vs. HN | No | ns | 0.7917 | HA vs. HN | No | ns | 0.4319 |
| LN vs. HN | No | ns | 0.3182 | LN vs. HN | No | ns | 0.1005 |
| LN vs. HA | No | ns | 0.8406 | LN vs. HA | No | ns | 0.8187 |
| NCS 901 |  |  |  | NCS 901 |  |  |  |
| HA vs. HN | No | ns | 0.6112 | HA vs. HN | No | ns | 0.0809 |
| LN vs. HN | No | ns | 0.9791 | LN vs. HN | Yes | **** | <0.0001 |
| LN vs. HA | No | ns | 0.8337 | LN vs. HA | Yes | **** | <0.0001 |
| SELHI |  |  |  | SELHI |  |  |  |
| HA vs. HN | No | ns | 0.9431 | HA vs. HN | No | ns | 0.9531 |
| LN vs. HN | No | ns | 0.999 | LN vs. HN | No | ns | 0.9753 |
| LN vs. HA | No | ns | 0.896 | LN vs. HA | No | ns | 0.9996 |
| SUFALDHULA | |  |  | SUFALDHULA | |  |  |
| HA vs. HN | No | ns | 0.5041 | HA vs. HN | No | ns | 0.3834 |
| LN vs. HN | No | ns | 0.662 | LN vs. HN | Yes | ** | 0.0049 |
| LN vs. HA | No | ns | 0.9937 | LN vs. HA | Yes | **** | <0.0001 |
| ZINYA KOLAMBA | |  |  | ZINYA KOLAMBA | |  |  |
| HA vs. HN | No | ns | 0.8052 | HA vs. HN | No | ns | 0.858 |
| LN vs. HN | No | ns | 0.8698 | LN vs. HN | No | ns | 0.9177 |
| LN vs. HA | No | ns | 0.9991 | LN vs. HA | No | ns | 0.9988 |
| OR117-8 |  |  |  | OR117-8 |  |  |  |
| HA vs. HN | No | ns | 0.4944 | HA vs. HN | No | ns | 0.6801 |
| LN vs. HN | No | ns | 0.9875 | LN vs. HN | No | ns | 0.7349 |
| LN vs. HA | No | ns | 0.3172 | LN vs. HA | No | ns | 0.1648 |
| ARC 10581 | |  |  | ARC 10581 | |  |  |
| HA vs. HN | No | ns | 0.9256 | HA vs. HN | No | ns | 0.9048 |
| LN vs. HN | No | ns | 0.86 | LN vs. HN | No | ns | 0.8569 |
| LN vs. HA | No | ns | 0.9982 | LN vs. HA | No | ns | 0.9994 |
| SXC 216 |  |  |  | SXC 216 |  |  |  |
| HA vs. HN | No | ns | 0.82 | HA vs. HN | No | ns | 0.9037 |
| LN vs. HN | No | ns | 0.2152 | LN vs. HN | No | ns | 0.3927 |
| LN vs. HA | No | ns | 0.6766 | LN vs. HA | No | ns | 0.791 |
| ARC 18325 | |  |  | ARC 18325 | |  |  |
| HA vs. HN | No | ns | 0.2078 | HA vs. HN | No | ns | 0.5266 |
| LN vs. HN | No | ns | 0.1243 | LN vs. HN | No | ns | 0.2871 |
| LN vs. HA | No | ns | 0.992 | LN vs. HA | No | ns | 0.9696 |
| MTU1010 |  |  |  | MTU1010 |  |  |  |
| HA vs. HN | No | ns | 0.4149 | HA vs. HN | No | ns | 0.1103 |
| LN vs. HN | No | ns | 0.2043 | LN vs. HN | No | ns | 0.0837 |
| LN vs. HA | No | ns | 0.9668 | LN vs. HA | No | ns | 0.999 |
| No. of forks |  |  |  | No. of tips |  |  |  |
| Sidak's multiple comparisons test | Significant? | Summary | Adjusted P Value | Sidak's multiple comparisons test | Significant? | Summary | Adjusted P Value |
| ARC 10799 |  |  |  | ARC 10799 | |  |  |
| HA vs. HN | No | ns | 0.8893 | HA vs. HN | No | ns | 0.9934 |
| LN vs. HN | No | ns | 0.9909 | LN vs. HN | No | ns | 0.9991 |
| LN vs. HA | No | ns | 0.9741 | LN vs. HA | No | ns | 0.9992 |
| BHAINSA MUNDARIYA |  |  |  | BHAINSA MUNDARIYA | | |  |
| HA vs. HN | No | ns | 0.5374 | HA vs. HN | No | ns | 0.6875 |
| LN vs. HN | No | ns | 0.2813 | LN vs. HN | No | ns | 0.4153 |
| LN vs. HA | No | ns | 0.9633 | LN vs. HA | No | ns | 0.9678 |
| BHU BHUSI |  |  |  | BHU BHUSI | |  |  |
| HA vs. HN | No | ns | 0.7623 | HA vs. HN | No | ns | 0.3757 |
| LN vs. HN | No | ns | 0.1249 | LN vs. HN | No | ns | 0.6622 |
| LN vs. HA | No | ns | 0.5602 | LN vs. HA | Yes | * | 0.0451 |
| DONGREM |  |  |  | DONGREM | |  |  |
| HA vs. HN | No | ns | 0.9998 | HA vs. HN | No | ns | >0.9999 |
| LN vs. HN | No | ns | >0.9999 | LN vs. HN | No | ns | >0.9999 |
| LN vs. HA | No | ns | >0.9999 | LN vs. HA | No | ns | >0.9999 |
| LOCAL |  |  |  | LOCAL |  |  |  |
| HA vs. HN | No | ns | 0.9922 | HA vs. HN | No | ns | 0.3186 |
| LN vs. HN | Yes | * | 0.0133 | LN vs. HN | Yes | ** | 0.0033 |
| LN vs. HA | Yes | ** | 0.0066 | LN vs. HA | No | ns | 0.1798 |
| NCS 603 |  |  |  | NCS 603 |  |  |  |
| HA vs. HN | No | ns | 0.4785 | HA vs. HN | No | ns | 0.0941 |
| LN vs. HN | No | ns | 0.1646 | LN vs. HN | Yes | * | 0.0133 |
| LN vs. HA | No | ns | 0.898 | LN vs. HA | No | ns | 0.8197 |
| NCS 901 |  |  |  | NCS 901 |  |  |  |
| HA vs. HN | No | ns | 0.2683 | HA vs. HN | No | ns | 0.625 |
| LN vs. HN | No | ns | 0.8632 | LN vs. HN | No | ns | 0.0627 |
| LN vs. HA | No | ns | 0.062 | LN vs. HA | Yes | ** | 0.0034 |
| SELHI |  |  |  | SELHI |  |  |  |
| HA vs. HN | No | ns | 0.7969 | HA vs. HN | No | ns | 0.2989 |
| LN vs. HN | No | ns | 0.9973 | LN vs. HN | No | ns | 0.322 |
| LN vs. HA | No | ns | 0.8882 | LN vs. HA | No | ns | >0.9999 |
| SUFALDHULA |  |  |  | SUFALDHULA | |  |  |
| HA vs. HN | No | ns | 0.4292 | HA vs. HN | No | ns | 0.8081 |
| LN vs. HN | No | ns | 0.7194 | LN vs. HN | No | ns | 0.888 |
| LN vs. HA | No | ns | 0.9616 | LN vs. HA | No | ns | 0.9981 |
| ZINYA KOLAMBA |  |  |  | ZINYA KOLAMBA | |  |  |
| HA vs. HN | No | ns | 0.9619 | HA vs. HN | No | ns | 0.9638 |
| LN vs. HN | No | ns | 0.662 | LN vs. HN | No | ns | 0.0611 |
| LN vs. HA | No | ns | 0.9086 | LN vs. HA | No | ns | 0.1574 |
| OR117-8 |  |  |  | OR117-8 |  |  |  |
| HA vs. HN | No | ns | 0.7721 | HA vs. HN | No | ns | 0.7376 |
| LN vs. HN | No | ns | 0.9781 | LN vs. HN | No | ns | 0.9895 |
| LN vs. HA | No | ns | 0.5348 | LN vs. HA | No | ns | 0.8908 |
| ARC 10581 |  |  |  | ARC 10581 | |  |  |
| HA vs. HN | No | ns | 0.9999 | HA vs. HN | No | ns | 0.8485 |
| LN vs. HN | No | ns | 0.9669 | LN vs. HN | No | ns | 0.9949 |
| LN vs. HA | No | ns | 0.9518 | LN vs. HA | No | ns | 0.7204 |
| SXC 216 |  |  |  | SXC 216 |  |  |  |
| HA vs. HN | No | ns | 0.453 | HA vs. HN | No | ns | >0.9999 |
| LN vs. HN | No | ns | 0.9548 | LN vs. HN | No | ns | 0.9786 |
| LN vs. HA | No | ns | 0.7599 | LN vs. HA | No | ns | 0.9822 |
| ARC 18325 |  |  |  | ARC 18325 | |  |  |
| HA vs. HN | No | ns | 0.9958 | HA vs. HN | No | ns | 0.5684 |
| LN vs. HN | No | ns | 0.2462 | LN vs. HN | No | ns | 0.2855 |
| LN vs. HA | No | ns | 0.3492 | LN vs. HA | Yes | * | 0.0207 |
| MTU1010 |  |  |  | MTU1010 |  |  |  |
| HA vs. HN | No | ns | 0.4953 | HA vs. HN | No | ns | 0.5503 |
| LN vs. HN | No | ns | 0.107 | LN vs. HN | No | ns | 0.1285 |
| LN vs. HA | No | ns | 0.7779 | LN vs. HA | No | ns | 0.7795 |

**Supplementary table 6. Sidak’s multiple comparison test on the effect of high nitrate (HN), high ammonium (HA) and nitrogen deficient (LN) conditions on nitrate reductase activity in shoots (NR), nitrate reductase activity in roots, shoot protein content, root protein content, glutamine synthetase in shoots (GS), glutamine synthetase in roots, glutamate synthase activity in shoots (GOGAT), glutamate synthase activity of rice seedlings, glutamate dehydrogenase in shoots (GDH), glutamate dehydrogenase in roots grown under hydroponic culture**

| NR – leaf (µmol NO_2_^-^ formed g^-1^ FW h^-1^) |  |  |  | NR – root  (µmol NO_2_^-^ formed g^-1^ FW h^-1^) |  |  |  |
| --- | --- | --- | --- | --- | --- | --- | --- |
| Sidak's multiple comparisons test | Significant? | Summary | Adjusted P Value | Sidak's multiple comparisons test | Significant? | Summary | Adjusted P Value |
| ARC 10799 | |  |  | ARC 10799 | |  |  |
| HA vs. HN | Yes | **** | <0.0001 | HA vs. HN | No | ns | 0.1194 |
| LN vs. HN | Yes | **** | <0.0001 | LN vs. HN | Yes | ** | 0.0039 |
| LN vs. HA | No | ns | 0.0646 | LN vs. HA | Yes | **** | <0.0001 |
| OR117-8 |  |  |  | OR117-8 |  |  |  |
| HA vs. HN | Yes | * | 0.0167 | HA vs. HN | No | ns | 0.4554 |
| LN vs. HN | No | ns | 0.9975 | LN vs. HN | Yes | **** | <0.0001 |
| LN vs. HA | Yes | * | 0.0258 | LN vs. HA | Yes | **** | <0.0001 |
| ARC 10581 | |  |  | ARC 10581 | |  |  |
| HA vs. HN | Yes | **** | <0.0001 | HA vs. HN | Yes | **** | <0.0001 |
| LN vs. HN | Yes | **** | <0.0001 | LN vs. HN | Yes | *** | 0.0005 |
| LN vs. HA | No | ns | 0.7151 | LN vs. HA | Yes | **** | <0.0001 |
| SXC 216 |  |  |  | SXC 216 |  |  |  |
| HA vs. HN | Yes | **** | <0.0001 | HA vs. HN | Yes | ** | 0.002 |
| LN vs. HN | Yes | **** | <0.0001 | LN vs. HN | Yes | **** | <0.0001 |
| LN vs. HA | Yes | **** | <0.0001 | LN vs. HA | Yes | **** | <0.0001 |
| ARC 18325 | |  |  | ARC 18325 | |  |  |
| HA vs. HN | Yes | **** | <0.0001 | HA vs. HN | Yes | **** | <0.0001 |
| LN vs. HN | Yes | **** | <0.0001 | LN vs. HN | Yes | **** | <0.0001 |
| LN vs. HA | Yes | **** | <0.0001 | LN vs. HA | No | ns | 0.1176 |
| MTU1010 |  |  |  | MTU1010 |  |  |  |
| HA vs. HN | Yes | * | 0.0176 | HA vs. HN | Yes | **** | <0.0001 |
| LN vs. HN | Yes | **** | <0.0001 | LN vs. HN | Yes | **** | <0.0001 |
| LN vs. HA | Yes | *** | 0.0004 | LN vs. HA | Yes | **** | <0.0001 |
| GS – shoot  (µmol γ glutamyl hydroxamate formed g^-1^ FW h^-1^) |  |  |  | GS - root  (µmol γ glutamyl hydroxamate formed g^-1^ FW h^-1^) |  |  |  |
| Sidak's multiple comparisons test | Significant? | Summary | Adjusted P Value | Sidak's multiple comparisons test | Significant? | Summary | Adjusted P Value |
| ARC 10799 | |  |  | ARC 10799 | |  |  |
| HA vs. HN | No | ns | 0.9998 | HA vs. HN | No | ns | 0.8245 |
| LN vs. HN | No | ns | 0.9012 | LN vs. HN | No | ns | 0.1867 |
| LN vs. HA | No | ns | 0.9314 | LN vs. HA | No | ns | 0.617 |
| OR117-8 |  |  |  | OR117-8 |  |  |  |
| HA vs. HN | Yes | * | 0.0162 | HA vs. HN | No | ns | 0.4816 |
| LN vs. HN | No | ns | >0.9999 | LN vs. HN | No | ns | 0.3898 |
| LN vs. HA | Yes | * | 0.0146 | LN vs. HA | No | ns | 0.9984 |
| ARC 10581 | |  |  | ARC 10581 | |  |  |
| HA vs. HN | No | ns | 0.9594 | HA vs. HN | Yes | **** | <0.0001 |
| LN vs. HN | Yes | **** | <0.0001 | LN vs. HN | No | ns | 0.3402 |
| LN vs. HA | Yes | **** | <0.0001 | LN vs. HA | Yes | **** | <0.0001 |
| SXC 216 |  |  |  | SXC 216 |  |  |  |
| HA vs. HN | Yes | **** | <0.0001 | HA vs. HN | No | ns | 0.9185 |
| LN vs. HN | Yes | **** | <0.0001 | LN vs. HN | Yes | **** | <0.0001 |
| LN vs. HA | No | ns | 0.893 | LN vs. HA | Yes | **** | <0.0001 |
| ARC 18325 | |  |  | ARC 18325 | |  |  |
| HA vs. HN | Yes | **** | <0.0001 | HA vs. HN | Yes | **** | <0.0001 |
| LN vs. HN | No | ns | 0.6213 | LN vs. HN | Yes | **** | <0.0001 |
| LN vs. HA | Yes | **** | <0.0001 | LN vs. HA | No | ns | 0.1879 |
| MTU1010 |  |  |  | MTU1010 |  |  |  |
| HA vs. HN | No | ns | 0.5309 | HA vs. HN | No | ns | 0.8979 |
| LN vs. HN | No | ns | 0.1214 | LN vs. HN | Yes | **** | <0.0001 |
| LN vs. HA | No | ns | 0.7754 | LN vs. HA | Yes | **** | <0.0001 |
| GOGAT – Shoot  ( µmol NADH oxidized mg^-1^ FW h^-1^) | |  |  | GOGAT – Root  ( µmol NADH oxidized mg^-1^ FW h^-1^) | |  |  |
| Sidak's multiple comparisons test | Significant? | Summary | Adjusted P Value | Sidak's multiple comparisons test | Significant? | Summary | Adjusted P Value |
| ARC 10799 | |  |  | ARC 10799 | |  |  |
| HA vs. HN | No | ns | 0.9572 | HA vs. HN | No | ns | 0.9487 |
| LN vs. HN | Yes | **** | <0.0001 | LN vs. HN | No | ns | 0.9861 |
| LN vs. HA | Yes | **** | <0.0001 | LN vs. HA | No | ns | 0.8159 |
| OR117-8 |  |  |  | OR117-8 |  |  |  |
| HA vs. HN | No | ns | 0.9983 | HA vs. HN | No | ns | 0.8662 |
| LN vs. HN | Yes | **** | <0.0001 | LN vs. HN | No | ns | 0.3823 |
| LN vs. HA | Yes | **** | <0.0001 | LN vs. HA | No | ns | 0.8264 |
| ARC 10581 | |  |  | ARC 10581 | |  |  |
| HA vs. HN | No | ns | 0.9929 | HA vs. HN | Yes | **** | <0.0001 |
| LN vs. HN | Yes | **** | <0.0001 | LN vs. HN | Yes | **** | <0.0001 |
| LN vs. HA | Yes | **** | <0.0001 | LN vs. HA | Yes | **** | <0.0001 |
| SXC 216 |  |  |  | SXC 216 |  |  |  |
| HA vs. HN | No | ns | 0.125 | HA vs. HN | No | ns | 0.9759 |
| LN vs. HN | No | ns | 0.8385 | LN vs. HN | Yes | **** | <0.0001 |
| LN vs. HA | No | ns | 0.4672 | LN vs. HA | Yes | **** | <0.0001 |
| ARC 18325 | |  |  | ARC 18325 | |  |  |
| HA vs. HN | No | ns | >0.9999 | HA vs. HN | No | ns | 0.0797 |
| LN vs. HN | No | ns | 0.3194 | LN vs. HN | Yes | ** | 0.0054 |
| LN vs. HA | No | ns | 0.2933 | LN vs. HA | No | ns | 0.6447 |
| MTU1010 |  |  |  | MTU1010 |  |  |  |
| HA vs. HN | No | ns | 0.233 | HA vs. HN | No | ns | 0.997 |
| LN vs. HN | No | ns | 0.3904 | LN vs. HN | Yes | ** | 0.0019 |
| LN vs. HA | Yes | ** | 0.0078 | LN vs. HA | Yes | ** | 0.0032 |
| GDH -shoot (µmol NADH oxidized mg^-1^ FW h^-1^) | |  |  | GDH -root (µmol NADH oxidized mg^-1^ FW h^-1^) | |  |  |
| Sidak's multiple comparisons test | Significant? | Summary | Adjusted P Value | Sidak's multiple comparisons test | Significant? | Summary | Adjusted P Value |
| ARC 10799 | |  |  | ARC 10799 | |  |  |
| HA vs. HN | Yes | **** | <0.0001 | HA vs. HN | No | ns | 0.3124 |
| LN vs. HN | Yes | *** | 0.0003 | LN vs. HN | No | ns | >0.9999 |
| LN vs. HA | Yes | **** | <0.0001 | LN vs. HA | No | ns | 0.317 |
| OR117-8 |  |  |  | OR117-8 |  |  |  |
| HA vs. HN | No | ns | 0.964 | HA vs. HN | No | ns | 0.9998 |
| LN vs. HN | No | ns | 0.2182 | LN vs. HN | No | ns | 0.9928 |
| LN vs. HA | No | ns | 0.098 | LN vs. HA | No | ns | 0.9801 |
| ARC 10581 | |  |  | ARC 10581 | |  |  |
| HA vs. HN | Yes | ** | 0.0041 | HA vs. HN | No | ns | >0.9999 |
| LN vs. HN | No | ns | 0.8199 | LN vs. HN | Yes | **** | <0.0001 |
| LN vs. HA | Yes | * | 0.0239 | LN vs. HA | Yes | **** | <0.0001 |
| SXC 216 |  |  |  | SXC 216 |  |  |  |
| HA vs. HN | No | ns | 0.7559 | HA vs. HN | No | ns | 0.7569 |
| LN vs. HN | No | ns | 0.9715 | LN vs. HN | No | ns | 0.6759 |
| LN vs. HA | No | ns | 0.9433 | LN vs. HA | No | ns | 0.999 |
| ARC 18325 | |  |  | ARC 18325 | |  |  |
| HA vs. HN | No | ns | 0.9971 | HA vs. HN | No | ns | 0.4507 |
| LN vs. HN | Yes | * | 0.0216 | LN vs. HN | No | ns | 0.9769 |
| LN vs. HA | Yes | * | 0.032 | LN vs. HA | No | ns | 0.2404 |
| MTU1010 |  |  |  | MTU1010 |  |  |  |
| HA vs. HN | No | ns | 0.1312 | HA vs. HN | No | ns | >0.9999 |
| LN vs. HN | No | ns | 0.0628 | LN vs. HN | No | ns | >0.9999 |
| LN vs. HA | No | ns | 0.9762 | LN vs. HA | No | ns | >0.9999 |
| Protein – Shoot (mg g-1 FW) | |  |  | Protein - Root (mg g-1 FW) | |  |  |
| Sidak's multiple comparisons test | Significant? | Summary | Adjusted P Value | Sidak's multiple comparisons test | Significant? | Summary | Adjusted P Value |
| ARC 10799 | |  |  | ARC 10799 | |  |  |
| HA vs. HN | Yes | * | 0.015 | HA vs. HN | No | ns | 0.2119 |
| LN vs. HN | Yes | ** | 0.0015 | LN vs. HN | No | ns | 0.0572 |
| LN vs. HA | No | ns | 0.7968 | LN vs. HA | No | ns | 0.9014 |
| OR117-8 |  |  |  | OR117-8 |  |  |  |
| HA vs. HN | No | ns | 0.1713 | HA vs. HN | Yes | ** | 0.0023 |
| LN vs. HN | Yes | **** | <0.0001 | LN vs. HN | Yes | ** | 0.001 |
| LN vs. HA | Yes | **** | <0.0001 | LN vs. HA | No | ns | 0.9894 |
| ARC 10581 | |  |  | ARC 10581 | |  |  |
| HA vs. HN | Yes | * | 0.038 | HA vs. HN | Yes | ** | 0.0084 |
| LN vs. HN | Yes | **** | <0.0001 | LN vs. HN | No | ns | 0.9986 |
| LN vs. HA | Yes | **** | <0.0001 | LN vs. HA | Yes | * | 0.0122 |
| SXC 216 |  |  |  | SXC 216 |  |  |  |
| HA vs. HN | No | ns | 0.5096 | HA vs. HN | No | ns | 0.1554 |
| LN vs. HN | Yes | **** | <0.0001 | LN vs. HN | Yes | **** | <0.0001 |
| LN vs. HA | Yes | ** | 0.0012 | LN vs. HA | Yes | **** | <0.0001 |
| ARC 18325 | |  |  | ARC 18325 | |  |  |
| HA vs. HN | No | ns | 0.9993 | HA vs. HN | Yes | **** | <0.0001 |
| LN vs. HN | Yes | **** | <0.0001 | LN vs. HN | Yes | **** | <0.0001 |
| LN vs. HA | Yes | **** | <0.0001 | LN vs. HA | No | ns | 0.9991 |
| MTU1010 |  |  |  | MTU1010 |  |  |  |
| HA vs. HN | No | ns | 0.7526 | HA vs. HN | No | ns | 0.2823 |
| LN vs. HN | Yes | *** | 0.0001 | LN vs. HN | Yes | **** | <0.0001 |
| LN vs. HA | Yes | **** | <0.0001 | LN vs. HA | Yes | **** | <0.0001 |
