## Supplementary material for "Association of haplotypes of AAP family amino acid transporters with nitrogen response and nitrogen use efficiency in rice grown under hydroponics and field conditions": Table 1

**Table 1. Details of the effects of the non- synonymous SNPs and 5’ and 3’ UTR variants retrieved from IRRI SNP-Seek Database (https://snpseek.irri.org/) and RiceVarMap (**[**https://ricevarmap.ncpgr.cn/**](https://ricevarmap.ncpgr.cn/)**)**

| **Gene** | **Variant ID** | **Varaition type** | **Locus (Chromosome:Position)** | **Variation** | **snpEFF Annotation and effect** | **PolyPhen2-Effect and Score** | **SIFT Effect and SIFT Score** |
| --- | --- | --- | --- | --- | --- | --- | --- |
| OsAAP3 | vg0621191370 | Non-synonymous SNP | LOC Os06g36180.1 (chr06:21191370) | T🡪C | Missense variant; Val379Ala; Moderate | Benign;  -0.418 | Tolerated; 0.51 |
| OsAAP5 | vg 0138118647 | Non-synonymous SNP | LOC Os01g65660.1  (chr01:38118647) | T🡪G | Missense variant; Glu350Ala; Moderate | Possible damaging;  -1.815 | Tolerated; 0.13 |
| OsAAP11 | Vg1104799528 | Non-synonymous SNP | LOC Os11g09020 (chr11: 4799528) | T🡪C | Missense variant; Met36Thr; Moderate | Probably damaging;  -2.408 | Tolerated; 1 |
| OsAAP11 | Vg1104799537 | Non-synonymous SNP | LOC Os11g09020 (chr11: 4799537) | T🡪C | Missense variant; Val39Ala; Moderate | Benign; 0.435 | Tolerated; 0.10 |
| OsAAP11 | Vg1104799770 | Non-synonymous SNP | LOC Os11g09020 (chr11: 4799770) | T🡪G | Missense variant; Ser117Ala; Moderate | Benign; -0.413 | Tolerated 0.52 |
| OsAAP11 | Vg1104800214 | Non-synonymous SNP | LOC Os11g09020 (chr11: 48000214) | A🡪G | Missensee variant; Thr233Ala;  Moderate | Benign; -1.408 | Tolerated 0.38 |
| OsAAP11 | Vg1104800544 | Non-synonymous SNP | LOC Os11g09020 (chr11: 48000544) | A🡪G | Missense variant; Lys308Arg; Moderate | Benign; 0.304 | Tolerated  0.54 |
| OsAAP11 | Vg1104800985 | Non-synonymous SNP | LOC Os11g09020 (chr11: 48000214) | C🡪T | Missense variant; Ala455Val; Moderate | Benign; 0.593 | Tolerated 0.40 |
